# From the fly connectome to exact ring attractor dynamics

**DOI:** 10.1101/2024.11.01.621596

**Authors:** Tirthabir Biswas, Angel Stanoev, Sandro Romani, James E. Fitzgerald

**Author notes:** These authors contributed equally to this work.

## Abstract

A cognitive compass enabling spatial navigation requires the neural representation of head direction (HD), yet the neural circuit architecture enabling this representation remains unclear. While various network models have been proposed to explain HD systems, these models rely on simplified circuit architectures that may be irreconcilable with empirical observations from connectomes. Here we construct a neural network model for the fruit fly HD system that satisfies both connectome-derived architectural constraints and the functional requirement of continuous head-direction representation. To achieve this, we first characterized an ensemble of continuous attractor networks where compass neurons providing local mutual excitation are coupled to inhibitory neurons. Our multipopulation model allowed us to discover a new class of ring attractor network with weaker symmetry requirements than usually assumed. For each of the four available fly connectomes, our analyses uncovered three distinct realizations of these networks. Furthermore, we found that synaptic variations among and around the fly connectomes can be compensated by cell-type-specific rescaling of synaptic weights, which could be potentially achieved through neuromodulation. The fly connectomes are ideally configured to take advantage of this mechanism, suggesting a novel design principle linking synapse-resolution connectivity to network computation.

---

The mechanistic complexity of neural circuits exceeds our conceptual understanding but is becoming measurable, presenting both a profound challenge and a transformative opportunity for neuroscience. Recent advances in large-scale neuroanatomical and functional datasets have mapped brain circuits at unprecedented resolution, with comprehensive connectomes [1–7], largescale neuronal activity measurements [8–11], and celltype–resolved transcriptomic atlases [12, 13] now allowing the field to link molecular and structural substrates to circuit computations [14–18]. These successes suggest that to uncover this relationship, it is essential to develop theoretical frameworks that can interpret such highdimensional biological detail and identify the circuit-level principles that support robust and flexible neural representations and computations.

The head-direction (HD) system has long served as a model for understanding how neural circuits generate cognitive representations underlying navigation [19, 20]. The discovery of HD cells that were tuned to head direction [21] revealed that HD cell populations can accurately encode an animal’s orientation through a localized bump of activity that moves coherently with self-motion and spans a ring-like neural activity manifold [22, 23]. Theoretical studies have posited that these dynamics arise from ring attractor networks—continuous attractors [24– 27] in which recurrent excitation among similarly tuned neurons and broad inhibition collectively form and stabilize the activity bump [28–39]. However, canonical ring attractor models rely on idealized symmetry and hand-designed synaptic weights, assumptions that conflict with the biological variability and multi-population organization revealed by data. This discrepancy raises fundamental questions about how real circuits achieve precise and stable head-direction representations.

The *Drosophila melanogaster* HD system [42] provides a powerful model for resolving these tensions between theoretical simplicity and biological complexity (Figs. 1a to 1c). Its compact central complex architecture, comprising fewer than fifty EPG compass neurons circularly organized within the ellipsoid body, exhibits clear signatures of ring attractor dynamics, including persistent activity bumps and angular velocity integration [38, 43, 44]. Previous qualitative analyses have suggested that excitatory EPG neurons and inhibitory Δ7 neurons together provide recurrent excitation and broad inhibition that could conceivably stabilize a head-direction representation [16, 45, 46]. Yet it remains unknown whether the measured synaptic connectivity in the fly’s central complex is sufficient to realize a continuous attractor consistent with theory and behavior, and how such a small network robustly sustains stable head-direction representations is unclear [39].

**FIG. 1.**
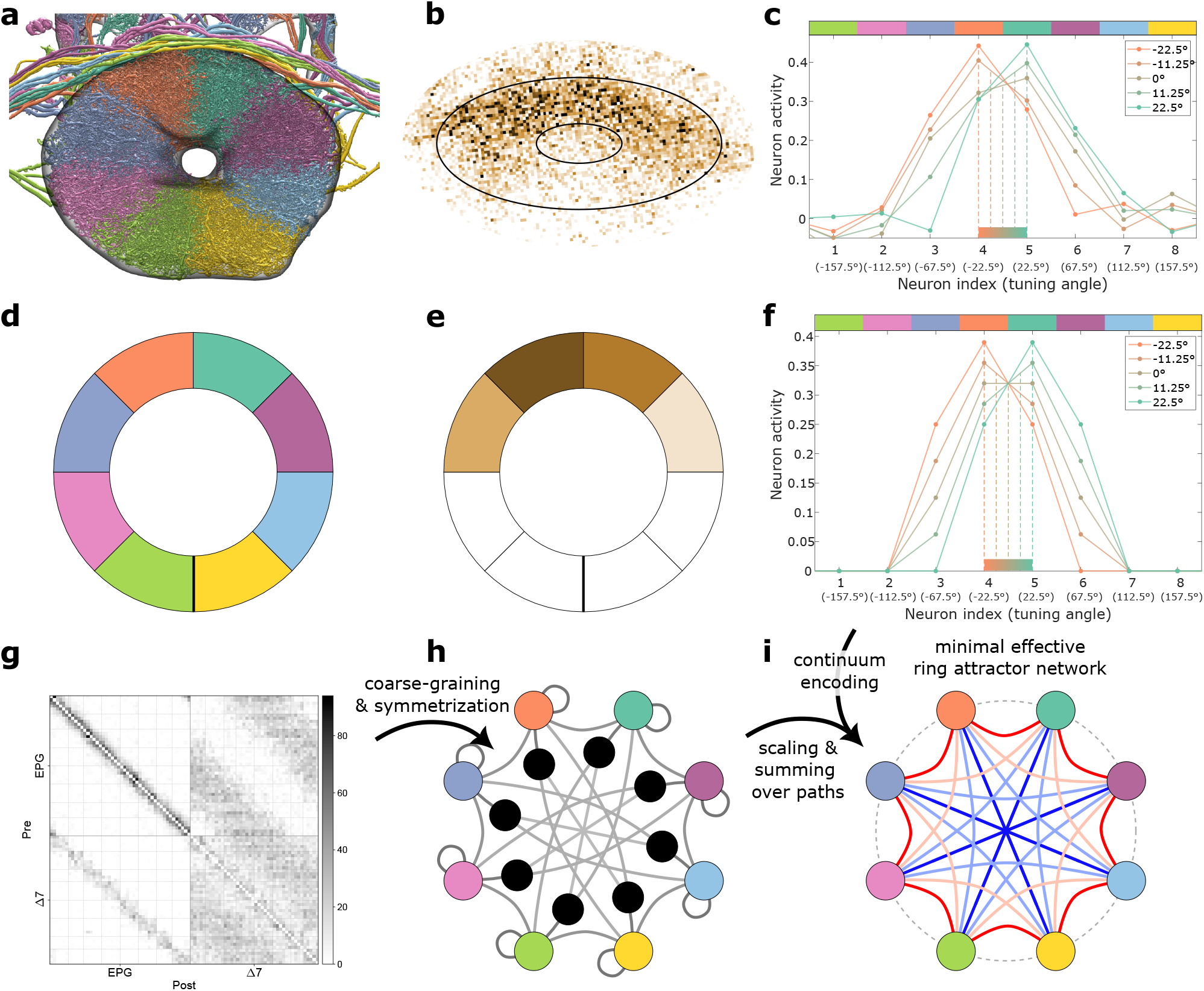
Constructing effective ring attractors from connectomic data. (a) EPG neurons from the hemibrain dataset [40] in the ellipsoid body (EB). Neuron processes are color-coded according to the EB wedges that they arborize. (b) Example calcium activity imaging data of the EPG neuron population in the EB. (c) Five experimentally observed activity profiles (solid lines) whose mean values (dashed lines) are distributed uniformly across the interval between the 4th and 5th wedges. (d) and (e) Schematics of the neural units’ identities and activities, respectively, according to the color codes in (a) and (b). (f) Example activity profiles with mean values matching the ones in (c), sampled from a continuum of solutions generated by a ring attractor model. (g) Synapse count matrix between the EPG and Δ7 neurons from the male CNS dataset [41]. Individual estimated wedges are separated by thin gray lines. (h) Schematic of a coarse-grained two-population model of an excitatory compass population (EPG, color-coded as in (d)) and an interconnected inhibitory population (black). (i) Schematic of an effective single-population ring attractor model derived by aggregating the two-population model and (h) parameterized to satisfy the continuum encoding conditions.

Here we develop a theoretical framework that bridges the gap between connectome-level structural detail [7, 40, 41, 46–49] and functional dynamics [38, 42–44, 46]. We start by characterizing the ensemble of synaptic weight matrices that allow a threshold-linear neural network to self-sustain a continuous ring attractor. We then map the EPG–Δ7 circuit onto this ensemble by converting synapse counts into synaptic weights by a few cell-type-specific scale factors [16, 18]. For each of the four available connectomes that contain the central complex - the Male CNS [41], hemibrain [40, 46], flywire-fafb [7, 47] and banc [49] datasets - we discover three shared connectome-constrained realizations of exact ring attractor dynamics that each reproduce the observed compass neuron activity but make different predictions for inhibitory neuron activity. We suggest that our mapping between connectomes and ring attractor networks endows robustness through modulability [50], as synaptic variations among and around the fly connectomes can be compensated by rescaling a few parameters through neuromodulation. Our analysis highlights the dimensions in connectome space most relevant for computation and reveals that the fly connectomes are remarkably well configured to exploit this mechanism. Our framework can be extended to incorporate additional cell types in the fly central complex and could reveal principles of neural circuits representing other continuous quantities, such as spatial location, across insects and vertebrates.

## Linking continuous attractors to connectomics

To foreshadow the analyses presented later in the paper, it’s helpful to provide a conceptual overview of our approach for constructing continuous ring attractor networks and determining whether such networks can be realized from the fly connectome.

To model the fly HD system, we consider a threshold-linear recurrent neural network comprising eight computational units or neurons (Fig. 1d), each corresponding to the mean activity of compass neurons observed in one neuronal compartment. Motivated by the experimental observation that approximately half of the neurons are active at any given time (Fig. 1b), we focus on configurations with four active neurons (Fig. 1e). The threshold-linear input–output function ensures that neurons receiving negative net input remain inactive, enabling the emergence of activity profiles that closely match experimental data (Fig. 1f). Since the activity bump persists for several seconds even in darkness—and because our goal is to construct a continuous ring attractor directly from experimentally measured recurrent synaptic connectivity—we require that the activity patterns representing head direction be self-sustaining and do not rely on any hypothetical external drives.

Our first goal will be to identify the ensemble of networks that can self-sustain a continuum of steady-state solutions spanning the full 360° of angular locations. Inspired by the connectomic data for the EPG compass neurons and Δ7 inhibitory neurons (Fig. 1g), we will consider a two-population network with 8-fold circular symmetry and reciprocal symmetry (both weights interconnecting two neurons of the same type are equal) (Fig. 1h). We will then identify *effective* networks involving just the compass neurons (Fig. 1i), where effective synaptic weights incorporate both direct synapses among the compass neurons and indirect pathways mediated by the inhibitory population. We will find that one class of effective networks has the same symmetry as the two-population network, resulting in only four independent effective weights (line colors in Fig. 1i, Appendix A). Somewhat surprisingly, the nonlinearity of the input–output function will also allow effective networks that break both original symmetries but retain a residual *mirror symmetry* (Appendix B). For both cases, we will be able to identify the full family of viable ring attractor networks.

Our second goal will be to determine whether any of these ring attractors are consistent with the measured fly connectome. We will analyze the EPG–Δ7 connectome (Fig. 1g), coarse-grained and symmetrized to match our modeling assumptions (Fig. 1h); note however that perfect symmetrization is not necessary (Supplementary Fig. S1, Appendix C). To convert synapse counts into synaptic weights, we will introduce four independent scaling parameters for the four classes of synaptic connections (EPG→EPG, EPG→Δ7, Δ7→EPG, and Δ7→Δ7). The key question is whether any choice of scaling parameters yields an effective network that is a continuous ring attractor (Fig. 1i).

## Ensemble modeling of exact ring attractors

We now proceed to our first goal of identifying the full ensemble of self-sustaining ring attractor networks. We begin by reducing the rotationally symmetric and reciprocally symmetric two-population model (Fig. 2a(i)) to an effective single-population model that has only compass neurons and reproduces all the steady-state configurations exactly (Appendix C). This reduction procedure can either preserve the symmetries of the two-population model (Fig. 2a(ii)) or break them and retain mirror symmetry (Fig. 2a(iii)). Symmetry breaking arises when inhibitory neurons become inactive, as this implies that an indirect synaptic pathway between two compass neurons can exist in one direction but not the other. For instance, Fig. 2a illustrates this mechanism for our network of eight neurons, indexed by *i* = 1, …, 8, where the active set is {3, 4, 5, 6} and we show contributions to the effective weights between neurons 3 and 4. Consequently, the effective connectivity among the four active neurons is parameterized by three independent weight parameters in the symmetric case (Fig. 2b) and six in the mirror-symmetric case (Fig. 2c). Note that we set all effective self-synapses to zero without loss of generality (Appendix D).

**FIG. 2.**
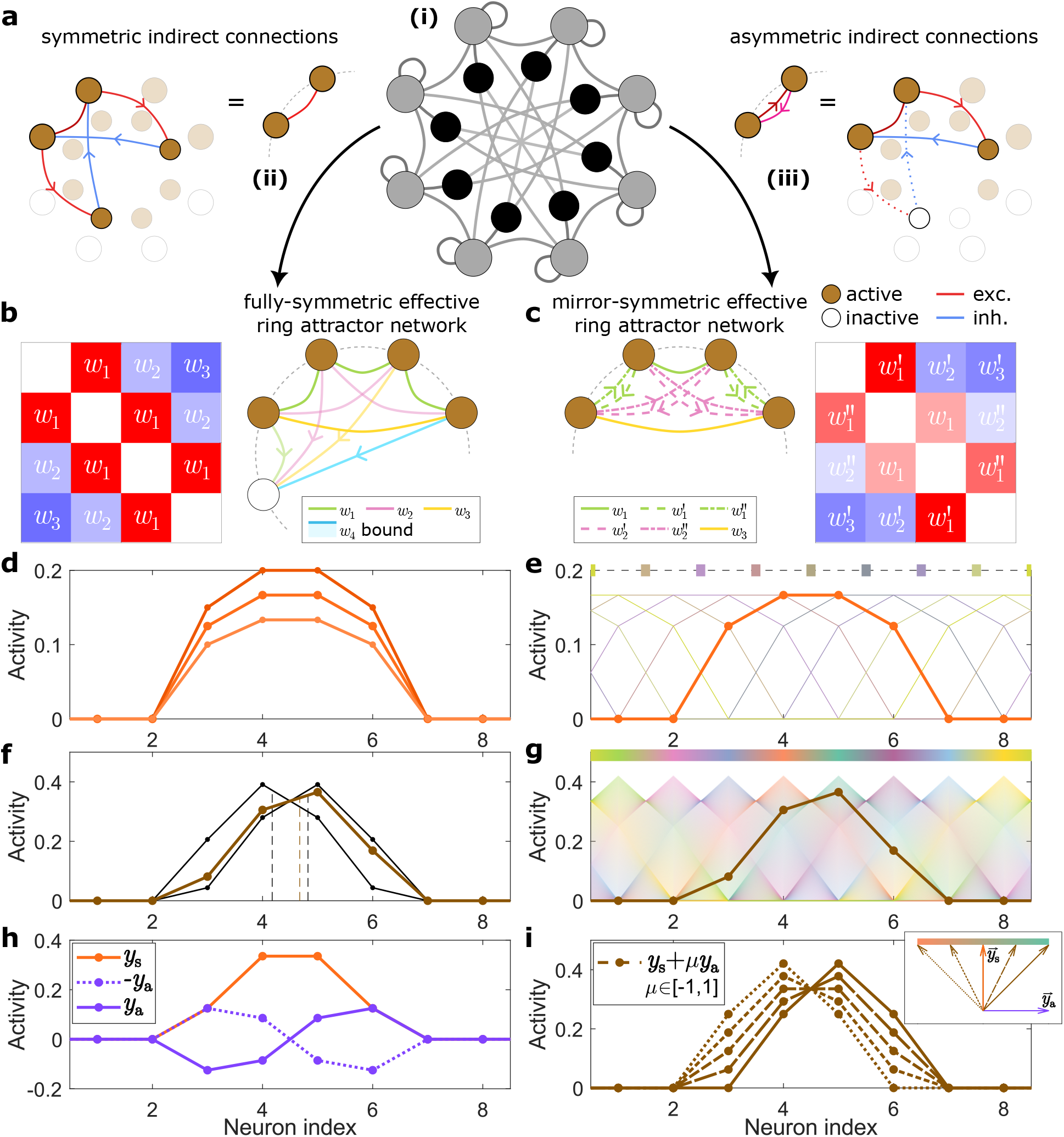
Requirements for effective ring attractor models to have the continuum-encoding property. (a) The two-population model (i) can be aggregated into two distinct effective ring attractor networks: (ii) the fully-symmetric network, by assuming all neurons in the second population (inner circle) are active, or (iii) the mirror-symmetric network, arising from asymmetric indirect connections when there are inactive neurons in the second population (white nodes in inner circle) in the steady state. In (iii) dotted lines denote the absent path. (b) and (c) Effective matrices for the four active neurons of the primary population for the fully- and mirror-symmetric models, respectively. (d) Example symmetric profile for the single-degeneracy case. Darker/lighter scarlet profiles depict the flexibility in amplitudes. (e) Eight total profiles and their respective mean values at discrete uniformly-distributed locations (top) for the single-degeneracy case. (f) Example profile (brown) (as in (d)) for the double-degeneracy case, which enables asymmetric mirrored pairs of basis profiles (black). (g) Continuum of profiles and their mean values (top) span the whole angular space. (h) Symmetric (scarlet) and antisymmetric (violet) profiles define the basis vectors for the activity profiles (i, eigenvectors in inset). (i) Different values of *µ* ∈ [−1, 1] define different steady-state profiles (or vectors, inset) in the specific angular interval (between neurons 4 and 5 in the figure).

A self-sustaining bump of activity requires that the steady-state activity of every active neuron equals the total recurrent input it receives from other neurons. If 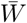 denotes the effective weight matrix between active neurons, this implies that the steady-state activity profile is an eigenvector of 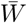 with eigenvalue 1. If 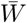 has only one such linearly independent eigenvector, that eigenvector must be symmetric (Appendix E). Although the bump amplitude may vary (Fig. 2d), this solution does not represent a ring attractor because it can only stabilize the activity bump at eight discrete angles (Fig. 2e). If the eigenvector with eigenvalue 1 is asymmetric, then the model’s symmetries imply that its mirror-reflected partner is also an eigenvector with eigenvalue 1. Together these two vectors span a two-dimensional space of eigen-vectors with eigenvalue 1 (Fig. 2f). This solution provides a ring attractor where the bump amplitude and shape can vary continuously to represent the continuum of angles (Fig. 2g). We note that continuous encoding can also be achieved with a single eigenvector having eigenvalue 1 if we do not require the bump to be self-sustaining, but we find that variations in the external drive can compromise angular encoding (Appendix F, Supplementary Fig. S2).

The two-dimensional space of eigenvectors with eigen-value 1 can be conveniently understood in a simple and interpretable basis. Because the eigenspace is two-dimensional, it has both a symmetric eigenvector 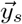 and an antisymmetric eigenvector 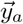 (Fig. 2h, Appendix E). Adding 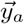 to 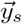 yields an asymmetric activity profile, as 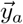 enhances activity on one side and suppresses it on the other. The relative weighting of the symmetric and asymmetric eigenvectors thereby sets the angular location of the bump, whereas the sum of their weights sets the activity amplitude (Fig. 2i). In this perspective, a finite range of bump locations arises for a given active set because the asymmetric eigenvector promotes negative activity that cannot be realized in a threshold-linear network. The magnitude of its weighting can therefore not exceed that of the symmetric eigenvector. Overall, the steady-state profiles are parameterized by two free parameters—an amplitude *σ* > 0 and a location parameter *µ* ∈ [−1, 1], which linearly encodes the angular location over the active set’s required 45° range.

Specifying the shape of the bump requires only one additional parameter, *r*_*a*_. This parameter corresponds to a ratio between the activity levels of the fourth and third (same as between fifth and sixth) neurons in the antisymmetric eigenvector, where we’ve again assumed that the active set is {3, 4, 5, 6} . This quantity is fully determined by the model’s effective weights and is therefore fixed for each ring attractor (Appendices A, B). As *r*_*a*_ increases, the bump becomes increasingly peaked (Figs. 3a and 3b). In the symmetric model, *r*_*a*_ is restricted to the range (0, 1) (Fig. 3a), whereas mirror-symmetric networks can exhibit *r*_*a*_ ≥ 1, producing even sharper activity profiles (Fig. 3b). This difference offers an experimental signature for distinguishing the two classes of ring attractors.

**FIG. 3.**
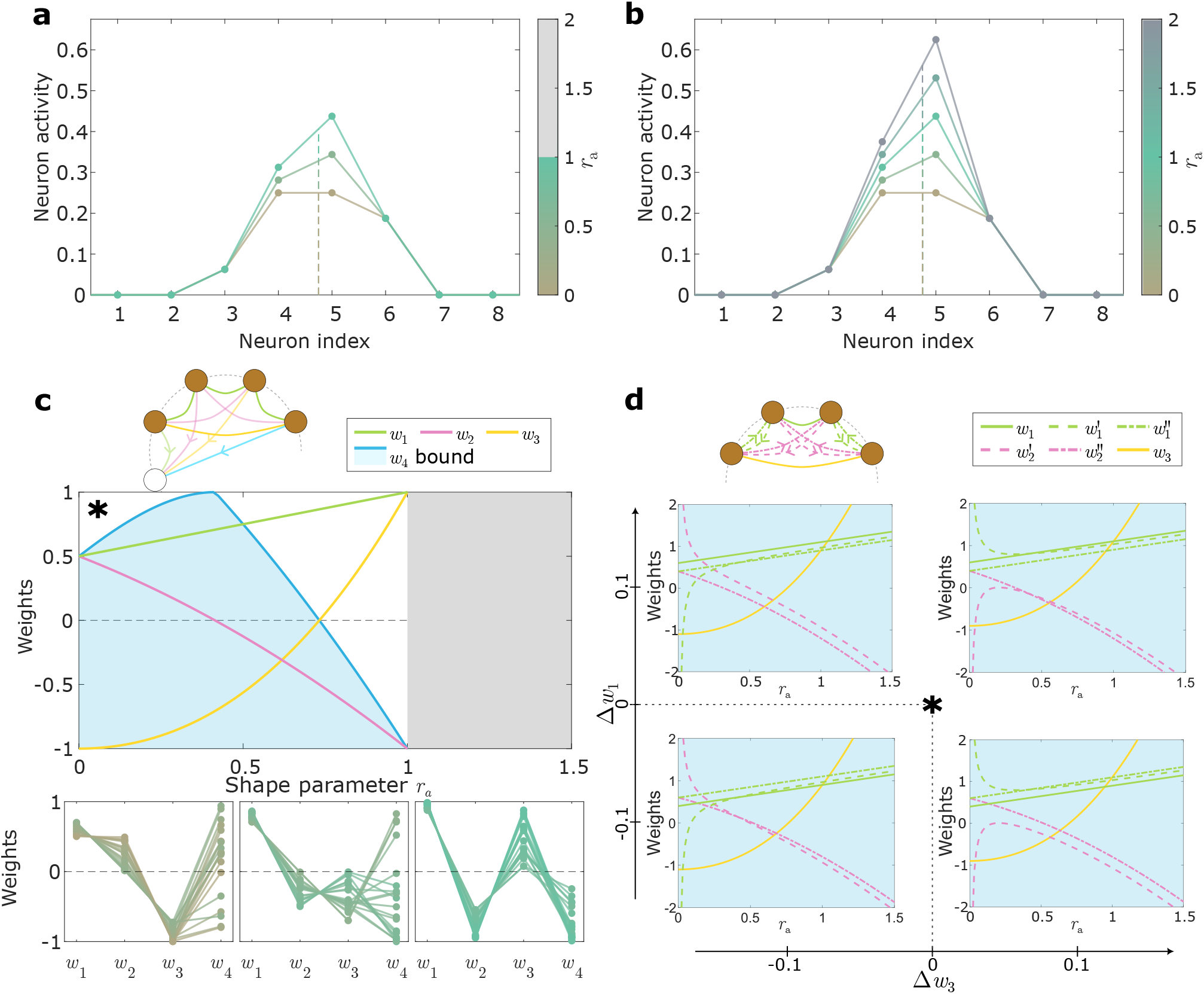
Solution space of effective ring attractor models. The shape parameter *r*_*a*_ ties the shapes of the weight and activity profiles. (a) and (b) Dependence of the shape of the neuron activity profiles on *r*_*a*_. (a) *r*_*a*_ ∈ [0, 1] for the fully-symmetric case and (b) *r*_*a*_ ∈ [0, 2] range for the mirror-symmetric case, which allows *r*_*a*_ > 1 unlike the symmetric model. (c) and (d) Dependence of the weights on the *r*_*a*_ parameter values for the fully- and mirror-symmetric cases, respectively. Sample weight profiles are shown for the fully-symmetric case (c, bottom) for three segments of *r*_*a*_ grouped by which weights are inhibitory (*w*_2_ *>* 0, *w*_3_*<* 0, left; *w*_2_ *<* 0, *w*_3_*<* 0, middle; *w*_2_ *<* 0, *w*_3_ *>* 0, right), demarcated by the sign-changing points of 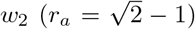 and 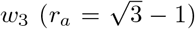. Values for *w*_4_ were randomly sampled between -1 and its upper bound dependent on *r*_*a*_ (blue region). Weight profile shapes for the mirror-symmetric case (d) are dependent on two additional parameters Δ*w*_1_ ≡ *w*_1_ − *w*_1,sym_(*r*_*a*_) and Δ*w*_3_ ≡ *w*_3_ − *w*_3,sym_(*r*_*a*_).

Fig. 3c fully characterizes the symmetric ring attractor ensemble (Appendix A). Bump stability and the eigenvalue constraints uniquely determine the three recurrent weights between neurons in the active set, *w*_*i*_ for *i* = 1, 2, 3, as functions of *r*_*a*_ (Fig. 3c, top). The self-consistency of the active set also requires that all inactive neurons receive non-positive input in every steadystate profile, which imposes an inequality constraint on *w*_4_ as *r*_*a*_ varies. The resulting solution space is two-dimensional. Notably, *w*_1_ must always be excitatory, while at least one of *w*_2_ or *w*_3_ must be inhibitory. Thus the model naturally captures the intuitive motif of local excitation and distal inhibition, albeit allowing substantial flexibility in the inhibitory components. The parameter *w*_4_ is comparatively unconstrained for most *r*_*a*_ values, but it must be negative when *r*_*a*_ is large and *w*_3_ becomes positive. Representative weight profiles are plotted in Fig. 3c (bottom).

The mirror-symmetric model is significantly more flexible (Fig. 3d, Appendix B). Here the recurrent weights between neurons in the active set can vary across a three-dimensional manifold: in the six-dimensional space of active set weight parameters, two are fixed by the eigen-value conditions and one by requiring that each active set encodes angles over a 45° range ensuring smooth transition from one active set to the next. Moreover, the weights involving the inactive neurons are barely constrained: they are independent of the weights between active neurons, mirror symmetry only reduces the number of weight parameters by half, and the non-positivity of inputs to the inactive set neurons only imposes inequality constraints. Overall, the ensemble of stable mirror-symmetric ring attractors is 25-dimensional, 28 independent weight parameters satisfying the two eigenvalue constraints and one angular-range/smoothness condition.

## Consistency with the EPG-Δ 7 connectome

Having constructed the ring attractor ensemble, we now examine whether synaptic interactions between EPG and Δ7 neurons (Fig. 4a) can realize a ring attractor network. The EPG neurons and Δ7 neurons comprise eight anatomical compartments containing four to six computationally similar neurons [45], and their recurrent connectivity exhibits an approximate eight-fold circular symmetry (Fig. 1g). Because our theoretical framework requires exact symmetry, *C*_*ij*_ = *C*_*i*+1,*j*+1_, we symmetrized the synapse count matrix between compartments by averaging all elements assumed to be equal (Fig. 4b, Methods).

**FIG. 4.**
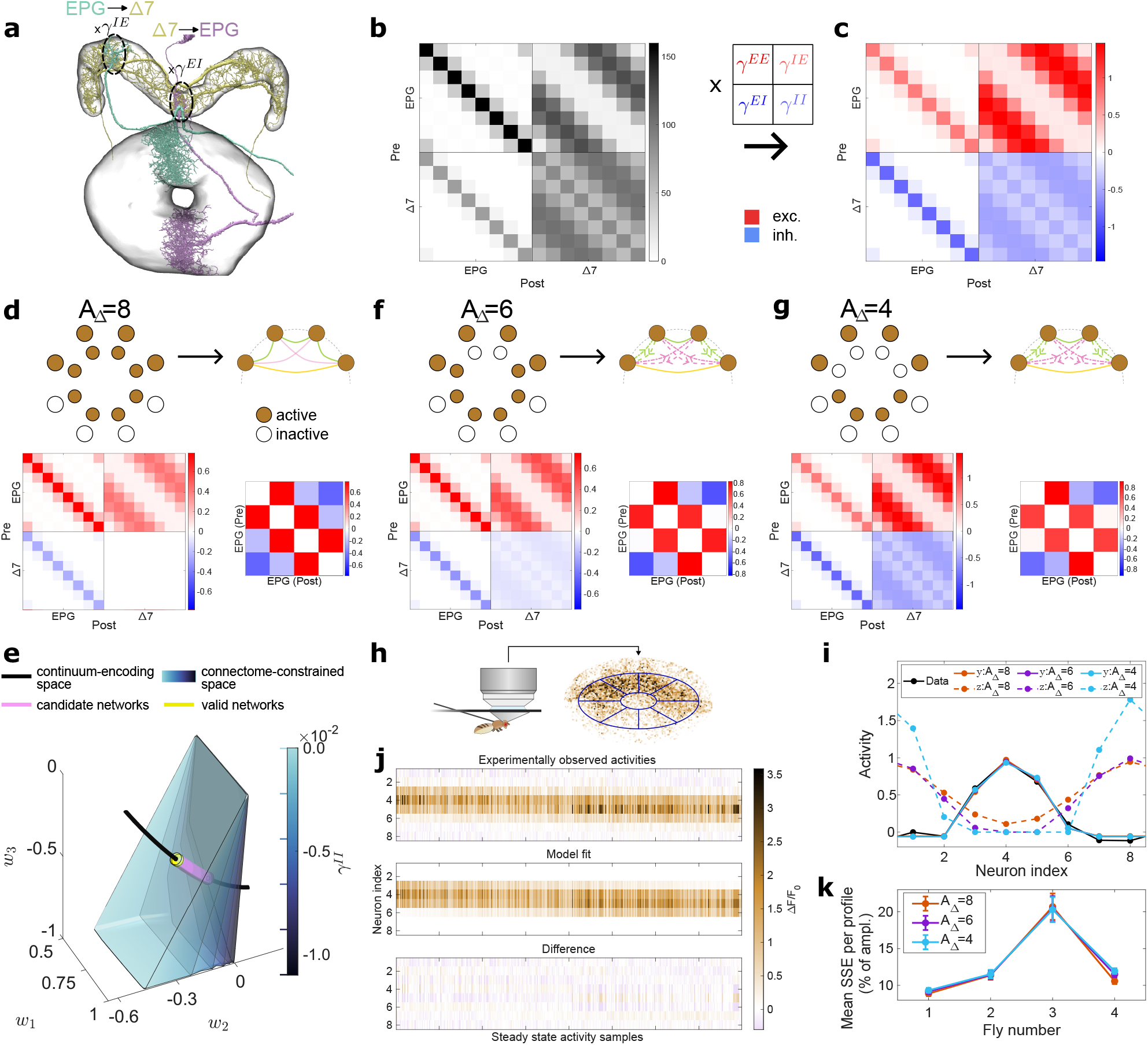
Several exact ring attractors match the EPG-Δ7 connectomic data. 4(a) EPG (from the EB) and Δ7 neurons link with each other in the protocerebral bridge (PB), depicted using the hemibrain dataset [6]. (b) Coarse-grained synaptic count matrix between the EPG and Δ7 neurons obtained from the male CNS connectome [41]. Each block submatrix is parameterized with a single *γ*-parameter that scales the values within that submatrix, to generate the weight-matrix in (c). Red - excitatory, blue - inhibitory weights. (d) Top: Two-population network (left; outer layer - EPG, inner layer - Δ7 population) with *A*_Δ_ = 8 active Δ7 neurons aggregates to an effective fully-symmetric network of active EPG neurons (right). Bottom: Corresponding solution and its effective weights. Value of *γ*^*II*^ = 0 was used. (e) Intersection points (pink line) between the theoretical conditions of the continuum- encoding space (black) and connectome-constrained space (blue-grey volume) yield candidate weight profile solutions of ring attractors. Further inequality constraints on the activity profiles narrow down the solution space of valid ring attractors (yellow). (f) and (g) Same as in (d), but for the cases with *A*_Δ_ = 6 and *A*_Δ_ = 4 active Δ7 neurons - mirror-symmetric effective networks. (h) Left: Schematic of the setup for two-photon calcium imaging of the EPG neuron population in the EB for tethered immobile flies in darkness. Right: Mean Δ*F*/*F* values of the calcium signal was computed in 8 regions of interest (ROIs) around the EB for each time point. (i) Example model fits for a single EPG activity profile (solid lines) and the respective predictions for the activity of the Δ7 population (dashed) for the three different models in (d, f, g). (j) Top: Activity profiles for different time points during which the EPG bump position was relatively stationary. Activity profiles were circularly shifted to align the mean position in the middle segment and were also sorted by it. Middle and bottom: Corresponding fits with the fully-symmetric model and their differences to the data, respectively. (k) Summary statistics (mean*±*s.e.m.) of the fitting data with the three aforementioned models for multiple flies. The SSE per profile is normalized to the bump amplitude.

Similar to previous *Drosophila* models [16, 18], we hypothesize that synapse strengths are proportional to synaptic counts with a proportionality constant that may depend on both the pre-synaptic and post-synaptic cell type (Figs. 4b and 4c). We therefore introduce four scale factors, *γ*^*ab*^, to relate synapse counts to synaptic weights, where *a, b* ∈ {*E, I*} index the post-synaptic and pre-synaptic cell types, *E* refers to excitatory EPG neurons, and *I* refers to inhibitory Δ7 neurons. However, because the reduction to an effective network always involves the product of *E*→*I* and *I*→*E* synapses, only *γ*^*EIE*^ ≡ *γ*^*EI*^ *γ*^*IE*^, appears in the effective network. Thus the connectome maps onto a three-dimensional space of realizable effective EPG networks.

For all four connectomes, we discovered three sets of scale factors that generate ring attractor dynamics (Supplementary Fig. S3). The first solution generates a fully symmetric effective model by requiring that all eight Δ7 neurons be simultaneously active (Fig. 4d). Dimensionality arguments suggest that we should expect a one-dimensional family of fully symmetric connectome-constrained solutions (purple curve,

Fig. 4e, Appendix C), because we have three flexible scale factors (*γ*^*EE*^, *γ*^*II*^, *γ*^*EIE*^) that have to satisfy the two eigenvalue conditions to realize a ring attractor. However, it is hard to achieve the constraint that all Δ7 neurons be active at once because there are so many inhibitory synapses between them, and the *γ*^*II*^ scaling parameter must therefore reduce the Δ7–Δ7 recurrent coupling to near-zero levels (Fig. 4d). This essentially yields a single fully symmetric solution (short yellow curve, Fig. 4e). The other two solutions generate mirror-symmetric effective models by requiring that six or four Δ7 neurons be active at any given orientation (Figs. 4f and 4g). Here the dimensionality arguments correctly predict that each of these solutions is a zero-dimensional point (Appendix C); the three scale factors need to satisfy two eigenvalue and one angular range condition. All scale factors are of comparable magnitude in the mirror-symmetric models.

These models allow us to predict the activity of EPG and Δ7 neurons in behaving flies, from connectomics data. To start testing these predictions, we measured the steady-state shape of the EPG activity bump when the fly was stationary with a wide range of orientations (Fig. 4h). Intriguingly, the values of the shape parameter for all three connectome-constrained networks turn out to be similar (*r*_*a*_ = 0.548, 0.437, and 0.391, for *A*_Δ_ = 8, 6, and 4, respectively) leading to very similar EPG activity profiles that closely matched the experimentally measured patterns (solid lines, Fig. 4i). Moreover, we could account for each measured EPG steady-state activity profile by varying only the amplitude parameter *σ* and location parameter *µ* of the models (Fig. 4j), with all three networks providing similarly good fits (Fig. 4k). In contrast, the three models produced clearly distinguishable Δ7 activity profiles with different numbers of active Δ7 neurons (dashed lines, Fig. 4i). Future experiments should measure Δ7 neuron activity to experimentally differentiate the models.

## Robust encoding through synaptic rescaling

The effective weights in our models must be precisely tuned to generate ring attractor dynamics. However, a biologically plausible encoding strategy must be able to tolerate natural variability in synaptic connectivity (Fig. 5a) [51]. We thus asked whether the four scale factors relating synapse counts to synaptic weights provide enough flexibility to retune a perturbed connectome back to a ring attractor (Fig. 5b). Remarkably, even when synapse counts in the EPG-Δ7 connectome varied by nearly 90%, we were almost always able to find scale factors that transformed the symmetrized count matrix back into a ring attractor (Fig. 5c, Methods).

**FIG. 5.**
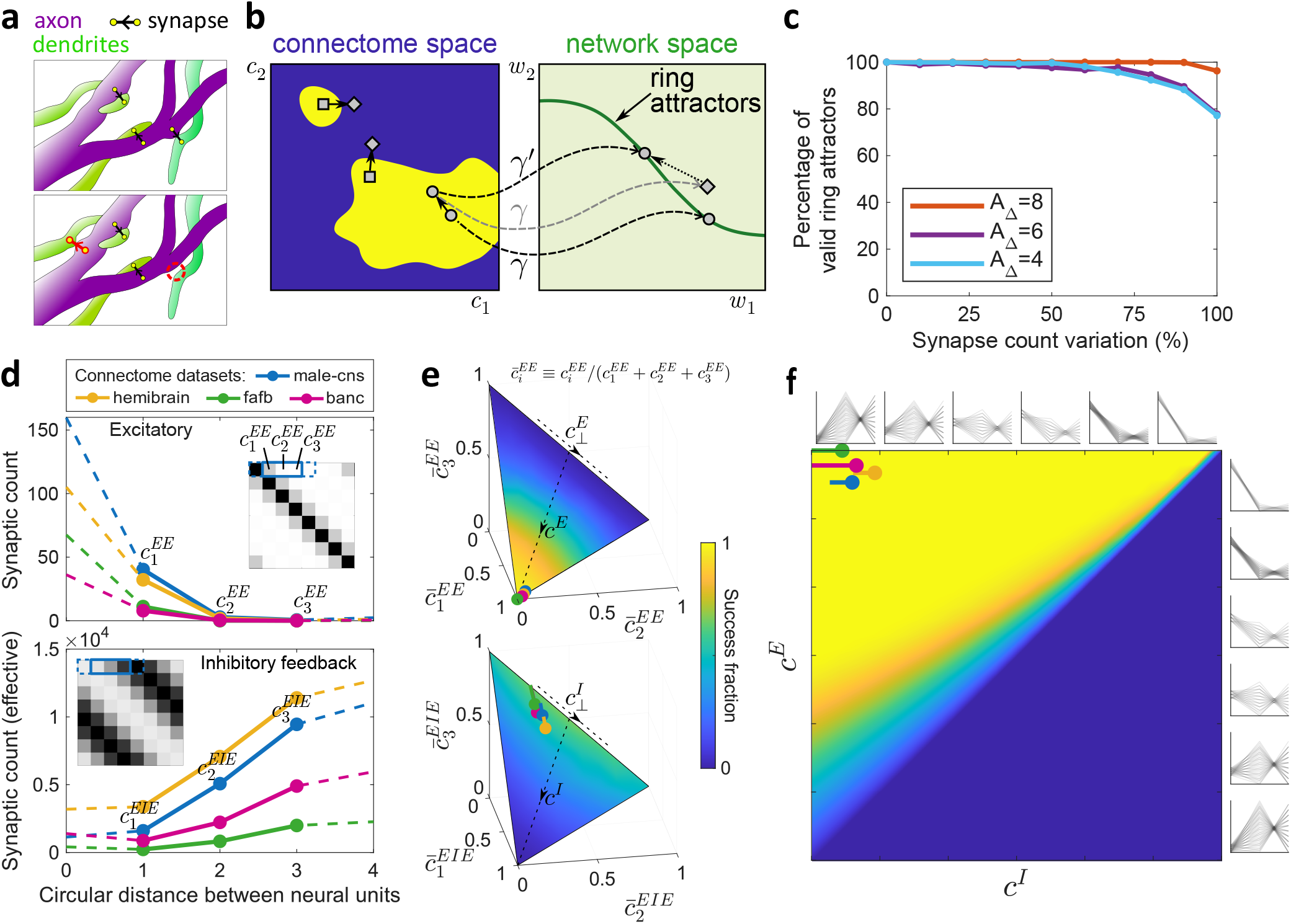
The fly connectome exibits robust ring attractor dynamics through its modulability. (a) Schematic drawing showing a possible scenario of changes over time (red) in axonal synaptic connectivity with nearby dendrites. (b) The yellow region indicates connectomes that can be mapped to a ring attractor network. Fluctuations in the connectome (○) may be compensated by changes in the scale factors ({*γ*}) to ensure that the effective network weights continue to realize a continuous ring attractor. However, this may not be possible if the connectomes (□) are poorly located. (c) Simulation results showing the success percentage of realizing the three different continuous ring attractors with *A*_Δ_ ∈ {4, 6, 8} as we vary the synapse counts (while maintaining circular symmetry) around the male CNS connectome [41]. Each synapse count parameter was uniformly randomly sampled in the *±*(% var.) range, and the success percentage computed as the percentage of successful mappings over 1000 samples per % variation. (d) The space of circularly symmetric fly connectomes can be described succinctly with an excitatory (top) and an inhibitory (bottom) connectivity profile, where each profile consists of five numbers that uniquely characterize the corresponding synapse count matrices and colored curves correspond to the four measured connectomes. (e, top) The success fraction (color bar shared between (e) and (f)) varies with the three normalized synapse counts, 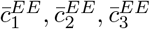, parametrizing the excitatory matrix. Here the success fraction is computed by averaging over the three normalized synapse parameters, 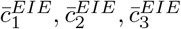 characterizing the inhibitory matrix. (e, bottom) This depicts success fraction as one varies, 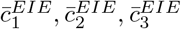, while averaging over 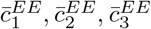. The colored circles on both graphs correspond to the different fly connectomes, and the tails indicate nonzero values for *γ*^*II*^ . (f) Here we plot the average success fraction as we vary the two median directions, *c*^*E*^, *c*^*I*^, as depicted in (e). The averaging is now performed over the two remaining orthogonal synapse count directions, 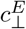 and 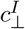. The colored circles and tails represent the connectomes as in (e). On the top and righthand side we depict the inhibitory and excitatory synaptic profiles that we average over.

We next sought to investigate how the robustness of the fly connectome compares to hypothetical non-biological alternatives. For simplicity, we examined this first for fully symmetric effective models and set *γ*^*II*^ = 0 (Fig. 4d). This implies that any connectome can be characterized by ten independent synapse-count parameters (Fig. 5d), five describing the EPG-EPG synapse count matrix, *C*^*EE*^, and five describing the product of the EPGΔ7 and Δ7-EPG count matrices, *C*^*EIE*^. We further reduce the dimensionality by rescaling the synapse counts and eliminating the shortest and longest range connections (Methods). This casts the space of connectomes as a four-dimensional manifold that factorizes into excitatory and inhibitory components (Fig. 5e). We note that we also have a symmetric effective network whenever *γ*^*II*^ is sufficiently small that the Δ7 neurons remain active. We also characterized these *γ*^*II*^ ≠ 0 cases by *C*^*EE*^ and an effective inhibitory connectivity similar to *C*^*EIE*^, except that the inhibitory connectivity now incorporated recurrence between Δ7 neurons (Methods).

The position of the fly data in this connectome space makes it near-optimal for robustness under synaptic variation, which explains why all available fly connectomes can realize ring attractor dynamics despite natural biological variability. To show this, we densely scanned the four dimensions of the connectome space and asked whether each connectome could support a ring attractor through some set of synaptic scale factors (Methods). Given the fly’s EPG-EPG connectome, almost any pattern of EPG-Δ7-EPG synaptic connectivity could enable ring attractor dynamics (Fig. 5e, top). In contrast, only a fraction of EPG-EPG connectomes permit ring attractor dynamics with the fly’s EPG-Δ7-EPG connectivity (Fig. 5e, bottom). This quantification revealed two directions in connectome space that heavily influenced whether a connectome was consistent with ring attractor dynamics, which we denote as *c*^*E*^ and *c*^*I*^ (Fig. 5e). These directions interpolate between having only local connectivity 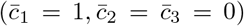 to having only distal connectivity 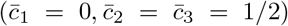. Given fly connectomes’ positions on these axes, we found that the other two dimensions of the connectome could vary freely without affecting performance (Fig. 5f). Furthermore, the fly connectomes were located deep within the region of parameter space where ring attractor dynamics were consistently achievable, ensuring robustness of the head-direction system. A quadratic discriminant analysis indicates that the fly connectome is also well positioned to generate mirror-symmetric ring attractor networks (Methods, Supplementary Fig. S4). This suggests that evolution has found a connectome that permits robust ring attractor dynamics through synaptic rescaling.

## Discussion

We show that continuous ring attractor dynamics can emerge directly from experimentally measured recurrent connectivity. Although our analysis focused on the EPG-Δ7 system, we identified double degeneracy of the effective weight matrix as the key structural condition for self-sustained continuous encoding. This insight generalizes to alternative connectome-constrained architectures, including broad inhibition from a single ER6 ring neuron [45] (Appendix G, Supplementary Fig. S5) and multi–cell-type networks (Appendix H). Our discovery of mirror-symmetric ring attractor networks adds to growing evidence that exact symmetry is not required for ring attractor dynamics [52–55].

Future connectome-constrained models could enhance our understanding of the head-direction circuit by detailing how it is shaped by its inputs and used by downstream circuits to generate behavior. Upstream, a key functional requirement of ring attractor networks is their ability to integrate velocity signals [28, 38]. EPG neurons also receive sensory inputs through ring neurons that anchor the bump to the external world and help calibrate arbitrary gain factors in the velocity integration circuit [42, 56–59]. While a detailed connectome-constrained implementation of these processes lies beyond the scope of this paper, Appendix I illustrates how our framework could accommodate them. Downstream, EPG/Δ7 signals are transmitted to the fan-shaped body, where vector computations involving various relevant directions enable goal-directed navigation [16, 60, 61]. While EPG neurons do not exhibit the sinusoidal activity pattern hypothesized to underlie such transformations [16, 46, 62– 65], the Δ7 activity pattern predicted by our fully symmetric model does. The predicted Δ7 activity in our mirror-symmetric models are less sinusoidal but may be more energetically efficient, as some Δ7 neurons remain inactive. We hypothesize that the head-direction system may dynamically switch between these ring attractor realizations to meet the state-dependent demands of down-stream computation.

Although detailed connectomic data from *Drosophila* motivated this study, our theoretical framework generalizes naturally across species. The insect central complex exhibits remarkable evolutionary conservation, as demonstrated by comparative analyses of cockroach [66] and locust [62] HD systems. Our model predicts that continuous angular encoding can arise in any species as long as the effective weight matrix exhibits double degeneracy, independent of species-specific differences in neuron number, compartmentalization, or synaptic detail. Strikingly, a recently discovered HD system in larval zebrafish shows unexpected parallels with the fly compass [67–69]. This cross-species convergence provides a natural bridge for understanding ring-attractor dynamics in vertebrate systems.

While we focused here on a specific cognitive variable—head direction—the structural principles we identify are not restricted to angular representations. For a given active set of neurons, our framework characterizes the ensemble of effective weight matrices capable of encoding any continuous variable, such as spatial location, goal direction, gaze angle, or reach or movement direction. A major open challenge is how to impose the correct topology of the encoded variable onto these networks so that their dynamics faithfully reflect the geometry of the represented space [23, 24, 70].

We found that even large perturbations in synapse counts around the fly connectome could be corrected by adjusting a few global scaling factors. Moreover, this robustness was not a generic feature of all hypothetical connectomes that could realize a ring attractor; the fly connectome seems to be ideally positioned to tolerate large fluctuations. Indeed, this robust location was crucial for accommodating natural biological variability across individual connectomes and successfully mapping them onto a ring attractor network. Converging evidence suggests that neuromodulation may play a key role in stabilizing attractor function [71, 72]. In line with our hypothesis, multiple neuromodulation-mediated homeostatic mechanisms have been proposed to mitigate the fine-tuning challenges of attractor networks by up- or down-regulating synaptic strengths [50, 72–74]. For instance, neuropeptides can enhance neuronal excitability and strengthen synaptic connectivity, whereas disrupting neuropeptidergic transmission weakens attractor dynamics [72]. In our model, perturbations to the connectome (Fig. 5b) cause the activity bump to decay in characteristic ways that could be detected by the system. We hypothesize that this could trigger compensatory changes of the global scaling factors via neuromodulation—thereby ensuring a robust and accurate head-direction representation in the fly.

## Methods Theoretical Model

We model the effective ring attractor dynamics according to the firing rate network model:

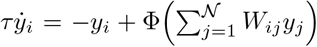, where *τ* is time constant for the firing rate dynamics, *W*_*ij*_ denotes the synaptic weight from neuron *j* to *i*, and Φ(*x*) = max(0, *x*) is the threshold-linear nonlinearity. Our analysis mostly focuses on self-sustained steady-state profiles that satisfies, 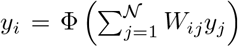. When analyzing dynamics, we set *τ* = 1. Detailed analysis and relevant generalizations of the modeling equations are provided in Appendices A through J, various notations and conventions are also summarized in a supplementary table.

### Experimental procedure

#### Fly preparation

Imaging experiments were performed on 6-8 days old female UAS-jGCaMP7f;SS00096-GAL4 flies, reared at 23°C in 60% relative humidity with a 16:8 light:dark cycle on standard cornmeal fly food. To express the genetically encoded calcium indicator jGCaMP7f [75] in EPG neurons specifically, jGCaMP7f flies (20XUAS-IVS-Syn21-jGCaMP7f-p10 in VK00005; RRID:BDSC 79031) were crossed with the stable split EPG GAL4 driver line SS00096 [76]. The flies were prepared for imaging as previously described [42, 77]. Briefly, flies were anesthetized at 4^°^C, their proboscis was fixed with wax to reduce brain movements, their thorax was UV-glued to a tether pin for stability and manipulation, and they had their legs removed and the stumps and dorsal abdomen glued to reduce spontaneous motor activity. The fly’s head was positioned with a micromanipulator in a holder with a recording chamber and UV-glued to it to immobilize the head for dissection and brain imaging. To gain optical access to the brain, a section of cuticle between the ocelli and antennae was removed, along with the underlying fat tissue and air sacs. Throughout the experiment, the head in the imaging holder was submerged in saline containing (in micromolar): NaCl (103), KCl (3), TES (5), trehalose (8), glucose (10), NaHCO3 (26), NaH2PO4 (1), CaCl2 (2.5) and MgCl2 (4), with a pH of 7.3 and an osmolarity of 280mOsm.

#### Two-photon calcium imaging

For each fly we collected imaging data during trials in darkness. Calcium imaging was performed with a custom-built two-photon microscope controlled with ScanImage (version 2022, Vidrio Technologies) [78]. Excitation of jGCaMP7f was generated with an infrared laser (920nm; Chameleon Ultra II, Coherent) with approximately 15*mW* of power, as measured after the objective (×60 Olympus LUMPlanFL/IR, 0.9 numerical aperture). Fast Z-stacks (eight planes with 8*µm* spacing and three fly-back frames) were collected at 10Hz by raster scanning (128 × 128 pixels, ∼75 × 75*µm*^2^) using an 8kHz resonant-galvo system and piezo-controlled Z positioning. Focal planes were selected to cover the full extent of EPG processes in the EB. Emitted light was directed (primary dichroic: 735, secondary dichroic: 594), filtered (filter A: 680 SP, filter B: 514/44) and detected with a GaAsP photomultiplier tube (H10770PB-40, Hamamatsu).

### Data analysis and model fitting

The recorded XYZ stacks of EPG activity in the EB were first averaged over the Z dimension, and then drift-corrected over time for alignment. The aligned XYZ stacks were then averaged over time to get a mean EPG activity profile in 3D. To account for the 3D orientation of the EB within the brain, relative to the objective angle, a 3D mask of the EB was constructed from the masks in each individual slice separating the EB signal pixels from background. PCA was then performed on the 3D signal intensity points within the EB mask to find the optimal plane on which the EB signal should be projected. An ellipse was fitted through the projected data, to account for the XY rotation of the EB, and automatically segmented into 8 wedges, similarly as in Fig. 4i, top. Average fluorescence signal *F* ^*i*^(*t*) was calculated for every wedge 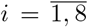 and time point 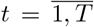, and the baseline fluorescence signal *F* ^*i*^_0_ per wedge was calculated as the 10^th^ percentile from the *F* ^*i*^(*t*) values. Finally, the Δ*F*/*F*_0_ = (*F* − *F*_0_)/*F*_0_ values were calculated for every wedge and time point, giving an 8 × *T*, Δ*F*/*F*_0_ data matrix.

We calculated the population vector average (PVA) at each time point *t* to estimate the EPG bump amplitude and orientation, as the circular mean of the 8-dimensional Δ*F*/*F*_0_ vector at *t* and the corresponding wedge angles (CircStat toolbox [79]). Time points where there is a lack of bump strength, as measured by the likelihood of uniformity of the circular data (p-value⩽0.5 of Rayleigh’s test; CircStat), were excluded from the analysis. A moving average with 11 frames of the PVA angle was used to estimate the instantaneous bump velocity. Time points where the bump velocity was greater than 20^°^/*s* were excluded from the analysis to filter out the non-stationary states.

To estimate the model fit of each of the three models (with *A*_Δ_ = 8, *A*_Δ_ = 6 and *A*_Δ_ = 4) on the EPG activity data, we used their estimated values for the spatial parameter *r*_*a*_. For each time point we then estimated the *σ* amplitude and *µ* position of the bump using (A.25, Appendix A), as well as an additional offset parameter to accommodate the baseline. Non-linear least squares (*lsqnonlin*, Matlab) was performed over the Δ*F*/*F*_0_ data circularly shifted for each of the 8 possible consecutive activity sets of EPG wedges, and the best fit was chosen for that time point and the active EPG set stored. Predictions for the Δ7 activities were subsequently generated using Eq C.66 (Appendix).

### Constructing connectome-derived ring attractors

To estimate the average synaptic strength in the connectivity between and within the computational units of EPG and Δ7 cell types from the male CNS dataset [40, 46], similarly as in [45], we estimate the average synaptic strength at the level of wedges in the EB (between EPG neurons) and glomeruli in the PB (EPG-Δ7 and between Δ7). Each Δ7 was assigned to a glomeruli (wedge) based on the maximal cumulative output of synaptic counts to EPG neurons grouped by wedges. The connections to every neuron were summed per presynaptic wedge to calculate the grouped inputs per neuron, and then postsynaptic neurons’ connectivity was averaged per wedge to calculate a final wedge-to-wedge asymmetric connectivity matrix. Average distance-based wedge-to-wedge connectivity vectors were calculated from this matrix and redistributed back to obtain a symmetric matrix. Due to the circular and reciprocal symmetry, for 8 computational units (wedges) the resulting vectors are 5-dimensional. For example, for EPG-EPG connectivity we obtain **c**^*EE*^ = (*c*^*EE*^_0_, *c*^*EE*^_1_, *c*^*EE*^_2_, *c*^*EE*^_3_, *c*^*EE*^_4_), where *c*^*EE*^_0_ is the average strength within a unit (self-loops), *c*^*EE*^_1_ is synaptic strength between neighboring units, while *c*^*EE*^_4_ is the synaptic strength between units on opposite sides of the EB.

To convert to synaptic weights, we use four scale factors, *γ*^*ab*^, *a, b* ∈ {*E, I*} . For the fully symmetric case by default we set *γ*^*II*^ = 0 unless otherwise stated, as its range was very narrow ([−3.08 ×10^−4^, 0]). A given connectome data (from the male CNS data or from the sampled connectomes), typically of the form 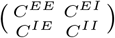, is constructed from the corresponding 5-dimensional synaptic count vectors **c**^*EE*^, **c**^*EI*^, **c**^*IE*^ and **c**^*II*^ . An effective inhibitory feedback matrix can also be computed as *C*^*EIE*^ = *C*^*EI*^ (*I* − *γ*^*II*^ *C*^*II*^)^−1^ *C*^*IE*^, and the corresponding count vector **c**^*EIE*^ from the first column. From the excitatory **c**^*EE*^ and inhibitory feedback **c**^*EIE*^ synaptic count vectors a 3× 2 effective connectome matrix is constructed: 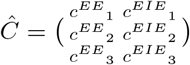, by subtracting out the self-loops. A linear combination of the two column vectors of *Ĉ* should yield the effective weights **w** = (*w*_1_, *w*_2_, *w*_3_), thus a two-dimensional mapping parameter vector 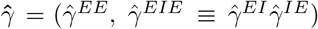 from the effective connectome matrix to the effective weights: 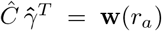 would yield the connectome-derived ring attractor, conditioned on the effective weights satisfying the analytically derived conditions (cited in Table 1, Appendix), expressed as a function of *r*_*a*_. Here 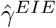 is a lumped parameter. This translates into finding the value *r*_*a*_ in the range [0, 1] that minimizes || (*Ĉ Ĉ*^†^ − *I*) **w**(*r*_*a*_)||_2_, which was computed numerically with precision of 10^−10^. The correct *γ* values were then calculated using 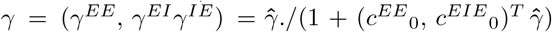, accounting for the normalization step due to the subtraction of the *w*_0_ self-loop weights. The lumped *γ*^*EIE*^ parameter was further distributed to 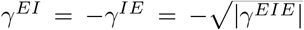, to account for the correct signs. The final weights were then tested to check if they satisfy the inequalities cited in Table 2 (Appendix; whether the presumed active and inactive EPG neurons have the correct signs and whether *w*_4_ is below its upper bound), and *γ* values were tested for *γ*^*EE*^ > 0, *γ*^*EI*^ < 0 and 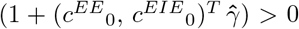. Furthermore, for the cases where *γ*^*II*^ < 0 (as in Fig. 4e) the correct signs of the active and inactive Δ7 neurons were also checked in steady state for *µ* = 1 and *σ* = 1 (G, Appendix).

**Table 1.** References to the equations that define the ring attractor conditions.

| Constraint | Symmetric | Mirror-symmetric |
| --- | --- | --- |
| Continuous encoding | (A.10) & (A.13) | (B.44) & (B.48) |
| Smooth transition | None | (B.54) |
| Stability | (A.21) | (B.50) |
| Profile requirements | (A.26) | (B.55) |

**Table 2.** References to the inequalities that limit the parameter space of valid solutions.

| Constraint | Symmetric | Mirror-symmetric |
| --- | --- | --- |
| Inactive EPG states | (A.35) | (B.63) |
| Active $z$ states | (C.82) | (C.82) |
| Inactive $z$ states | (C.81) | (C.81) |

In the mirror-symmetric case, there are effectively three *γ* parameters that would map the connectome data to the effective weights. There are also three equations that the effective weights should satisfy for generating a ring attractor (Table 1, Appendix), thus solving a system of nonlinear equations would yield the fixed-point solutions. For this, we estimated the residuals of the three equations in a 100 × 100 × 100 grid of the parameter space (bounds: *γ*^*EE*^ ∈ [0, 0.05], *γ*^*EIE*^∈ [−0.001, 0] and *γ*^*II*^ ∈ [−0.05, 0]) to find clusters of candidate points that have zero crossing for all three parameter values. The minimal residual point of each such cluster was selected as an initial condition for solving the system. For this, we used Matlab’s *fsolve* solver with the Levenberg-Marquardt algorithm. For each candidate *γ* vector the effective matrix was computed as

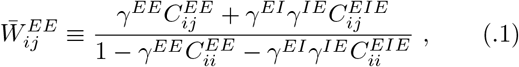

for all *i ≠ j*, and the diagonal terms were set to zero, 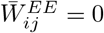, see Appendix C for details. From the effective matrix the 6 effective weights are extracted and left-hand sides of the equations in Table 1 (Appendix) calculated. Through iterative procedure the sum of squares of the left-hand side values is minimized by *fsolve* to calculate the fixed point. Similarly with the fully-symmetric case, the corresponding additional inequalities are checked to filter out invalid solutions.

For both, the fully-symmetric and the mirror-symmetric ring attractors the marginal stability of the system was verified numerically by perturbing the continuum states with additive noise of 10% and 50% from the maximal state component and checking whether the system converges back to the continuum states using the *ode23* solver in Matlab. This happened 100% of the time for the three computed connectome-derived ring attractors.

### Generating random samples around the connectomic matrices

To generate a random connectivity matrix sample with a given synaptic count % variation around the male CNS connectomic data (as in Fig. 5c), we generated uniformly distributed random samples around each value of the **c**^*EE*^, **c**^*EI*^, **c**^*IE*^ and **c**^*II*^ vectors, in the range 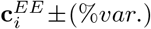, with *i* being the vector index and %*var*. ∈ [0, 100]. We then compiled the samples into a matrix form. The resulting network was then tested to ascertain whether when combined with the *γ* scale factors it fulfills the stability and continuum conditions, as well as whether the assumed active states of EPG and Δ7 neurons were correct. Value of *γ*^*II*^ = 0 was used for the fully-symmetric case. 1000 samples per % variation were drawn for each case.

### Comprehensive sampling of all possible connectivity matrices

For the fully-symmetric case we also have fully-symmetric excitatory and inhibitory feedback matrices. Given that the self-loops (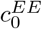 and 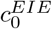) in the network can be subtracted out (Appendix) and the synaptic weights towards the opposite side of the network (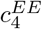 and 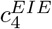) only influence inequalities and impose relatively weak constraints, we end up with 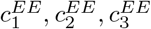 and 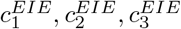 as parameters for sampling connectivity matrices. Further, as the *γ* scaling factors introduce arbitrary scaling, by introducing normalized synapse counts, 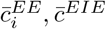, that satisfied 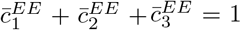 and 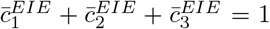, we end up comprehensively sampling the whole space of possible connectivity matrices with 4 free parameters. For each sample we then estimate whether it can be mapped to a successful continuous ring attractor using the *γ* scale factors. In Fig. 5d we show the projection of the 4-dimensional space into the excitatory matrix subspace (top) or inhibitory matrix subspace (bottom), averaging over the remaining two dimensions to calculate the success fraction. In both cases we identify the same main axis along which the variation of success rate is the highest - the median of the traingle (*c*^*E*^ and *c*^*I*^). We use these vectors to compile a two-dimensional projection plot (Fig. 5e) showing the success fraction variation as a function of the combination of excitatory (right) and effective inhibitory profiles (top).

For a more general approach, as shown in Supplementary Fig. S4, we uniformly sampled each component of the four 5D vectors **c**^*EE*^, **c**^*EI*^, **c**^*IE*^ and **c**^*II*^, for a total of 10^6^ parameter sets. Each 5D vector was amplitude-matched to the respective vector from the Male CNS dataset. Each parameter set was tested for a solution for each of the three cases (*A*_Δ_ ∈ {8, 6, 4}). Following that, we performed a quadratic discriminant analysis (QDA) to find two primary discriminant axes, one for the excitatory vector and another one for the combination of inhibitory feedback vectors, that best separate the successful points from the unsuccessful ones [80]. QDA was used as the variance of the successful set of points is much smaller than the one of the unsuccessful one, and to account for potential non-linear parameter dependencies and boundaries between the two classes. We projected the points on the 2D space defined by those two discriminant axes, and colored them by their success value. 2D histograms were computed over the projected scatter points, capturing the success fraction per bin. The projections from the four connectome datasets were also plotted on the 2D space.

### Generative AI

We would like to acknowledge the use of OpenAI’s ChatGPT for assisting with language and grammar refinement.

## Acknowledgments

We would like to thank Vivek Jayaraman, Daniel Turner-Evans, Marcella Noorman, and Ann M. Hermundstad for insightful discussions. We are grateful to Vivek Jayaraman and Brad K. Hulse for experimental support. This work was supported by the Howard Hughes Medical Institute and the Janelia Visiting Scientist Program. JEF acknowledges support from the National Institute for Theory and Mathematics in Biology through the National Science Foundation (grant number DMS-2235451) and the Simons Foundation (grant number MPTMPS-00005320).

---

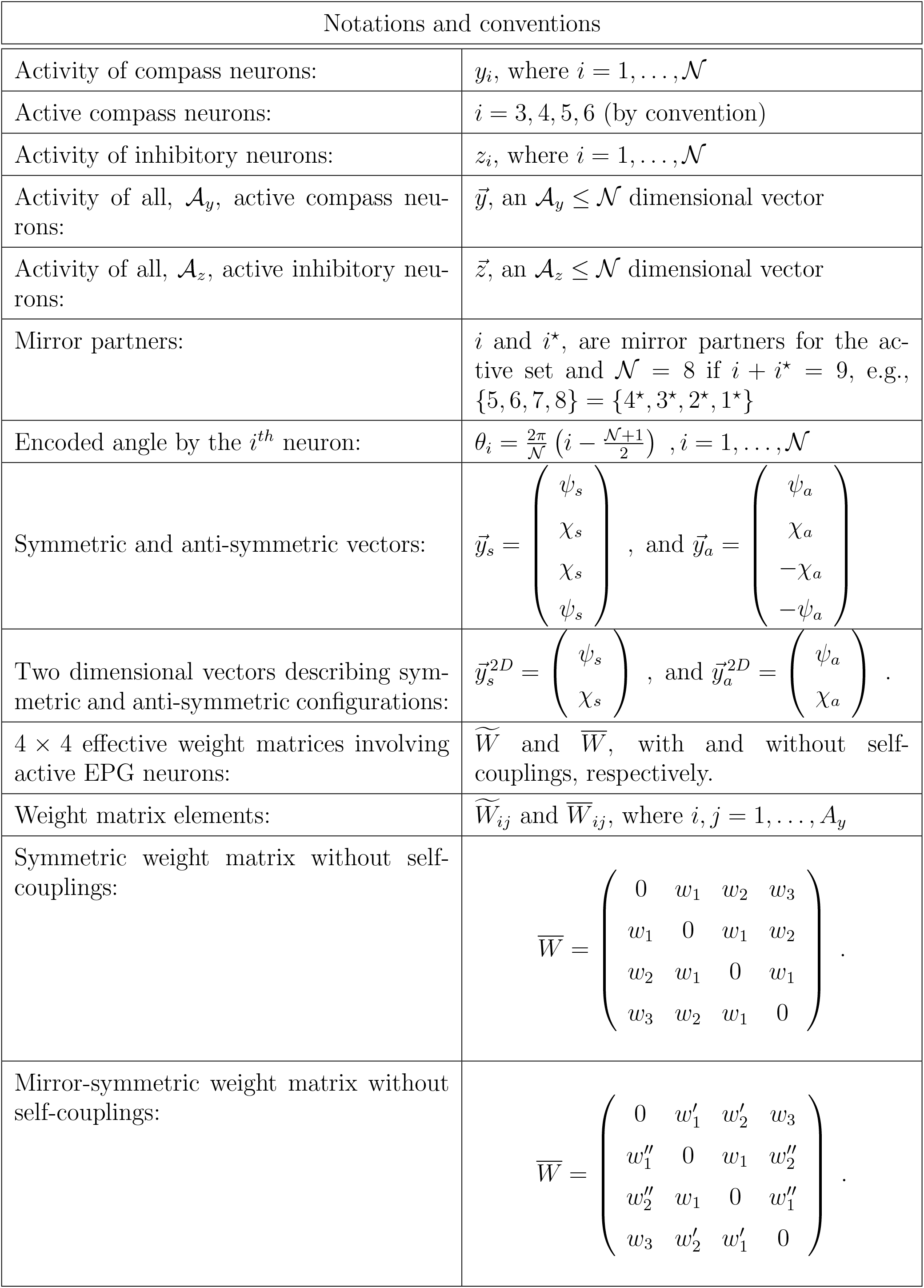

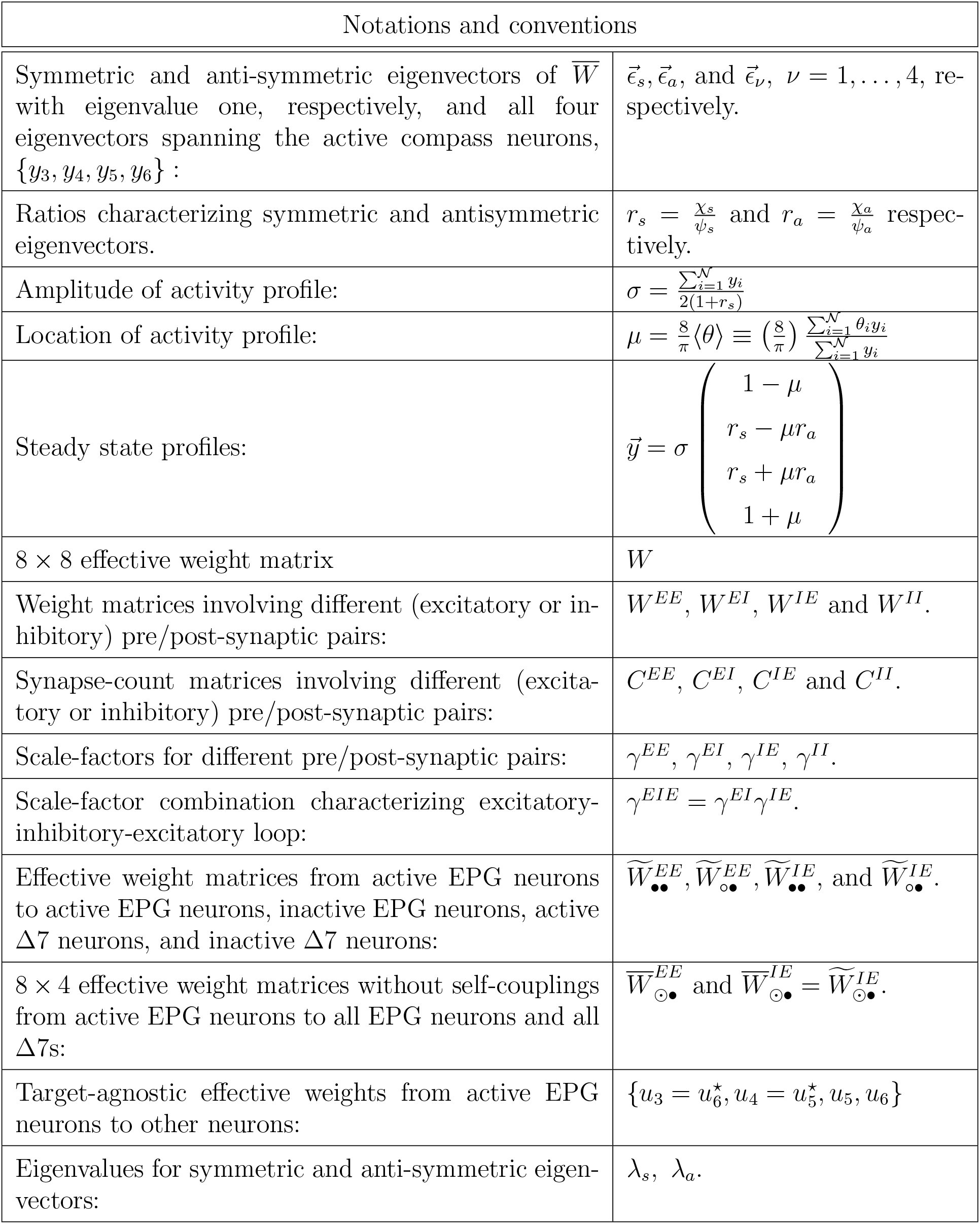

**Supplementary Figure S1.**
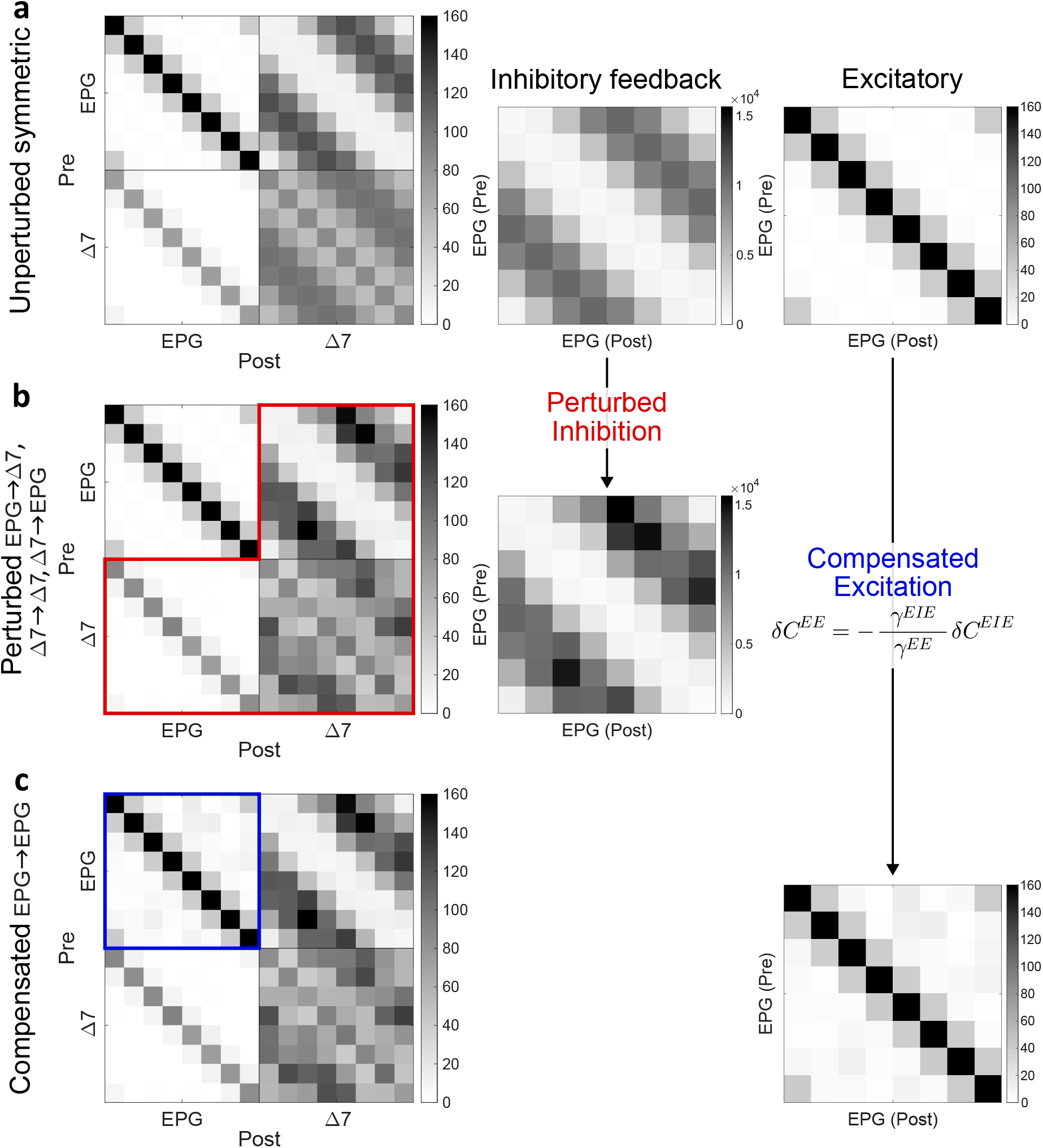
An exact ring attractor from asymmetric connectome. (a) (left) Coarse-grained and symmetrized fly EPG-Δ7 connectome, same as Fig. 4b; (middle) the product *C*^*EIE*^ = *C*^*EI*^(*I* −*γ*^*II*^*C*^*II*^)^−1*C*^^*IE*^ characterizing the indirect inhibitory loop for symmetric ring attractors with *γ*^*II*^ = −1.54 × 10^−4^; and (right) *C*^*EE*^ representing the direct connections between the EPG neurons. (b) (left) Perturbed inhibition where we have added 30% perturbations to the *C*^*EI*^, *C*^*II*^ and *C*^*IE*^ synapse count matrices; (middle) the resulting perturbed inhibitory loop matrix, *C*^*EIE*^. (c) (right) Excitation, *C*^*EE*^, to compensate the perturbations in the inhibitory loop and maintain an exact ring attractor; (left) the resulting asymmetric EPG-Δ7 connectome that can realize a symmetric continuous ring attractor.

**Supplementary Figure S2.**
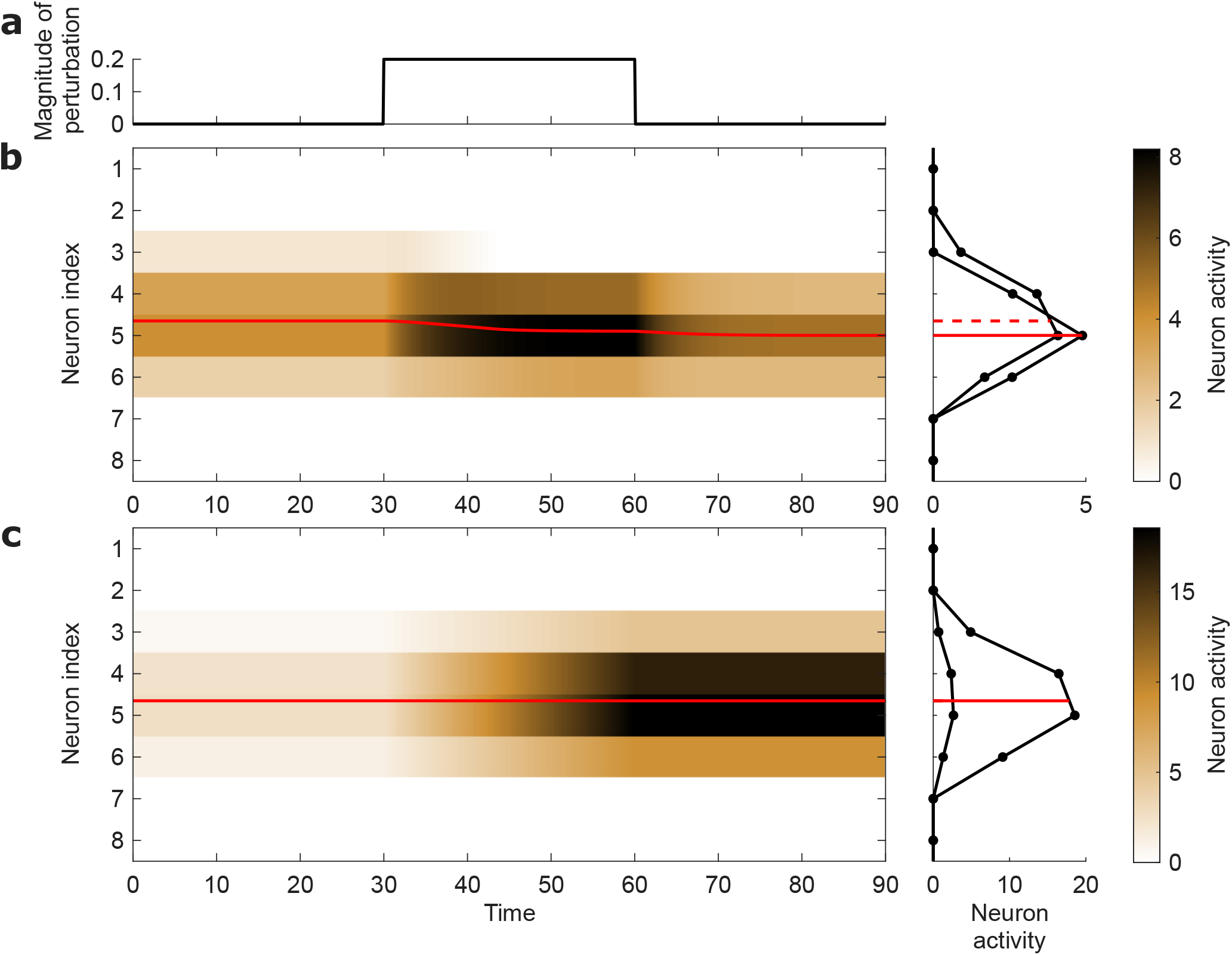
Potential decoding problems in driven ring attractors. (a) Depicts the amplitude of an external input that is proportional to the (initial) steady state activity profile. In our simulation we initially set the amplitude to be zero (no external input). We turn on the input at a certain time and switch it off at a later time. (b) While driven ring attractor networks can sustain steady state solutions corresponding to a continuum of angles, the angular representation (red curve) drifts in the presence of the external input (left). Even after the external input is removed, the encoded angle do not return to its initial value (right). (c) An equivalent simulation for the self-sustaining network shows no angular drift.

**Supplementary Figure S3.**
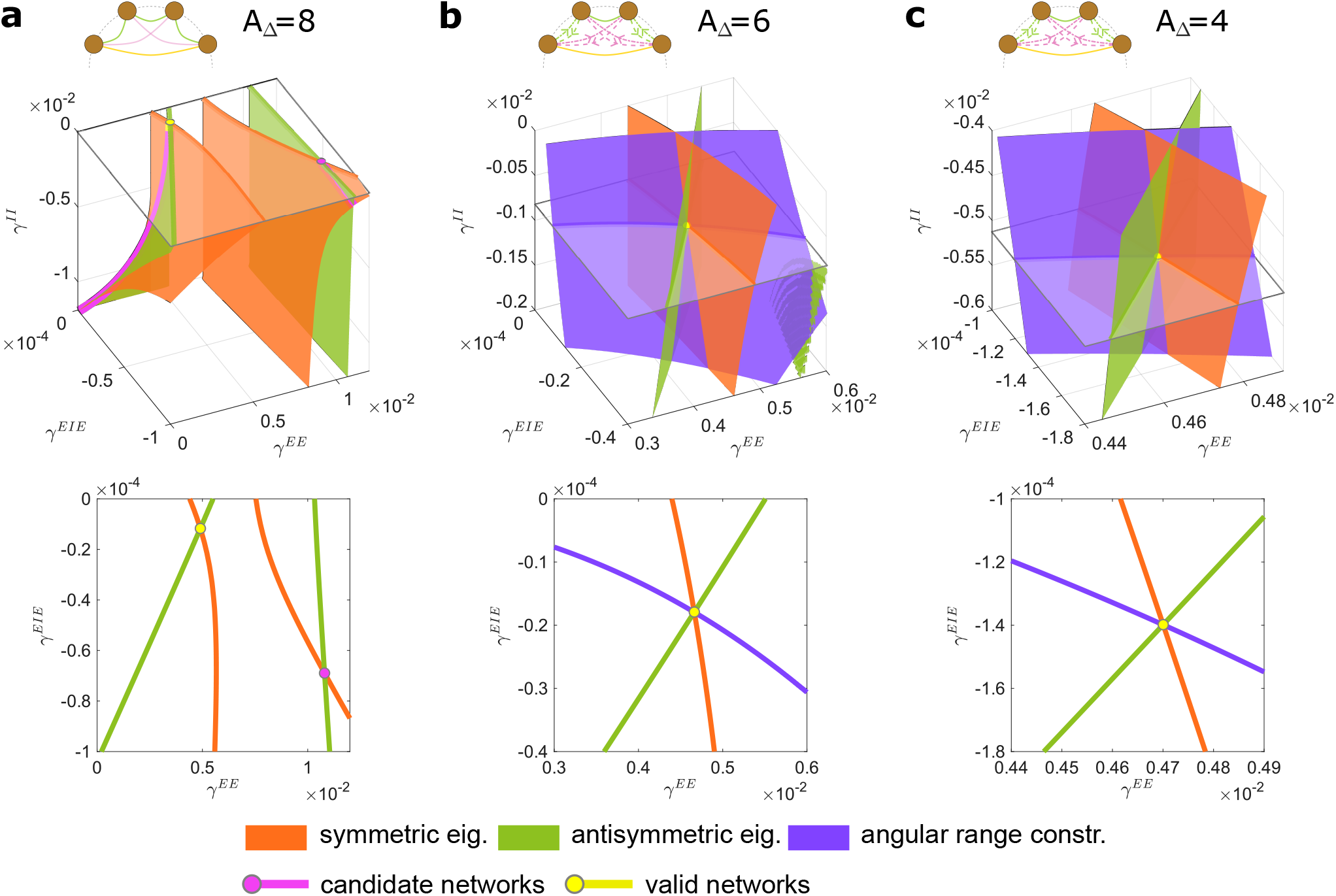
Constraints relating the connectome to a continuous ring attractor. (a, top) *γ*-parameter view of the connectome-constrained symmetric ring attractor realization in Figs. 4d and 4e. The orange and green surfaces represent where the two symmetric and anti-symmetric eigenvalue conditions are satisfied respectively, and the pink curves their intersections. Various inequality conditions further restrict the space of viable continuous ring attractors to the yellow curve where 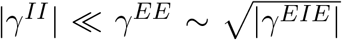. (a, bottom) 2D slice of the *γ*-parameter space where *γ*^*II*^ = 0. (b and c, top) Analogous plots for the mirror-symmetric realizations of connectome-constrained ring attractors for 𝒜_Δ_ = 6 and 𝒜_Δ_ = 4, respectively, showing additionally the angular-range constraint (purple surfaces) that the mirror-symmetric networks have to satisfy. In each of the plots, the three surfaces intersect at a unique point (yellow dot) which also satisfies all the inequalities to make it a valid continuous ring attractor. (b and c, bottom) 2D slices at the *γ*^*II*^ values of the valid solutions.

**Supplementary Figure S4.**
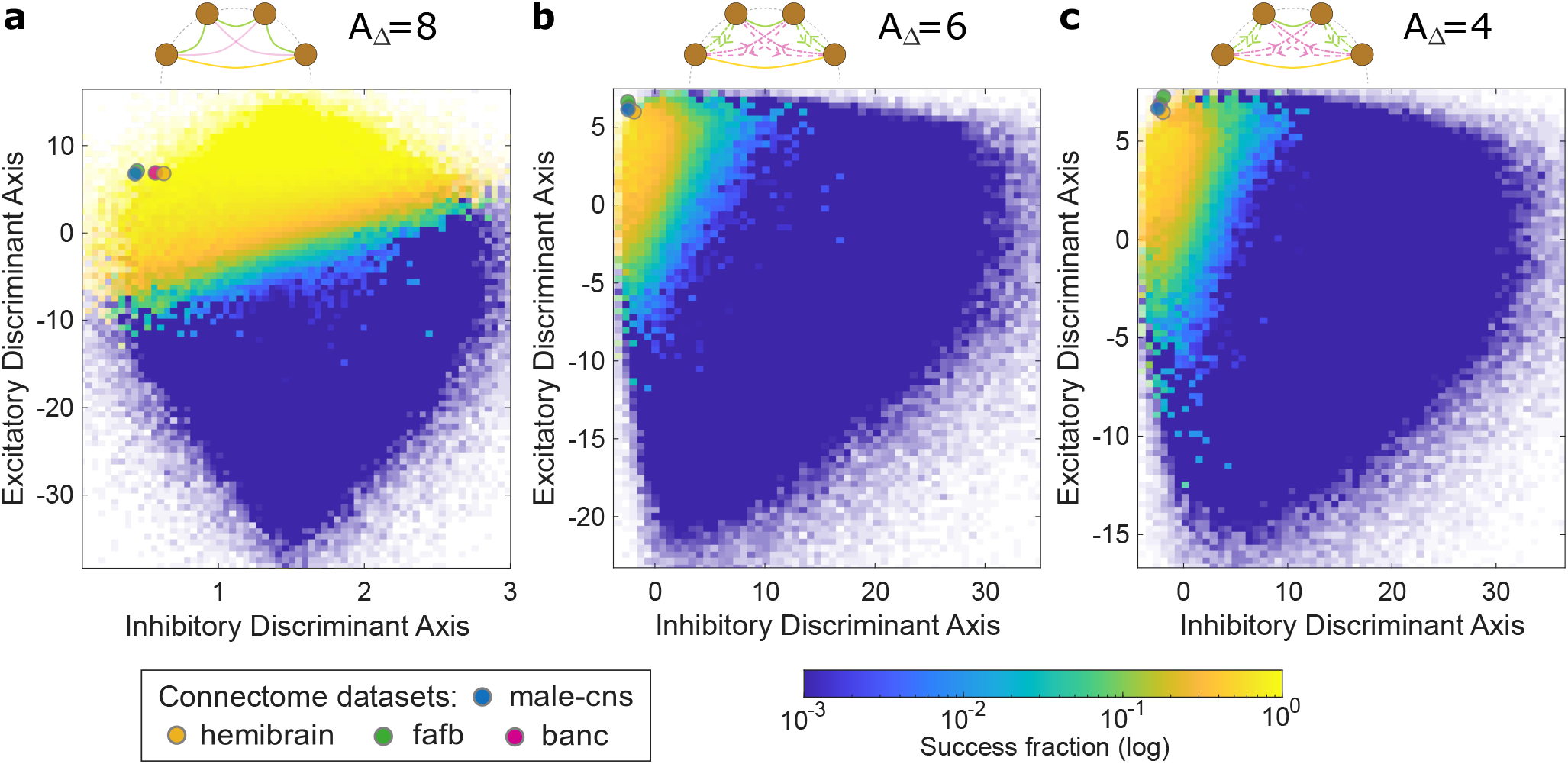
Success fraction along the primary excitatory and inhibitory discriminant axes. The average success fraction is shown for the cases for *A*_Δ_ = 8 (a), *A*_Δ_ = 6 (b), and *A*_Δ_ = 4 (c) as a projection on the 2D space made from the primary excitatory and inhibitory discriminant axes, analogous to *c*^*E*^ and *c*^*I*^ in Fig. 5f. 10^6^ 20-dimensional samples were drawn, where each component of **c**^*EE*^, **c**^*EI*^, **c**^*IE*^ and **c**^*II*^ was uniformly sampled, and the success of the found solutions was assessed. The primary discriminant axes were derived using quadratic discriminant analysis (Methods). 2D histograms were computed over the projected scatter points, capturing the success fraction per bin.

**Supplementary Figure S5.**
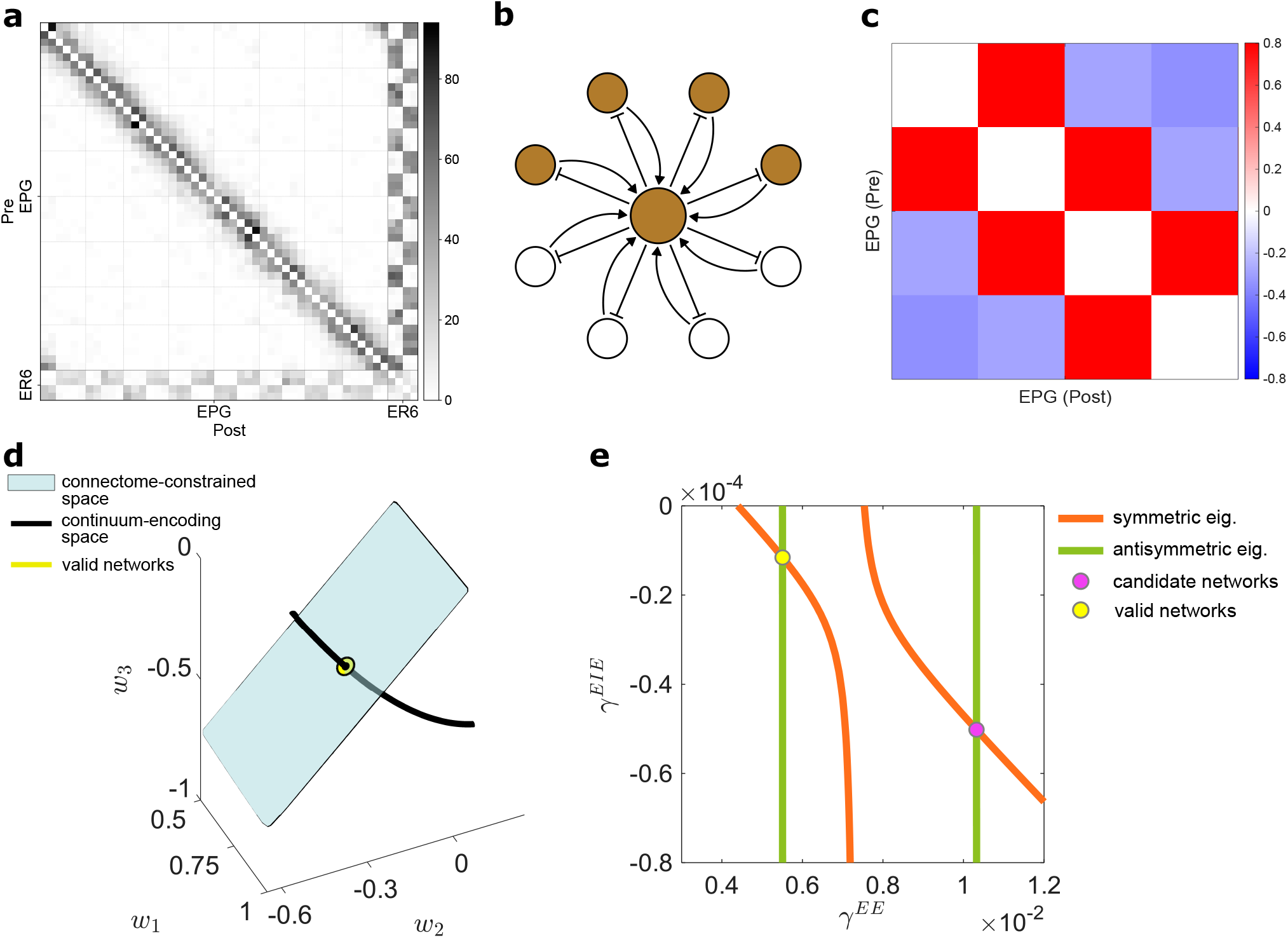
Constructing ring attractors from the EPG-ER6 connectome. (a) EPG and ER6 neurons link between each other in the gall and ellipsoid body. Synaptic count matrix from the Male CNS dataset is shown that illustrates how uniformly the ER6 population connects to and from the eight EPG computational units. (b) Two-population network model (outer layer - EPG, inner circle - ER6), wherein EPG neurons receive uniform, rather than angularly tuned, inhibition. (c) Effective connectivity between the active EPG neurons corresponding to the successful connectome-constrained ring attractor model, analogous to Fig. 4d. Red - excitatory, blue - inhibitory weights. (d) The intersection point (yellow) between the theoretical conditions of the continuum-encoding space (black curve) and connectome-constrained space (blue plane) yields a unique candidate weight profile of a connectome-constrained ring attractor, analogous to Fig. 4e. (e) *γ*-parameter view of the solution space, as in Supplementary Fig. S3, bottom.

## Appendix A Continuous symmetric ring attractor networks

In this appendix, we analyze when a recurrent network with discrete rotational symmetry can support a *continuous* ring attractor with a continuous family of steady-state activity profiles. We focus on a threshold-linear network with 𝒩 neurons, all-to-all connectivity, and no self-couplings. This idealized formulation is designed to capture the essential structure underlying the fruit fly’s head-direction system while allowing for analytic characterization.

### A.1 The network

Consider a recurrent threshold-linear network of an even number 𝒩 of neurons with activities *y*_*i*_ (*i* = 1, …, 𝒩). We label neurons such that each one encodes a preferred angle

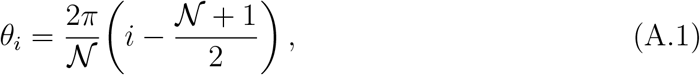

ensuring that preferred directions are evenly distributed and centered around zero. For example, when 𝒩 = 8, the encoded angles are *±π*/8, *±*3*π*/8, *±*5*π*/8, and *±*7*π*/8.

The firing rates evolve according to

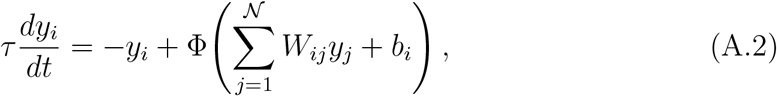

where *τ* is the neuronal time constant (assumed to be the same for all the neurons in the network), *W*_*ij*_ denotes the synaptic weight from neuron *j* to *i, b*_*i*_ is an external bias, and Φ(*x*) = max(0, *x*) is the threshold-linear nonlinearity. We focus on *self-sustained* attractors and therefore set *b*_*i*_ = 0; the nonzero-bias case is treated in Appendix F.

Throughout this appendix we impose:

I. **Rotational symmetry:** *W*_*i*+1,*j*+1_ = *W*_*ij*_,
II. **Reciprocal symmetry:** *W*_*ij*_ = *W*_*ji*_,
III. **Zero self-coupling:** *W*_*ii*_ = 0,

the last of which entails no loss of generality (Appendix D).

Steady states satisfy

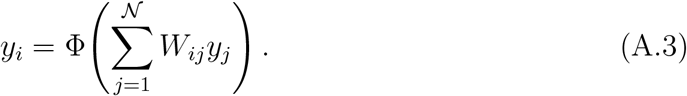

A crucial observation (discussed in the main text) is that for a network to support a *angular continuum* of self-sustained activity bumps, the 𝒩_*y*_ × 𝒩_*y*_ submatrix 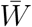 acting on the active neurons must be *doubly degenerate*, or equivalently, 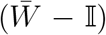 must have two zero eigenvalues. To see this, let us label the active neurons with indices 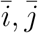. Since active neurons receive nonnegative input, the threshold is inactive and their activities must satisfy the linear system

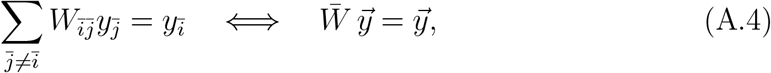

so 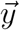 is an eigenvector of 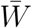 with eigenvalue 1.

For convenience we consider networks where both 𝒩 and 𝒜_*y*_ are even. We define the “mirror partner” of neuron *i* by *i*^⋆^ satisfying *i* + *i*^⋆^ = 𝒩 + 1, which implies the symmetry

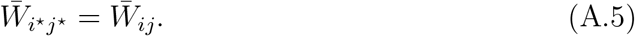

Appendix E shows that if 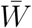 has only *one* eigenvector with eigenvalue 1, then that eigenvector must be either (i) symmetric, 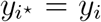, or (ii) antisymmetric, 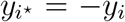. Antisymmetric steady states are impossible in threshold-linear networks (they require negative activities), so only symmetric solutions remain. Such profiles will have an average angular value,

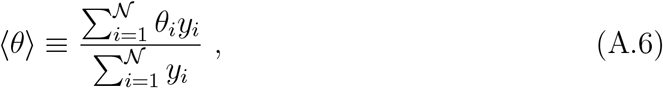

that only encodes the *central* angle of the active set. They therefore cannot represent the head direction continuously. This demonstrates that a self-sustaining ring attractor requires at least a two-dimensional eigenspace with eigenvalue 1.

Note that if 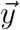 is an eigenvector with eigenvalue 1, then by symmetry the vector 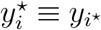 is also an eigenvector. Thus if 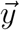 is neither symmetric nor anti-symmetric, then the two together span the required two-dimensional space of steady-state solutions. Different linear combinations of them correspond to different encoded angles.

### A.2 Eigenvalue constraints on the weights

In Appendix E, we will show that the two-dimensional eigenspace can also be described as a vector space spanned by a symmetric and an anti-symmetric eigenvector. Thus to obtain all the steady-state activity profiles, we will find the symmetric and antisymmetric eigenvectors separately and obtain the constraints they impose on the weight space along the way.

For concreteness we specialize to the biologically relevant case of 𝒩 = 8 and 𝒜_*y*_ = 4, while noting that the same approach generalizes. Let the active neurons be *i* = 3, 4, 5, 6. A general symmetric eigenvector can be written as

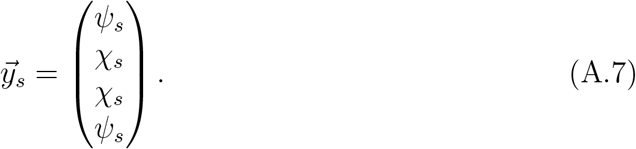

Rotational and mirror symmetry constrain the 4 × 4 matrix 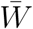 to take the form

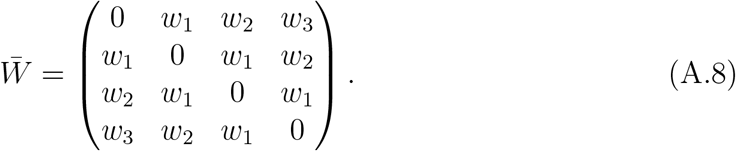

Substituting (A.7) into 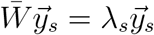 yields a reduced eigenvalue problem:

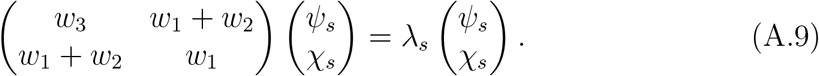

Setting *λ*_*s*_ = 1 leads to the first constraint on the weights (Appendix Fig. 1):

**Appendix Fig. 1.**
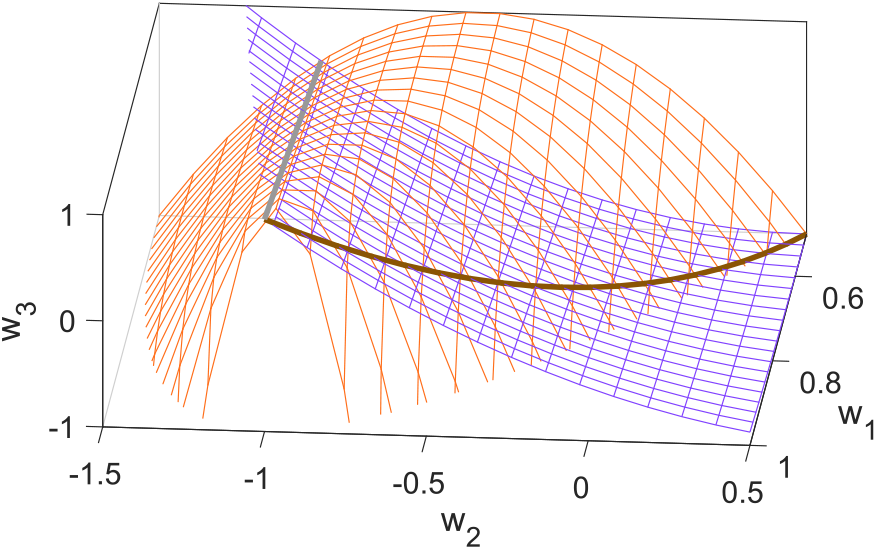
Weight manifold supporting continuous angular encoding. Intersection points (brown) between the weight submanifolds satisfying the symmetric conditions (scarlet) and the antisymmetric conditions (violet) yield valid ring attractors. The grey intersection points lead to unphysical activity profiles.

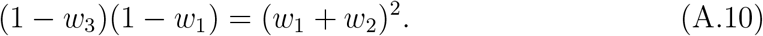

Stability requires that the eigenvalue of the *other* symmetric eigenvector be less than 1, which is equivalent to

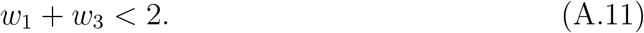

The two eigenvalues are explicitly

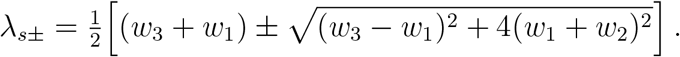

The antisymmetric eigenvectors take the form

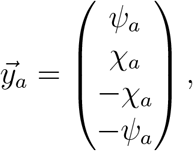

and obey the reduced eigenvalue problem

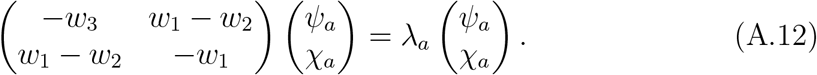

Imposing *λ*_*a*+_ = 1 gives a second constraint:

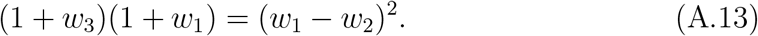

Stability additionally requires

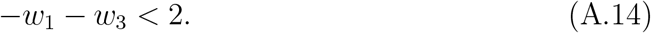

Combining (A.11) and (A.14) yields

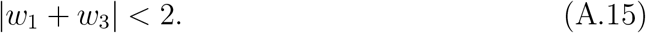

We can express *w*_3_ from both (A.10) and (A.13) which results in

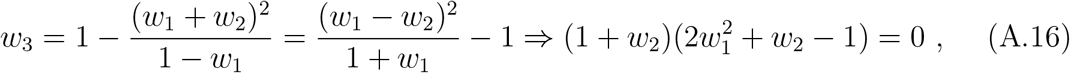

after some straightforward algebra. The solution with *w*_2_ = −1 is incompatible with nonnegative activity profiles (see below), leaving

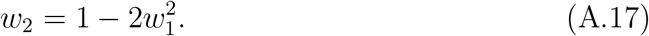

Substituting into either eigenvalue constraint yields

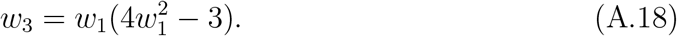

The stability condition (A.15) reduces to

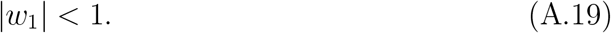

We can see this by substituting *w*_3_ in the inequalities (A.15) to obtain,

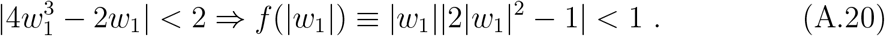

We note that while |*w*_1_| is a monotonically increasing function spanning the interval [0, 1], the second term, |2|*w*_1_|^2^ − 1|, has a minimum at 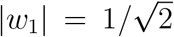 and reaches the value of one at |*w*_1_| = 0, 1. Thus within the interval [0, 1), the product is less than one. At |*w*_1_| = 1 the product is one, and beyond that (|*w*_1_| *>* 1) both terms monotonically increase and therefore the product is greater than one violating (A.20). In other words, (A.20) is satisfied if and only if

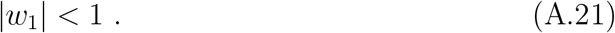

### A.3 Constraints from requiring single-bump activity profiles

The steady-state activity profile is a linear combination of the symmetric and antisymmetric eigenvectors. Letting *σ* = *ψ*_*s*_ and writing *ψ*_*a*_ = −*µσ*, we obtain

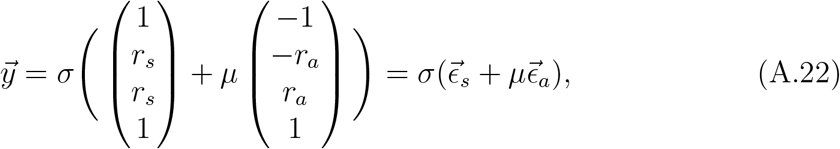

Where

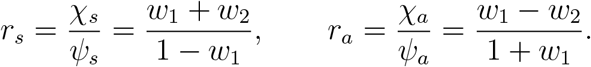

Using the viable solution branch (A.17), (A.18), these simplify to

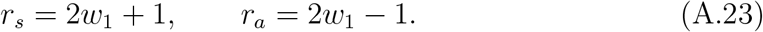

Hence

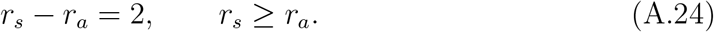

It is instructive to compute ⟨*θ*⟩. Using our conventions, for the active set, *i* = 3, 4, 5, 6, encoding angles 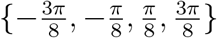, respectively, we have

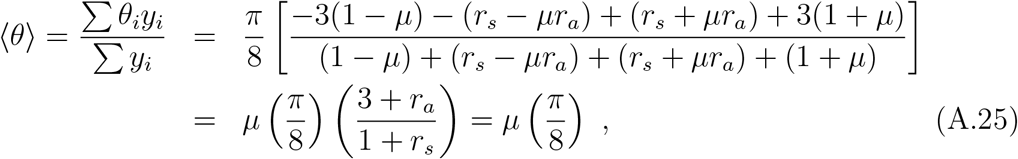

where *y*_*i*_ values were substituted from (A.22), and in the last step we used the relation between the ratios (A.24). Thus, as the location parameter, *µ*, varies in the interval, [−1, 1], the attractor covers exactly the *π*/4 range needed so that eight active sets tile the full 2*π* space.

Since *r*_*a*_ affects the shape of all the activity profiles (A.22) in a given ring attractor network, we will refer to it as the shape parameter. For instance, to obtain a single-bump profile, we require *r*_*a*_ ≥ 0, which prevents alternating peaks and troughs by guaranteeing that 1 − *µ* ≤ *r*_*s*_ − *µr*_*a*_ for all *µ* ∈ [−1, 1]. This yields

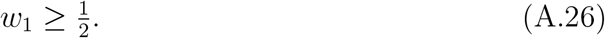

Together with (A.21), we obtain

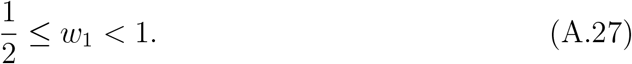

### A.4 Inequality constraints from inactive neurons

Finally, let us look at the constraints needed to ensure that the total input drive to the inactive neurons is non-positive. We need to check consistency for all the steady-state profiles parametrized by *µ* and *σ*. Due to the symmetry properties we only need to consider drives, *d*_*i*_, onto the neurons, *i* = 7, 8.

The inactivity of *y*_7_ implies

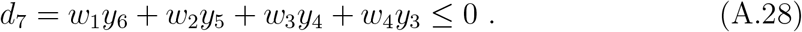

Substituting the activity profile (A.22), one can rewrite the above inequality as

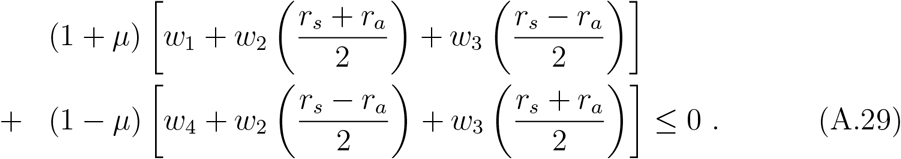

We see that the factors multiplying (1 *±µ*) must both be negative, otherwise at either *µ* = 1 or −1, the inequality will be violated. So,

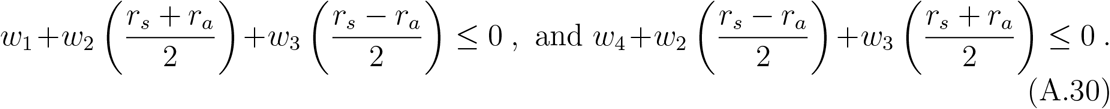

Further, since (1 *± µ*) is always positive, the above condition guarantees that the inequality will be satisfied for all values of *µ*. The inequalities simplify once one uses the expressions for *r*_*s*_, *r*_*a*_:

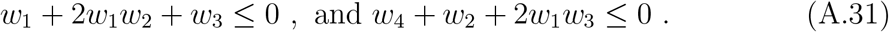

It is easy to check using (A.17, A.18) that the left hand side of the first inequality is identically zero and therefore the inequality doesn’t lead to any new constraints. The second inequality can be re-expressed as a bound on *w*_4_ in terms of *w*_1_:

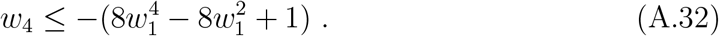

One can perform calculations that are very similar to above to derive constraints from inactivity of *y*_8_. First,

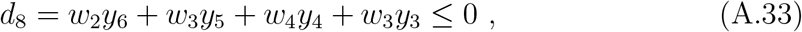

leads to the inequality

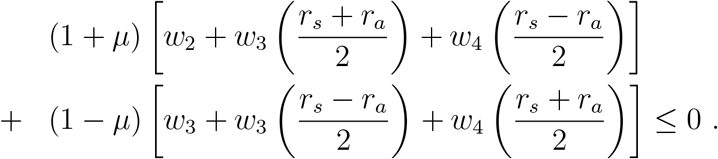

This inequality is satisfied if and only if,

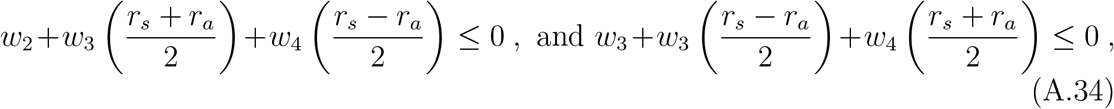

which can be simplified to yield

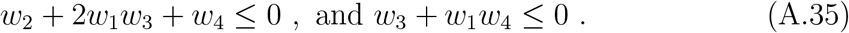

The first inequality is the same as was obtained earlier, while the second is a new constraint that gives rise to another bound for *w*_4_:

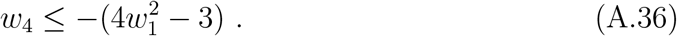

### A.5 Concluding remarks

To summarize, the ensemble of self-sustaining continuous ring attractors obeying the symmetries assumed is characterized by two parameters, *w*_1_ and *w*_4_, where the latter must satisfy the bounds (A.32, A.36), *w*_1_ must lie in the range (A.27), and *w*_2_, *w*_3_ are given by (A.17, A.18). It is interesting to note that one could parametrize the weights in terms of an anlge, *ϕ* according to

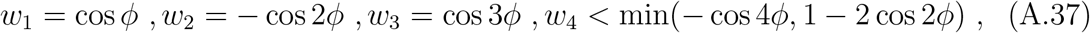

## APPENDIX B B Mirror-symmetric ring attractor networks

In this section we generalize the analysis above to effective weight matrices with reduced symmetry properties that arise naturally from the connectomic features present in fruit flies.

### B.1 Eigenvalue constraints on the weights

As we will discuss in Appendix E, once the compass neurons whose activity encodes angular locations can drive each other indirectly via other neuronal populations, the effective connectivity between the compass neurons need not obey all the symmetries assumed in the previous appendix. For instance, even if the entire connectivity structure preserves circular symmetry, the effective connectivity between the compass neurons need not preserve this symmetry due to the presence of the threshold non-linearity. In Appendix E, we will argue that for the fruit fly head-direction system a reduced *mirror-symmetric* effective weight matrix defined via

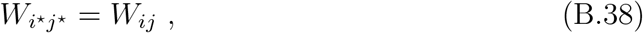

is what is appropriate to consider.

For the active set we are considering (*i* = 3, 4, 5, 6), the mirror-symmetric weight matrix is parameterized by six weight parameters, 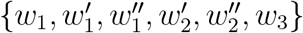:

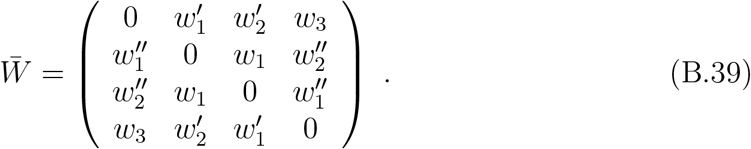

The arguments of the previous section, Appendix A, on the eigenvalue requirements of the active weight submatrix go through for the mirror-symmetric configuration as well, and therefore 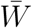 must still have two degenerate eigenvectors with eigenvalues of one. As before, we need one of the eigenvectors to be symmetric and the other to be anti-symmetric. Substituting the symmetric eigenvector (A.7) in the eigenvalue equation (A.4), we again obtain a reduced eigenvalue equation:

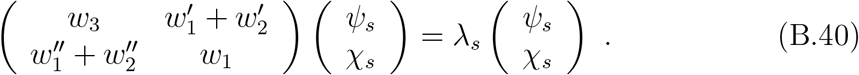

The eigenvalues of a 2 × 2 matrix have closed form expressions:

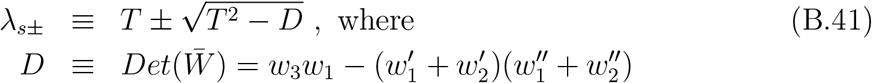

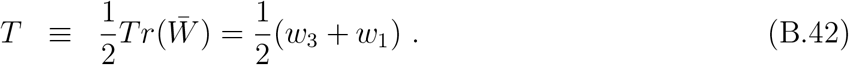

In order to have a stable attractor then, we must have

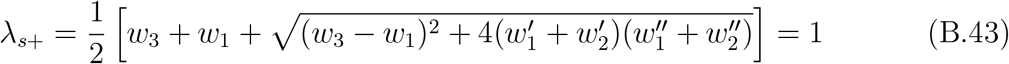

The above equation is equivalent to requiring

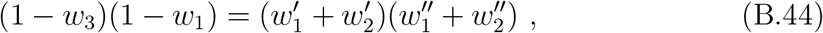

and just like in the symmetric network (A.11), 2*T* = *w*_1_ + *w*_3_ *<* 2.

Let us next determine the degenerate antisymmetric eigenvector,

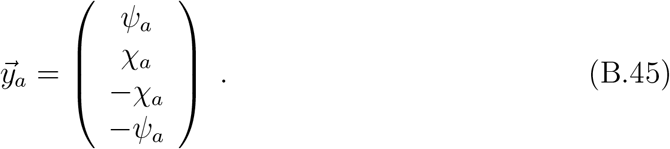

As with the symmetric case, the eigenvalue problem reduces to a two-dimensional problem:

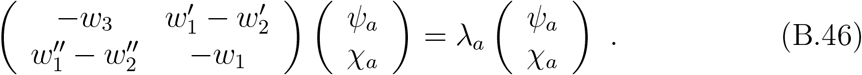

The eigenvalues now read

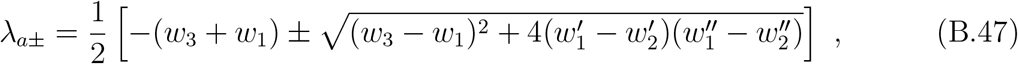

or equivalently, the weights must satisfy the relation

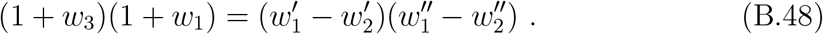

The pertinent inequality, (A.14), becomes *w*_1_ + *w*_3_ *>* −2.

To summarize, to have a stable continuous ring attractor we must satisfy

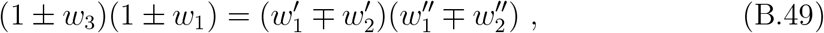

and

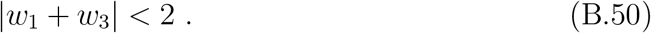

We point out that these equations reproduce the symmetric model if we set 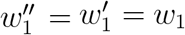 and 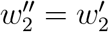.

### B.2 Constraints from neuronal activity

Exactly as in the symmetric model, the existence of two degenerate eigenvectors ensures that we have a two-dimensional space of attractor profiles (A.22). The ratios *r*_*s*_ and *r*_*a*_ can be calculated similarly to the symmetric case as well:

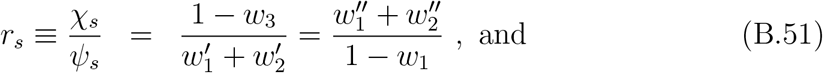

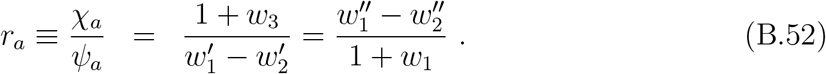

To ensure positivity of all the responses^1^, we must satisfy

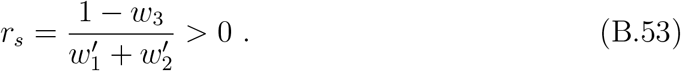

If we further demand that *µ* = 1 should correspond to a symmetric configuration involving *i* = 4, 5, 6 that can then smoothly transition to the next set of active set, *i* = 4, 5, 6, 7, then we must have *y*_4_ = *y*_6_ at *µ* = 1:

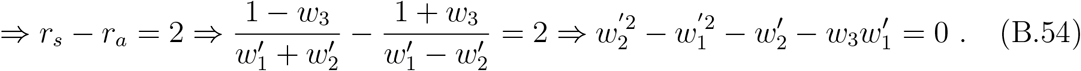

Finally, requiring the bump has a single maximum implies

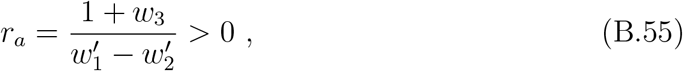

which immediately ensures the previous inequality (B.53) by virtue of (B.54).

### B.3 Solution Space

To summarize, we have seen that a mirror-symmetric weight matrix has six independent weight parameters characterizing the weight matrix involving the active set. There are however three equations, (B.49, B.54), that these parameters must satisfy to produce a smooth continuous ring attractor. Thus we expect a three-dimensional solution space of ring attractors. We will now provide a convenient parametrization of the solution space that we frequently use for analysis and results.

The three parameters we will choose to parametrize the solutions are *w*_1_, *w*_3_, and *r*_*a*_. We note that the weights *w*_1_ and *w*_3_ are the only two weights that are symmetric, while *r*_*a*_ enables us to characterize the activity profiles in a uniform way that is valid for both the symmetric and mirror-symmetric models. Using the definitions of *r*_*a*_ and *r*_*s*_ = *r*_*a*_ + 2, one can straightforwardly obtain 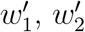 in terms of *w*_3_ and *r*_*a*_:

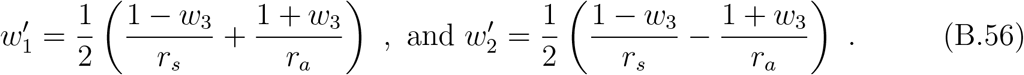

The constraints from the eigenvalue conditions can then be used to express 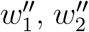 in terms of *w*_1_ and *r*_*a*_:

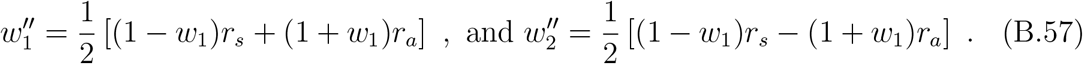

The ranges of the three parameters are restricted as

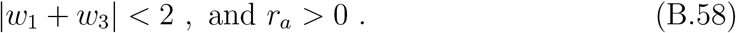

It is useful to point out a couple of key differences between the mirror-symmetric and the symmetric case. Unlike the symmetric case, here *w*_1_ and *w*_3_ are free parameters, and thus the allowed range of *w*_1_ can vary depending upon the value of *w*_3_:

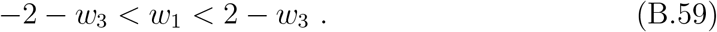

*w*_1_ and *r*_*a*_ are also independent in the general mirror-symmetric ring attractor unlike the symmetric case. Thus while *r*_*a*_ inherited a maximum bound of one from the bound of *w*_1_, this is no longer true for mirror-symmetric ring attractors

### B.4 Constraints from inactive neurons

Finally, we consider the constraints on the weights from the active to the inactive neurons, since the net drive to each inactive neuron must be negative for all activity patterns. Note that the synapses onto inactive neurons are now completely independent of the recurrent weights between the active neurons. Thus, if 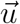 denotes the synapse vector onto any inactive neuron, we must have

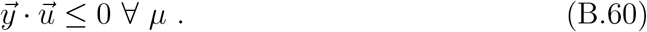

Substituting the activity profile (A.22), we obtain

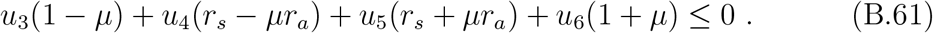

As in the symmetric case, one can re-express the left-hand side as a linear combination of two terms involving prefactors (1 *± µ*), so that the above inequality is satisfied ∀ *µ* provided the coefficients corresponding to (1 *± µ*) are negative:

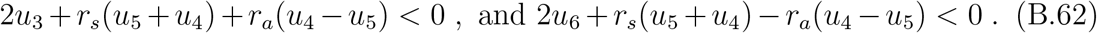

Thus, there are four independent weights that only have to satisfy two inequalities, which can be further simplified to

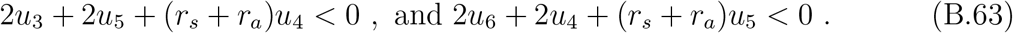

This leaves substantial flexibility: a given weight can take on a wide range of values as long as the other weights are adjusted accordingly.

While (B.63) provides an analytical inequality condition for a viable ring attractor network, there is a simple numerical way to check that these inequalities are satisfied. As argued above, in order for (B.61) to hold for all profiles, it suffices to verify that the conditions are met for the two extreme configurations, *µ* = *±*1, which can be checked numerically with ease.

## APPENDIX C C From the fly connectome to a ring attractor

In this appendix we consider a neural network that, in addition to EPG compass neurons, includes other neuronal populations that provide indirect pathways between compass neurons. We consider this scenario because, while EPG neurons can directly provide local excitatory inputs to each other, the broader, distal mutual inhibition must arise indirectly via other neuronal populations. Here we study the simplest case in which a single additional neuronal population provides indirect pathways between the EPG neurons.

This appendix is organized as follows. First, we describe how such a multi-population network can be mapped to an effective network model involving only the recurrently connected EPG compass neurons. This relies on the ability to absorb indirect pathways involving inhibitory neuronal populations into effective weights between compass neurons. Next, we derive the constraints that these effective weights must satisfy in order to realize a consistent ring attractor network. Since the effective weights are functions of the original network weights, the equalities and inequalities derived below indirectly constrain the original network. We then use measured synapse-count matrices involving the relevant neuronal populations to obtain a family of realizable networks parameterized by a few scaling parameters that convert synapse counts to weights. If there exists an intersection between the theoretical ensemble of continuous ring attractors and the realizable networks from the connectome, then we have identified a *connectome-constrained continuous ring attractor network*. Here we primarily focus on recurrent connections involving the EPG and the Δ7 neurons; at the end, we briefly explain how the analysis can be generalized to include other neuronal populations, such as ring neurons.

### C.1 Effective neuronal network that includes indirect neuronal pathways

#### C.1.1 Effective connectivity between the EPG neurons

##### Recurrent weights between active neurons

Let us consider an additional set of recurrent neurons, *z*_*i*_, *i* = 1, …, 𝒩, that provide indirect inhibitory pathways between the EPG neurons for supporting a localized bump of activity. The network dynamics are modeled as

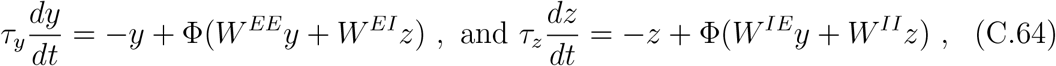

where *τ*_*z*_, *τ*_*y*_ are the neuronal time constants, which from now on will be set to one, and *W* ^*EE*^, *W* ^*EI*^, *W* ^*IE*^, *W* ^*II*^ are all 𝒩 × 𝒩 weight matrices. Now suppose 𝒜_*z*_ is the set of active *z* neurons when a given set of 𝒜_*y*_ compass neurons are active. We will assume that this set does not change with the profile parameters *µ, σ* characterizing the neuronal activity profile, except at the extreme configurations when 𝒜_*y*_ = 3. In this case, the steady-state equations for the active neuronal set are

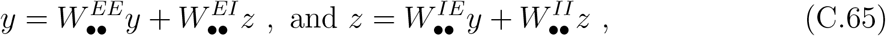

where 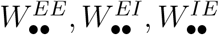 and 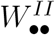 are respectively 𝒜_*y*_ × 𝒜_*y*_, 𝒜_*y*_ × 𝒜_*z*_, 𝒜_*z*_ × 𝒜_*y*_, and 𝒜_*z*_ × 𝒜_*z*_ submatrices of *W*^*EE*^, *W*^*EI*^ *W*^*IE*^ and *W*^*II*^, respectively. Henceforth we use • and º to indicate the activity state (on or off, respectively) of neurons.

The second equation can be solved^2^ to yield *z* in terms of *y*:

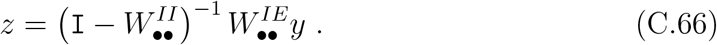

Substituting this expression into the first equation gives

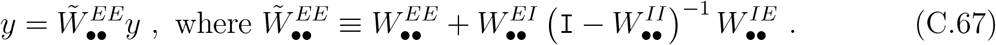

The *effective* weight matrix 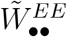 captures not only the direct recurrent connections between EPG neurons (first term) but also indirect pathways between EPG neurons mediated by the *z* neurons (second term). Henceforth, we will use to denote effective weights and matrices that include these indirect pathways. The interpretation of the second term as a sum over indirect pathways becomes explicit upon expanding the inverse:

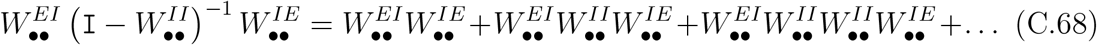

The first term represents contributions from paths that start from a compass neuron, project to an inhibitory neuron, and return to another compass neuron. The second term additionally includes an intermediate *hop* between inhibitory neurons before returning to the compass layer, and higher-order terms include progressively more such hops.

We can also infer which symmetries the effective network inherits from the symmetries of the full biological network. Motivated by the approximate symmetries of the connectome, we will assume that the full connectome and the active sets obey mirror symmetry:

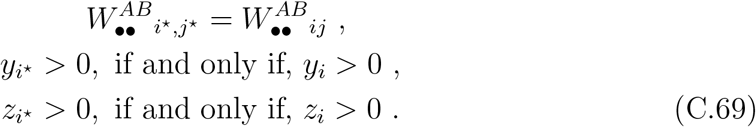

One can then see that

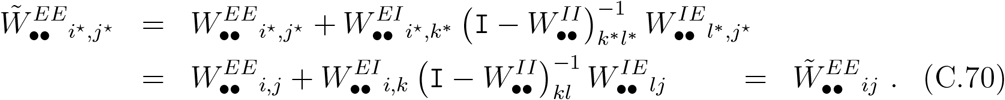

In other words, the effective weight matrix preserves mirror symmetry.

##### Effective Feedforward weights from active to inactive EPG neurons

While the requirement to realize a continuous ring attractor is most stringent on the effective recurrent couplings between the active EPG neurons (see Appendices A and B), one must also check that the assumed activity states (active or inactive) of all other neurons in the network are consistent with the input drives they receive. For the network studied here, we need to verify that inactive EPG neurons, active *z* neurons, and inactive *z* neurons receive negative, positive, and negative input drives, respectively. Such consistency checks are most easily carried out by deriving effective feedforward couplings from the active EPG neurons to the other neuronal populations. We begin by deriving the effective couplings from active to inactive EPG neurons.

The total drives, 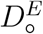, that the inactive EPG neurons receive can be written as

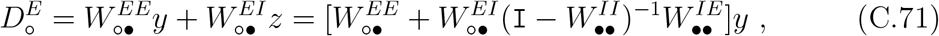

where 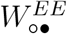 and 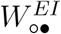 are (𝒩 − 𝒜_*y*_) × 𝒜_*y*_ and (𝒩 − 𝒜_*y*_) × 𝒜_*z*_ submatrices of *W* ^*EE*^ that connect the active EPG and *z* neurons, respectively, to the inactive EPG neurons. Therefore, the effective feedforward off-diagonal (𝒩 − 𝒜_*y*_) × 𝒜_*y*_ matrix connecting the active and inactive sectors of the EPG population is

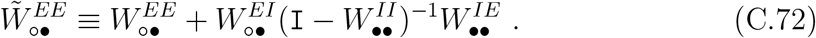

##### Effective feedforward weights from EPG to other neuronal populations

Since the derivation of effective recurrent weights between EPG neurons required us to posit the activity states of other neuronal populations, we must check that the signs of the drives to these neurons are consistent with those assumptions. Here we continue to focus on a single *z* population, but the results can be generalized straightforwardly to incorporate more than one neuronal population.

The drives, 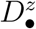, that the active *z* neurons receive are given by

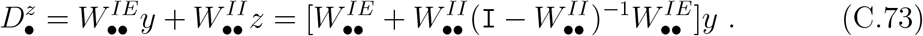

Accordingly, the 𝒜_*z*_ × 𝒜_*y*_ effective feedforward weight matrix connecting active EPG neurons to active *z* neurons is

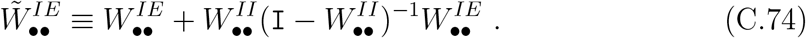

In a very similar manner, one can derive the effective weight matrix connecting active EPG neurons to inactive *z* neurons:

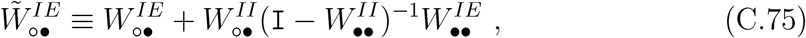

where 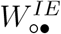 and 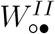 are (𝒩 − 𝒜_*z*_) × 𝒜_*y*_ and (𝒩 − 𝒜_*z*_) × 𝒜_z_ submatrices of *W* ^*IE*^ and *W* ^*II*^ that connect the active EPG and *z* neurons, respectively, to the inactive *z* neurons.

#### C.2 Ring attractor constraints on the effective weights

To summarize, in the last section we provided a prescription to obtain effective weight matrices 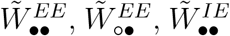, and 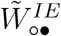, which represent, respectively, recurrent connectivity between active EPG neurons (*y*) and from the active EPG neurons to inactive EPG neurons, active *z* neurons, and inactive *z* neurons. We now describe the constraints these matrix elements must satisfy in order to realize a continuous attractor.

##### C.2.1. Constraints on effective EPG network

We observe that the 𝒜_*y*_ × 𝒜_*y*_ and (𝒩 − 𝒜_*y*_) × 𝒜_*y*_ effective weight matrices, 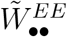 and 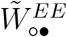 given by (B.67) and (B.72), respectively, can naturally be combined into an (𝒩 × 𝒜_*y*_ effective matrix connecting active EPG neurons with all EPG neurons. As discussed in Appendix D, any recurrent network with self-couplings is equivalent (at the level of steady-state solutions) to a recurrent network without self-couplings. According to (D.92), the equivalent weights, 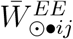, for our effective EPG network without self-couplings are

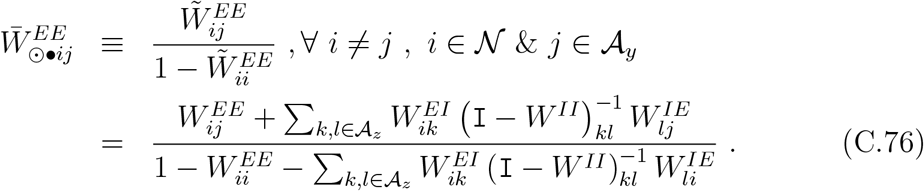

One can now read off the effective independent weight parameters (*w*’s and *u*’s) from their definitions in Appendices A and B, and impose the conditions they must satisfy to realize a continuous ring attractor.

For the symmetric model the constraints are

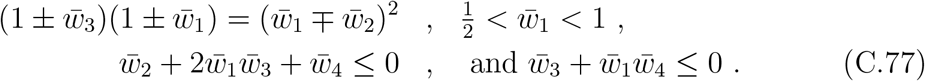

where the “bar” indicates that we are referring to the effective weights derived from (B.76). For the mirror-symmetric model the relevant equations for the effective recurrent weights read

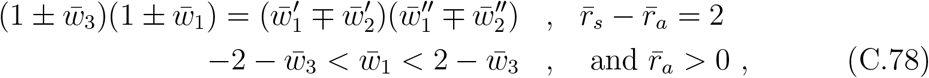

where we have defined

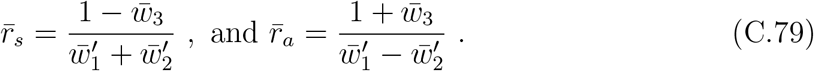

Unlike the symmetric model, in the mirror-symmetric model the effective feedforward weights from active to inactive EPG neurons are not related to the recurrent effective weights above. For every inactive neuron, the effective weights onto it must satisfy

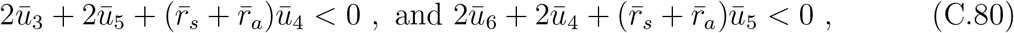

where *ū*_3_, *ū*_4_, *ū*_5_, *ū*_6_ refer to the effective weights from the active EPG neurons *y*_3_, *y*_4_, *y*_5_, *y*_6_ to the inactive EPG neuron.

##### C.2.2 Constraints on effective feedforward weights from active EPG to *z* neurons

The inequality constraints on weights from active to inactive EPG neurons were derived by requiring that the effective feedforward drive received by inactive neurons from the active EPG neurons is negative for all activity profiles (−1 ≤ *µ* ≤ 1). Since we have also derived effective feedforward weights from the active EPG neurons to the *z* neurons, the constraints implied by our assumptions about the activity state (active or inactive) of the *z* neurons can be obtained by following the same mathematical steps as for inactive EPG neurons. Explicitly, in the context of the more general mirror-symmetric model, the effective weights onto a given neuron must satisfy (B.63)

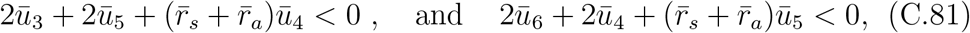

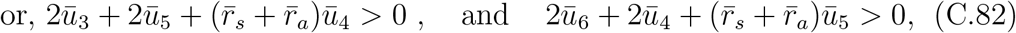

depending on whether the *z* neuron was assumed to be inactive or active, respectively. Here *ū*_3_, *ū*_4_, *ū*_1_, *ū*_6_ refer to the effective feedforward weights (C.74 and C.75) from the active EPG neurons *y*_3_, *y*_4_, *y*_5_, *y*_6_ to the *z* neuron, respectively.

For convenience, we have summarized all the constraints and their origin in Appendix C.4.

#### C.3 Scaling parameters connecting synapse counts to synapse weights

In the previous subsections we obtained the constraints that synaptic weights must satisfy in order to realize a continuous ring attractor. The connectome, however, does not directly measure synaptic weights; instead, it provides synapse counts between neurons. We assume that for each pair of pre- and post-synaptic neuron types, there exists an independent scaling parameter that converts synapse counts proportionately into synaptic weights [1]. Since there are four distinct neuronal pairs, we introduce four scaling parameters:

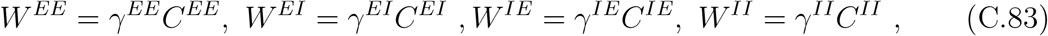

where the *C*’s are synapse-count matrices inferred from data, and the *γ*’s are proportionality constants relating synapse counts to synaptic weights.

We can now express all effective matrices as functions of these scaling factors. Explicitly, the effective weight matrices are

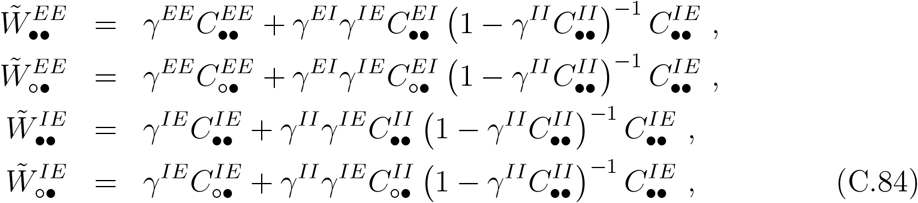

where the *C*_••_ and *C*_º•_ matrices represent synapse-count submatrices involving active neurons and active-to-inactive connections, respectively.

As noted above, the matrices involving EPG neurons can be combined into an equivalent effective weights without self-couplings^3^:

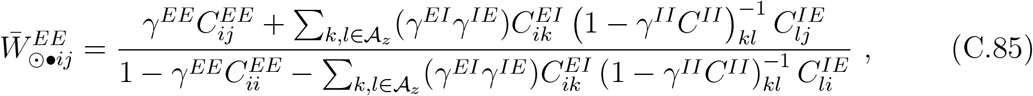

∀*i* ∈ 𝒩_*y*_ and *i* ≠ *j* ∈ 𝒜_*y*_. Here, and in what follows, we have assumed that the self-coupling terms are less than one for stability reasons (Appendix D) so that the denominators in (B.85) are positive, and the sign of the weights do not change. In a similar manner one can define an effective matrix that combines weights from the active EPG to the active and inactive *z* neurons, *i*.*e*., the matrices 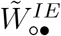 and 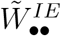 into another 𝒩 × 𝒜_*y*_ matrix, 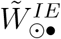:

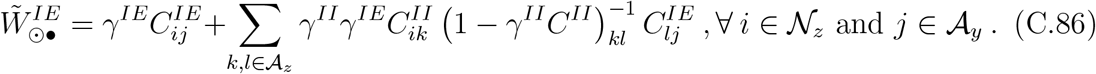

We note that since only the signs of the drives matter, rescaling the matrix elements to eliminate self-couplings is unnecessary.

To summarize, all the effective weights 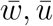 become functions of three^4^ scaling parameters:

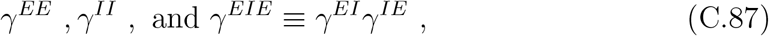

according to (C.85, C.86). As we have seen, to be a continuous ring attractor the effective synaptic weights must satisfy several equalities and inequalities. This implies that the above three scale-factors must satisfy corresponding equalities and inequalities in order to map the connectome into a viable ring attractor.

#### C.4 Finding scale-factors that lead to ring attractors

To check whether a given connectome can lead to weight matrices that satisfy all the equalities and inequalities, it is technically convenient to work in *γ*-space. Recall that starting from the synapse-count matrix, the methods described above yield all the effective weights as functions of three parameters: 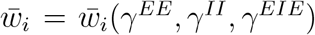. As we have discussed, in order to encode a continuous angular coordinate smoothly, consistently, and robustly, these effective synaptic weights must satisfy several equalities and inequalities, which can be recast as equalities and inequalities in *γ*-space. We now tabulate these, referring to the appropriate equations derived earlier:

Next, we will discuss constraints on effective feedforward weights from the active sector to all other neurons, which arise from requiring that the signs of the drives to these neurons are consistent with the assumed activity states. In addition to 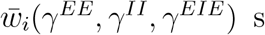, these also involve effective feedforward weights, *ū*(*γ*^*EE*^, γ^*II*^, γ^*EIE*^) s, from active EPG neurons to the other neurons, and again translate to inequalities in the *γ*-space. The inequality constraints that limit the ranges of *γ* s are tabulated as

Let us now delve into the details of how we found the three connectome-constrained ring attractor networks. We will first consider the case of symmetric effective networks. Since there are three *γ* parameters and two equations that continuous ring attractors must satisfy, a naive counting argument suggests a one-dimensional space of solutions. In Supplementary Fig. S3, we set *γ*^*II*^ = 0 and show the equality constraints as the scarlet (symmetric eigenvector condition) and violet (anti-symmetric eigenvector condition) curves in *γ*-space. The colored regions indicate excluded regions imposed by inequalities: stability of the bumps (blue), requirement of singly-peaked bump profiles (light blue), drives to active and inactive EPG neurons being positive (green) and negative (purple), respectively, and drives to the active and inactive Δ7s being positive (yellow). As shown, there are multiple discrete solutions to the equalities where the scarlet and violet curves intersect (not all shown in the plot), but only one intersection lies in the allowed white region. This point therefore corresponds to a connectome-constrained ring attractor network. While varying *γ*^*II*^ generates additional such ring attractors, as soon as 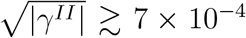, recurrent inhibition between the Δ7s renders some of them inactive, thereby violating the symmetry requirement needed to generate a symmetric effective network. In other words, we obtain a tiny one-dimensional family of viable *γ* parameters labeled by 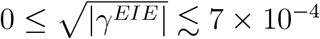, i.e., essentially a single ring attractor realization.

We next turn to mirror-symmetric networks. Since the mirror-symmetric model must satisfy three equations, and we have three *γ* parameters, we expect only discrete solutions. In Supplementary Fig. S3 we depict the *γ*-space plots for *A*_Δ_ = 6 and *A*_Δ_ = 4. For these plots we choose values of the recurrent inhibitory scale-factors such that the third equality constraint related to uniqueness and smoothness of angular representation (red curve) also passes through the intersection of the other two equality constraints. For mirror-symmetric networks, in addition to the inequality constraints relevant for the symmetric case, we must also check that the drives to the inactive Δ7 neurons are negative. For both mirror-symmetric networks we find that, once all inequalities are imposed, only unique solutions survive, as depicted in the plots.

#### C.5 Compensating mechanisms for asymmetric connectomes

While a general theory of how fully asymmetric connectomes might support continuous ring attractors is beyond the scope of this work, here we highlight a simple compensatory mechanism that allows certain asymmetric connectomes to realize a viable ring attractor. A central point is that the ring attractor constraints apply to the *effective weights*, rather than directly to synapse counts. In particular, the most stringent constraints arise from the eigenvalue degeneracy conditions, which impose equality relations among effective weight parameters. Because each effective weight is composed of multiple synapse-count parameters, the same effective connectivity can be realized by many distinct connectomes. This redundancy provides a natural route for compensation in asymmetric networks.

As a simple illustration, consider compensatory asymmetric changes between the direct EPG→EPG pathway and the indirect EPG→ Δ7 →EPG inhibitory loop,

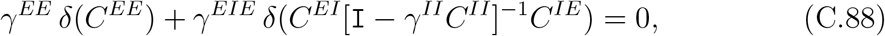

where we’ve set *γ*^*II*^ to a constant value for simplicity. To demonstrate this mechanism explicitly, we generated asymmetric matrices 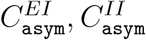 and 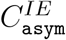 by applying random 30% perturbations^5^ to the symmetrized fly synapse-count matrices *C*^*EI*^, *C*^*II*^ and *C*^*IE*^, respectively. We then computed the compensatory 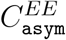 required to preserve the effective connectivity:

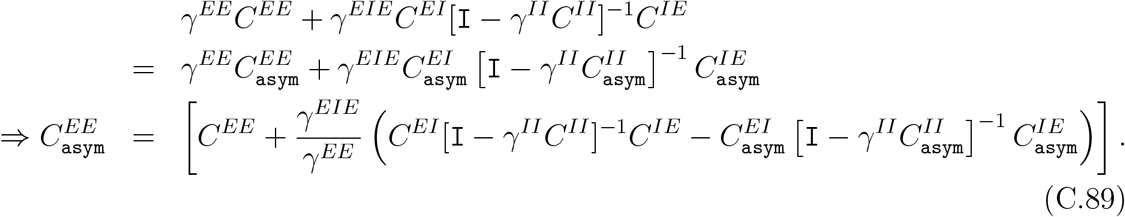

An explicit example of an asymmetric connectome that supports ring attractor dynamics is shown in Supplementary Fig. S1.

## APPENDIX D Equivalence between ring attractors with or without self-couplings

In this appendix we show that steady-state solutions in a recurrent network with self-couplings can be mapped one-to-one onto steady-state solutions of an equivalent recurrent network with no self-couplings, and vice versa. This allows us to work with the slightly simpler class of networks without self-couplings, without loss of generality.

We begin with the steady-state equation

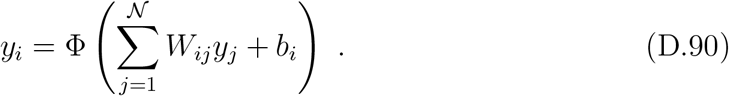

If *i* = 1, …, 𝒜_*y*_ denote the active neurons, their activities satisfy linear steady-state equations:

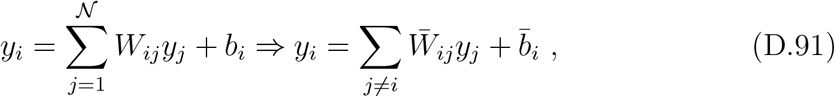

where we have defined the *equivalent* weights and drives

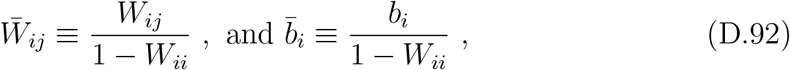

assuming *W*_*ii*_ ≠ 1 for all *i*. Thus, finding steady-state solutions in the original network is equivalent to finding steady-state solutions in a rescaled network with no self-couplings.

To obtain solutions to the nonlinear equations (A.3) one must also satisfy the inactivity inequalities

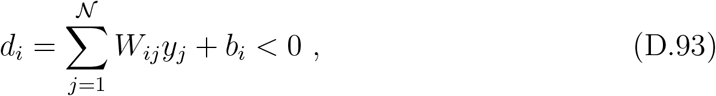

for all neurons *i* that are inactive. Typically^6^ self-couplings greater than one leads to unstable growing modes of neuronal activity, and therefore we will impose, *W*_*ii*_ < 1. Then, the above inequalities can be rewritten straightforwardly as

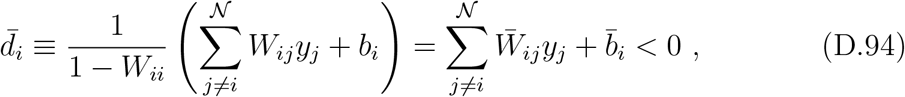

where the *i* term in the sum can be removed because *y*_*i*_ = 0. To summarize, finding a steady-state solution in an arbitrary network can be reduced to finding a solution in a network with no self-couplings, with all other weights and external drives rescaled according to (D.92).

We can now characterize the ensemble of ring attractors in networks with self-couplings using the corresponding characterization for the *equivalent* networks without self-couplings. Suppose we have already characterized weight matrices without self-couplings and their associated bump profiles, 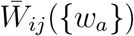 and *y*_*i*_(*µ, σ*), respectively, that produce continuous attractors. Then, by (D.92), any network with self-couplings that reproduces 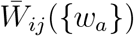 is also a valid ring attractor network. Explicitly, the ensemble of such weight matrices is

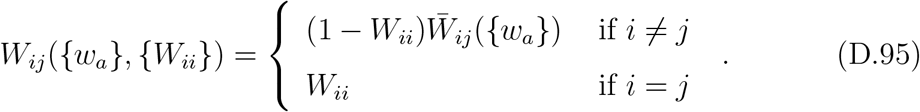

In this representation, the diagonal entries {*W*_*ii*_} provide additional free parameters that characterize the family of weight matrices, while the activity profiles remain unchanged.

## APPENDIX E Symmetries of eigenvectors for (mirror) symmetric weight matrices

In this appendix we prove a set of results about the symmetry structure of the relevant eigenvectors. These results will be used repeatedly to characterize the ensemble of continuous ring attractors and the continuum of steady states that such networks support.

### E.1 Single Eigenvector

Consider an eigenvector of *W* with eigenvalue *λ*:

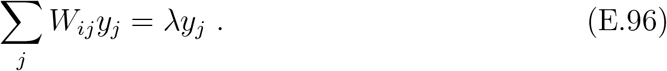

We claim that if there is only one eigenvector with eigenvalue *λ* and *W* is mirror-symmetric, then that eigenvector must be either symmetric or anti-symmetric.

Recall that a mirror-symmetric matrix satisfies

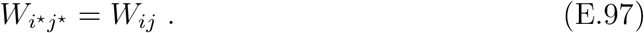

Given an eigenvector 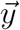, define its mirror partner 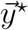 by

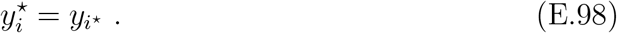

Then

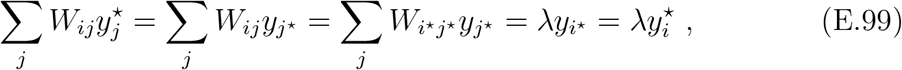

where the equalities follow, respectively, from (E.98), (E.97), (E.96), and (E.98). Thus *y*^⋆^ is also an eigenvector with eigenvalue *λ*. By assumption, the eigenspace at *λ* is one-dimensional, so 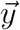 and 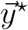 must be proportional. Since they also have the same norm, the proportionality factor must be *±*1, implying

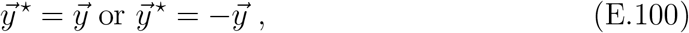

*i*.*e*., the eigenvector is symmetric or anti-symmetric, respectively. Finally, we note that a purely anti-symmetric eigenvector is ruled out in our setting because it implies a steady-state activity profile that contains negative activities.

### E.2 Symmetry properties of double degenerate eigenspaces

Next, consider a two-dimensional eigenspace associated with an eigenvalue *λ*. Let 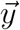 be an eigenvector that is neither symmetric nor anti-symmetric. Then 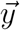 can be decomposed into a symmetric and an anti-symmetric component (see next subsection),

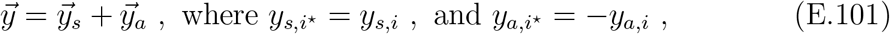

where both 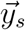 and 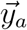 are non-zero. We now show that 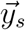 and 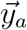 are themselves eigenvectors.

Start from the eigenvector equation and expand using the decomposition:

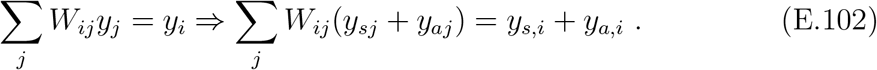

Now use the symmetry properties of 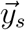 and 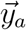 together with (E.97) to rewrite the same equation in mirrored form:

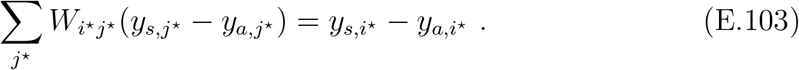

Thus both (*y*_*s,i*_ + *y*_*a,i*_) and (*y*_*s,i*_ − *y*_*a,i*_) are eigenvectors with the same eigenvalue, and therefore so are 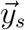 and 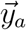. In other words, any doubly-degenerate eigenspace can be decomposed into a symmetric and an anti-symmetric eigenspace.

If the entire two-dimensional eigenspace were symmetric, then the average encoded position would always be at zero and the network could not support continuous angular encoding; we therefore do not consider this case. A completely anti-symmetric eigenspace is disallowed because negative activities are not permitted for the steady states. Hence, the only viable case for continuous angular encoding is a mixed eigenspace with one symmetric and one anti-symmetric eigenvector.

### E.3 Decomposition in terms of symmetric and antisymmetric vectors

In this section we show that both the dynamics and the steady-state equations naturally decompose into separate equations for the symmetric and anti-symmetric components of the activity. First, note that any activity vector can be decomposed as

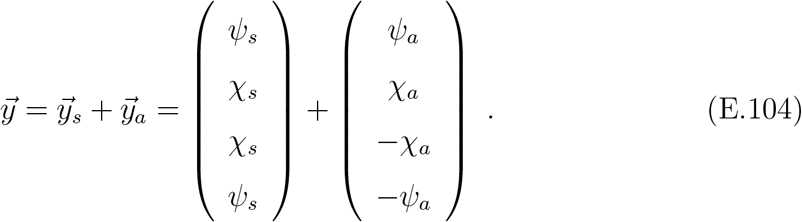

The full space is spanned by two symmetric basis vectors 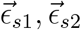 and two anti-symmetric basis vectors 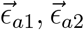. Moreover, the symmetric and anti-symmetric sub-spaces are orthogonal. The symmetric and anti-symmetric projections of any vector can be obtained via

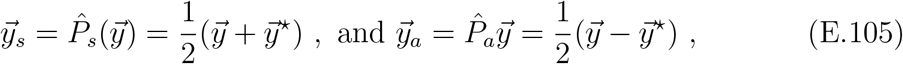

where 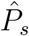 and 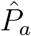 are the symmetric and anti-symmetric projection operators, respectively.

Now consider the time-evolution equation for neurons in the active set

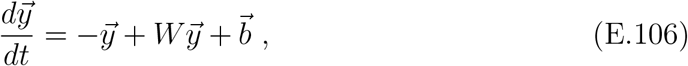

Where 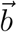 is a constant external drive and from here onwards we will set the neuronal time constant to be one. In the driven ring attractor network (Appendix F) we will take 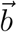 to be symmetric, while for velocity integration (Appendix I) an anti-symmetric 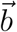 becomes relevant. Applying the projection operators yields two equations,

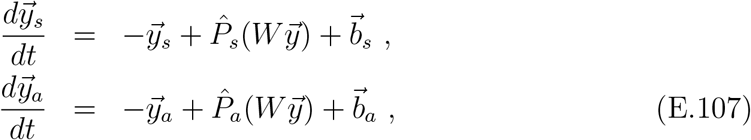

one for the symmetric and one for the anti-symmetric component. We now show that for a mirror-symmetric matrix the dynamics of these components decouples.

For the symmetric part,

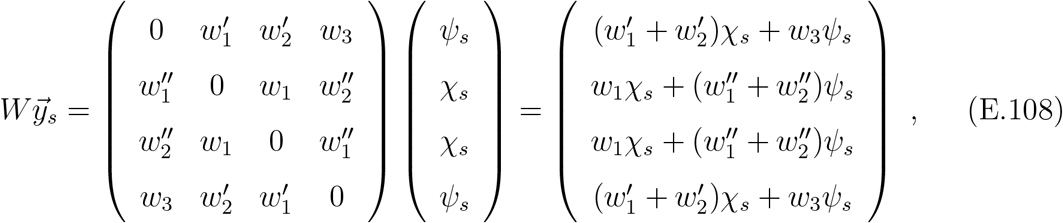

which is again symmetric. It is therefore convenient to work with the associated two-dimensional vector

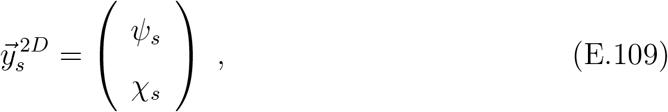

whose dynamics are given by

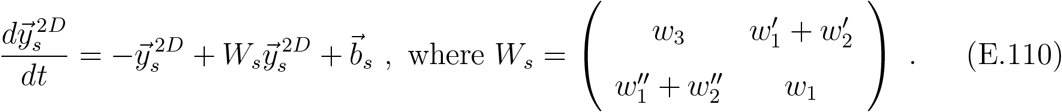

Similarly, for the anti-symmetric part,

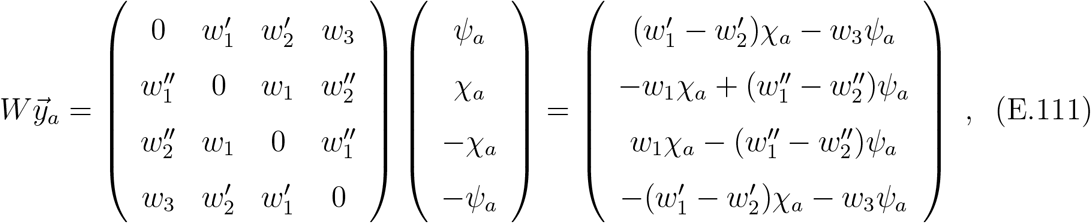

which is anti-symmetric. Again it is convenient to work with the two-dimensional vector

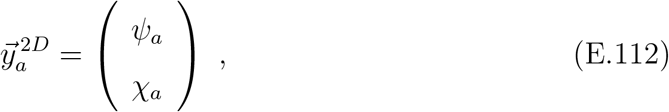

with dynamics equation

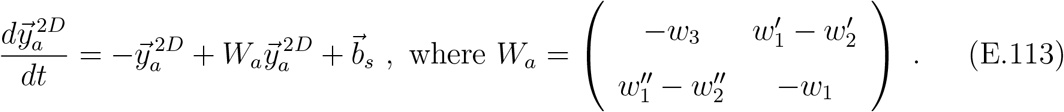

## APPENDIX F Driven continuous attractors

In this appendix we consider ring attractors that can encode angles continuously but require an external drive to sustain a localized bump of activity. Specific examples of such attractors in the context of threshold-linear networks have been identified recently [2]. Here we provide general conditions for obtaining threshold-linear symmetric networks of this type. One advantage of driven models over self-sustaining networks is that only a single degeneracy condition is required. However, as we will see, the resulting angular representation is less robust to external inputs and noise.

### F.1 Considerations for obtaining a continuum of steady-state profiles

Since an external drive is present, we consider dynamical equations of the form

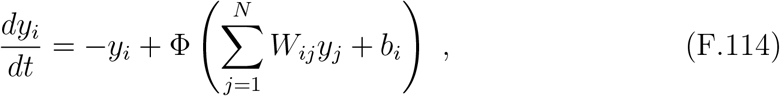

where *b*_*i*_ is a constant drive and from here onwards we will set the neuronal time constant to be one. As with the self-sustaining networks, we focus on steady-state solutions, which must satisfy

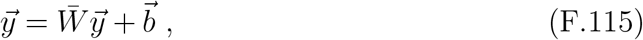

where 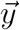 and 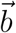 are *A*_*y*_-dimensional vectors corresponding to the active neuronal directions, as in Appendix E. We assume that 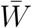 has a single eigenvector with eigenvalue one. Since the active neurons comprise a linear dynamical system, the general solution to (F.115) can be written as a sum of a homogeneous and a particular solution:

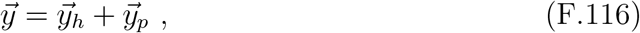

where the homogeneous component satisfies the eigenvalue equation

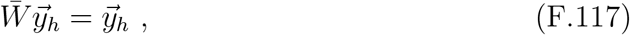

and, as we show below, is responsible for generating the required continuum of steady-state profiles, and 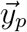 is any particular solution of (F.115).

We first show that the particular solution can be taken symmetric. Consider any 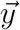 satisfying

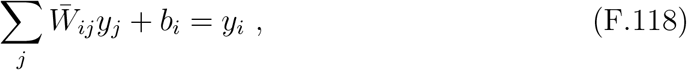

and assume that the external drive does not depend on neuron identity, so that *b*_*i*_ = *b* for all *i*. Then its mirror 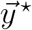 also satisfies the same equation:

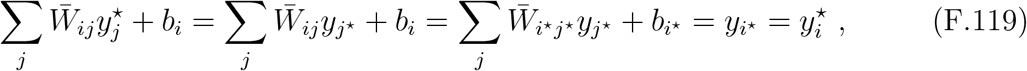

where in the first equality we used the definition of a mirror, 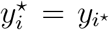, and in the second equality we used the symmetry property 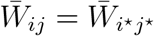 together with 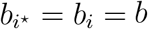. Therefore one can form a symmetric configuration

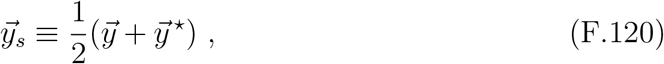

which also satisfies (F.115) and can serve as a particular solution. We note that 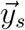 depends linearly on *b*, which will be important later.

We next consider homogeneous solutions. As shown in Appendix E, when there is only a single eigenvalue equal to one, the corresponding eigenvector must be either symmetric or anti-symmetric. If both the homogeneous and particular solutions are symmetric, then a continuous attractor with varying average location is not possible: the average location is pinned to the center. Thus we require an anti-symmetric eigenvector 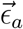, *η* being a free parameter, so that the homogeneous solutions take the form 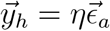, and the general steady-state solution can be written as

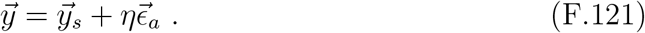

### F.2 Steady-state profiles for the symmetric driven ring attractor networks

We now focus on the case of 𝒩 = 8 neuronal/computational units with steady-state configurations involving 𝒜 = 4 active neurons, as is pertinent for the fly ring attractor network. For simplicity, we specialize to the symmetric case, although most of the results should generalize to mirror-symmetric networks as well. In Appendix A we derived the condition for having an anti-symmetric eigenvector in the symmetric model. To summarize, the weights must satisfy

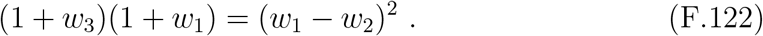

Stability is ensured as long as

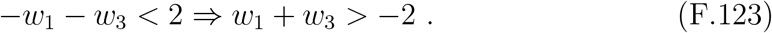

The anti-symmetric states are given by

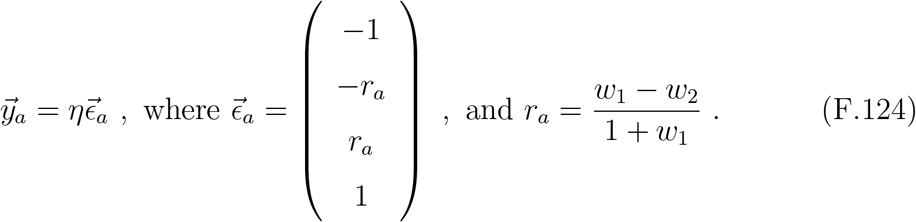

We next construct a symmetric particular solution. Substituting the symmetric ansatz

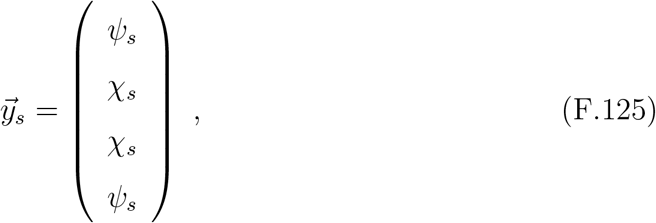

into (F.115), we obtain the matrix equation

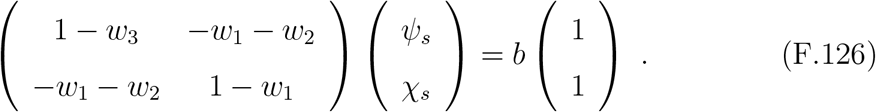

Since we have assumed 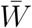 is singly degenerate, the above matrix is invertible, and therefore

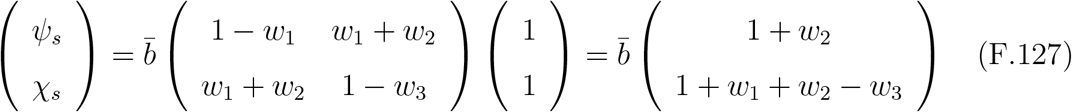

where we have defined

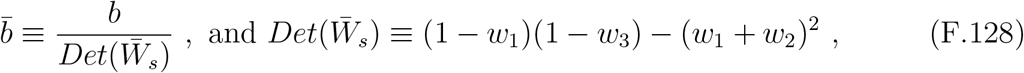

is the determinant of 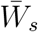, which must be nonzero because 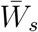 is nonsingular.

Thus the full bump profile can be written as

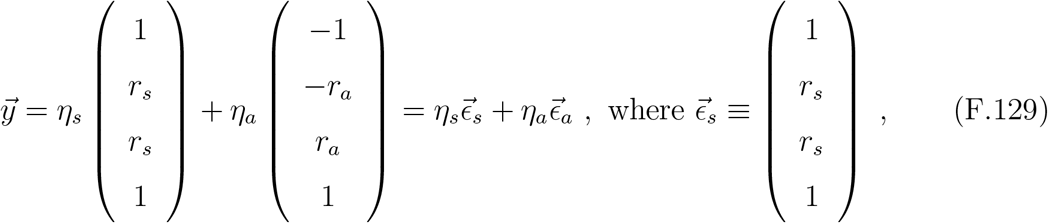

and

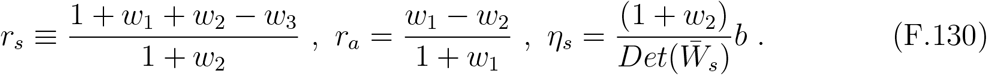

As in the self-sustaining network, we require all the activities to be positive. Requiring *y*_3_ *>* 0 for *η*_*a*_ = 0 implies *η*_*s*_ must be positive and therefore *r*_*s*_ must also be positive to ensure *y*_4_ *>* 0. Further to ensure that *y*_3_ and *y*_6_ remain non-negative, the range of *η*_*a*_ gets constrained to lie within, *η*_*s*_ ≤ *η*_*a*_ ≤ *η*_*s*_. Next let us consider the extreme geometry when *η*_*a*_ = *η*_*s*_, *y*_3_ = 0. To ensure that we have a single peak in the middle, *y*_5_ *> y*_4_, and therefore *r*_*a*_ must be positive. Thus to summarize, the following inequalities must be satisfied.

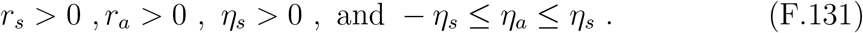

### F.3 Robustness of angular representation

(F.129) and (A.22), the steady-state activity profiles for the self-sustaining and driven cases look very similar: in both cases they can be expressed as a linear combination of a symmetric and an anti-symmetric vector,

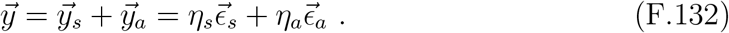

There is, however, a key difference between the two models. In the self-sustaining model the space of steady states is two-dimensional, and both *η*_*s*_ and *η*_*a*_ can vary continuously. In the driven case, *η*_*s*_ is fixed by the external drive *b*, and only *η*_*a*_ can vary. Equivalently, while 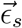 is an eigenvector of 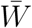 with eigenvalue one in the self-sustaining model, it is not an eigenvector in the driven case. As we now explain, this can lead to systematic perturbative instabilities in the angular representation.

#### F.3.1 Driven case

Consider the linearized dynamics under an additional perturbative drive 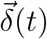:

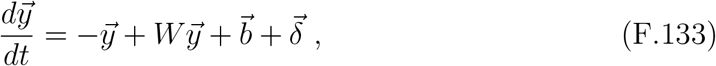

where we have set the neuronal time constant *τ* = 1. In general, the perturbation can be expanded in the eigenbasis:

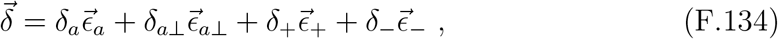

where 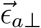 denotes an antisymmetric eigenvector orthogonal^7^ to 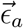, and 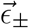 denote the pair of symmetric eigenvectors. If the *δ*’s are small fluctuations around zero, one expects the angular representation to fluctuate around its mean value. Since the angular variable corresponds to an integrating mode, one expects diffusive growth of its variance [3], a generic feature of continuous attractor dynamics, in both the driven and self-sustaining networks. The question we address here is what happens under a systematic perturbation.

While a comprehensive analysis is beyond the scope of this paper, we consider the case where the perturbation is aligned with the current activity direction, as might occur for a visual stimulus (landmark) aligned with the head direction. Suppose that at *t* = 0,

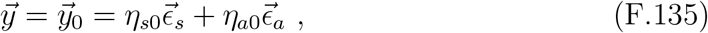

and that a systematic perturbation 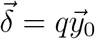 persists until time *t* = *t*_*e*_. One would expect the encoded angle to remain stable under such a perturbation. We will show that while this is true for the self-sustaining model, the driven model exhibits systematic angular drift.

To see this, consider a generic perturbation and decompose the activity as

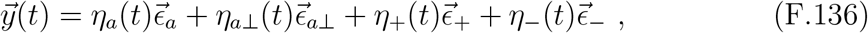

which is possible as long as the active set does not change and linearity holds. Substituting into (F.133) gives

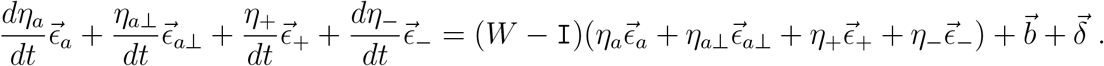

Noting that 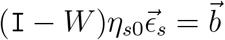, and defining *ξ*’s via

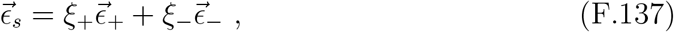

we obtain, using the eigenvalue equations,

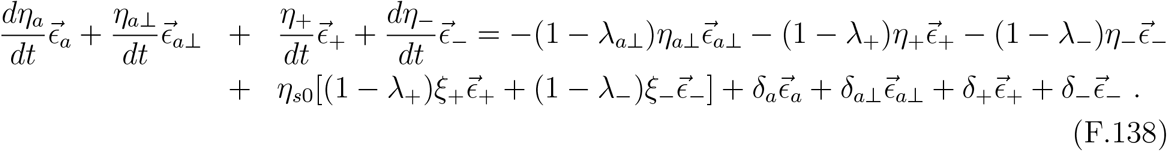

Equating components yields

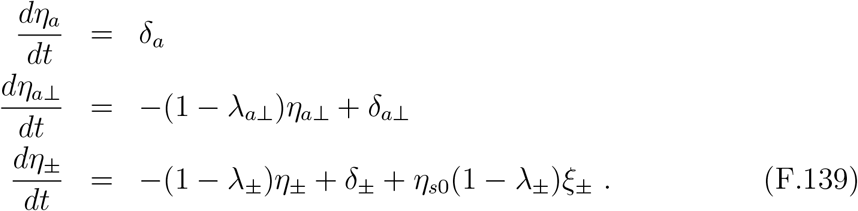

For a perturbation aligned with 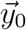, we have

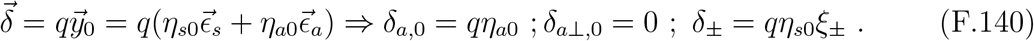

We note that 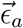 is an integrating mode, while the remaining modes decay on neuronal time scales. Accordingly, the 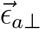 mode is not excited, and the angular coordinate evolves as

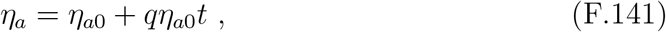

whereas the symmetric components quickly saturate (*dη*_*±*_/*dt* → 0):

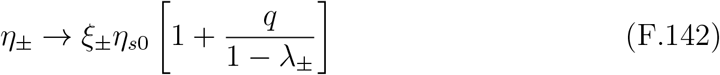

The angular representation is given by

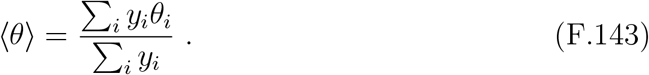

By symmetry, the numerator depends only on the antisymmetric part 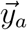, whereas the denominator depends only on the symmetric part 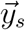. It then follows from (F.141) and (F.142) that the numerator varies linearly in time while the denominator rapidly approaches a constant, leading to a time-dependent angular representation. Moreover, the angle does not return to its original value once the perturbation is removed. When *q* → 0, *η*_*a*_ remains at its value at *t* = *t*_*e*_:

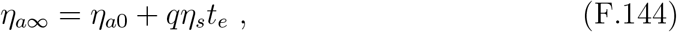

while the *η*_*±*_ modes return to their original values due to decay:

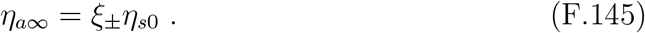

In other words,

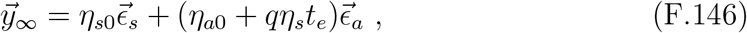

which clearly represents an angle that is different from the initial value. Supplementary Fig. S2 demonstrates the angular drift during the transient perturbation aligned with the initial head direction.

#### F.3.2 Self-sustaining network

To contrast this behavior, we consider the self-sustaining model. In this case both 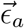 and 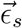 are eigenvectors with eigenvalue one. Decomposing the network dynamics in terms of 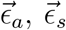, and their orthogonal symmetric and antisymmetric counterparts 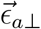 and 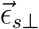, one obtains component equations analogous to the driven case:

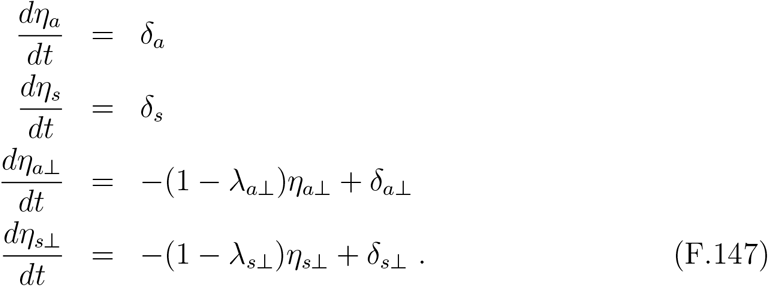

A perturbation 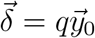 yields

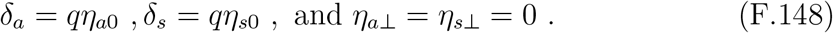

Thus, the decaying modes are never excited and the zero modes integrate:

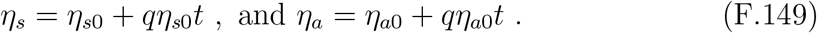

It is now clear that the angular representation remains stable throughout:

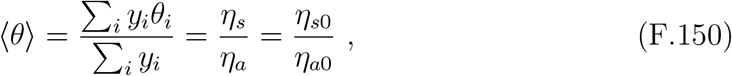

as simulations demonstrate in Supplementary Fig. S2.

Although in general perturbations will not be exactly aligned with the current head direction, the analysis highlights a broader point. In the driven network, systematic perturbations along any direction in the two-dimensional space of steady states spanned by 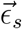 and 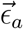 lead to an error in the represented angle. In contrast, in the self-sustaining network errors do not occur for perturbations along the 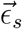 direction. Moreover, for both networks one naturally expects errors under systematic perturbations along the 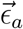 direction, since that corresponds to a velocity signal. Thus, self-sustaining networks provide a more robust angular representation, whereas driven networks are susceptible to systematic angular drift.

## APPENDIX G Connectome constrained models with a single inhibitory neuron

### G.1 Effective EPG network

Throughout most of the paper, we considered a two-population model in which each population had 𝒩 = 8 neurons and the synapse counts obeyed 8-fold symmetry. Another biologically relevant architecture that preserves 8-fold symmetry is one in which a single^8^ inhibitory neuron is coupled uniformly to all excitatory neurons.

Let *z* denote the activity of the inhibitory neuron that provides indirect inhibitory pathways between the EPG neurons. The network dynamics are modeled as

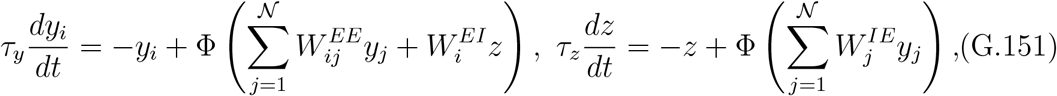

where, as before, *τ*_*z*_ and *τ*_*y*_, are neuronal time constants, and *W* ^*EE*^ is the recurrent weight matrix for the EPG neurons. In contrast, *W* ^*EI*^ and *W* ^*IE*^ are 𝒩 -dimensional vectors that reciprocally couple the excitatory neurons to *z*. We have assumed that there is no self-coupling term for the *z* neuron.

Focusing on a steady-state configuration for the active excitatory neurons, we obtain

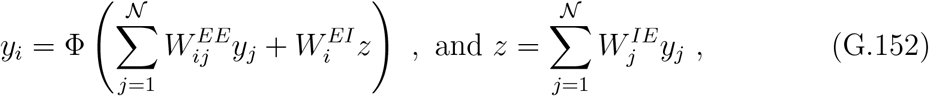

where we assumed that the inhibitory neuron is active in order to allow inhibitory interactions to contribute to the steady-state activity profile. Substituting the *z* neuron’s activity into the first equation yields

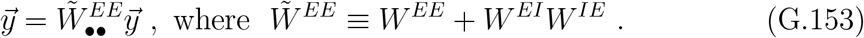

As before, 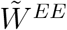 is the *effective* weight matrix involving the active excitatory neurons. Explicitly, its matrix elements are

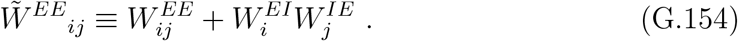

### G.2 Correspondence of the two different architectures

Since there are no recurrent inhibitory synapses, we can connect weight matrices to synapse-count matrices using only three scaling parameters:

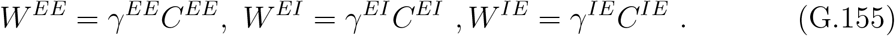

The effective matrix can then be written as

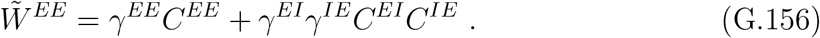

At this point it is useful to impose 8-fold circular symmetry on the synapse-count matrices. This implies that the *E* ↔ *I* connections do not depend on the identity of the excitatory neuron:

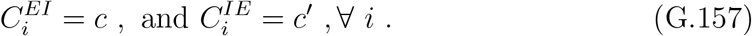

Therefore, the elements of the effective weight matrix are

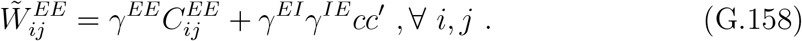

From inspection of the effective-weight expression in the architecture with 𝒩= 8 inhibitory neurons, we see that if we substitute

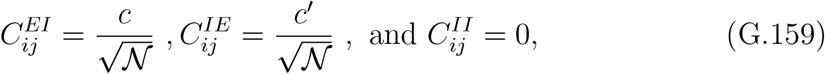

then the expressions for effective weights, 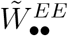 and 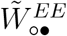, derived for 𝒩 inhibitory *z* neurons in (B.84) are equivalent to (G.158). Thus, the present architecture is a special case of the more general inhibitory-ring architecture analyzed previously.

Continuing on, one can compute the effective weights of a network without self-couplings using (D.92):

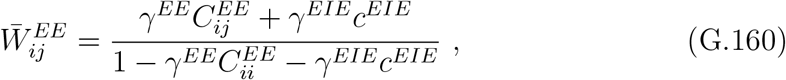

where we have defined

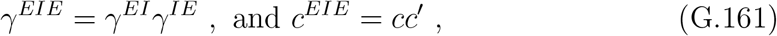

for convenience. An important difference between the two architectures is that while in the symmetric inhibitory-ring architecture the effective weight matrix depended on three scale-factors, here it depends only on two: 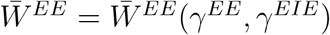. Since these effective weights must satisfy two equations to realize a continuous attractor, we therefore expect only a finite number of such instantiations (if any) to exist. In contrast, the symmetric inhibitory-ring architecture admits a continuum of such instantiations when solutions exist.

In addition to the equality constraints (A.10, A.13), the effective weights must also satisfy the inequality constraints (A.11, A.14, A.35, A.26) to ensure that we indeed have 𝒜_*y*_ = 4. We also must enforce consistent synapse signs, *γ*^*EE*^ > 0 and *γ*^*EIE*^ < 0. Finally, note that the inequalities involving Δ7 (in)activity are trivial here: since *z* receives only excitatory input from the *y*’s, it will be active as long as *γ*^*IE*^ > 0.

### G.3 Connectome-constrained ring attractor with ER6 ring neurons

The biological motivation for analyzing this model case is that a group of ring neurons (ER6 neurons) has been proposed as an alternate source of inhibition to Δ7 neurons [4]. The EPG and ER6 neurons form an excitatory–inhibitory loop, analogous to the EPG–Δ7 loop, as can be seen from the connectome in Supplementary Fig. S5a. The bidirectional synapse counts are approximately uniformly distributed along the EPG computational units, and thus it is appropriate to treat the ER6 neurons as a single population (*z*) providing inhibition to the EPG neurons, as depicted in Supplementary Fig. S5b.

As discussed in the previous subsection, the two scale parameters *γ*^*EE*^ and *γ*^*EIE*^ parametrize the realizable effective networks and their effective weights. This defines a two-dimensional surface in the three-dimensional weight space (Supplementary Fig. S5c). The ensemble of ring attractor networks intersects this surface at a unique location, demonstrating the viability of a unique connectome-constrained ring attractor. Supplementary Fig. S5d shows the existence of this continuous attractor network in *γ*-space, analogous to Supplementary Fig. S3d-f for the EPG–Δ7 network. The corresponding effective weight parameters are *w*_1_ = 0.797, *w*_2_ = −0.271, *w*_3_ = −0.365, which imply *r*_*a*_ = 0.559, quite close to the *r*_*a*_ values obtained from the EPG–Δ7 network. Accordingly, the EPG activity profiles generated in this network are quite similar to those in the connectome-constrained network involving EPG and Δ7 neurons (Supplementary Fig. S5e).

## APPENDIX H Dimensionality arguments

In this appendix we present simple counting arguments that can be used to estimate the dimensionality of both the effective ring-attractor spaces and the connectome-constrained ring-attractor spaces for multi-population models that are more general than those considered in the main text. We first explain the logic behind these counting arguments using the networks considered for the fly compass system. We then show how these arguments can be generalized to larger effective ring-attractor networks by considering networks in which both the total number of neurons (𝒩) and the number of active neurons (𝒜) are arbitrary even numbers^9^. Finally, we explain how details of the network architecture must be incorporated in order to derive the dimensionality of connectome-constrained ring-attractor networks, using an example architecture of interest.

### H.1 Networks relevant for fly head direction system

We begin by considering the symmetric model in which all neurons other than the compass neurons are required to remain active. Since in the fly there are 𝒜 = 4 active compass neurons, there are (𝒜 − 1 =) three independent recurrent weight parameters^10^ within the active subnetwork that must satisfy two eigenvalue constraints. Thus we expect a one-dimensional space of ring-attractor networks, which was parameterized by *r*_*a*_ in Appendix A. In addition, there is one effective weight parameter connecting the active compass neurons to the inactive compass neurons. Therefore, the total dimensionality of the symmetric effective ring-attractor network is𝒜 − 2 = 2. Note that the inequalities that the ring-attractor networks must also satisfy make this one-dimensional space finite.

Next, let us consider mirror-symmetric networks. An arbitrary network of 𝒜 neurons without self-couplings has 𝒜 (𝒜− 1) independent weights, but mirror symmetry reduces this number by half. For the fly (𝒜 = 4), we therefore have six independent weight parameters characterizing the recurrent active subnetwork of compass neurons. These parameters must satisfy two eigenvalue constraints as well as one smoothness condition, resulting in a three-dimensional submanifold of ring-attractor networks. This submanifold was parameterized by *r*_*a*_, *w*_1_, *w*_3_ in Appendix B. It is further bounded by inequalities along certain directions.

One can also compute the dimensionality of the ring-attractor networks within the space of fully mirror-symmetric networks involving all neurons. Due to mirror symmetry, such a network has 𝒩 (𝒩 − 1)/2 independent weights. For the 𝒩= 8 fly network, this number equals 28, which must still satisfy three equations, leading to a 25-dimensional solution space.

It is also useful to determine the dimensionality of the parameter space that characterizes the relevant effective network (Appendix B), involving only the recurrent weights within the active neurons and the feedforward weights from the active to the inactive neurons. Since the outgoing effective weights from each active neuron to all other (𝒩 − 1) neurons are independent of each other, while the outgoing synapses from its mirror pair are characterized by the same parameters, we obtain a total of 𝒜 (𝒩 −1)/2 independent parameters. For the fly network this number equals 14, which again must satisfy the three equations. This leaves an eleven-dimensional manifold of continuous ring attractors within the effective network parameter space.

Finally, let us determine the dimensionality of ring-attractor networks that can be realized given a fly-like connectome that admits at least one such solution. As discussed earlier, it is natural to associate each type of synaptic pair with an independent scale factor that converts synapse counts into synaptic weights. Since the EPG-Δ7 connectome has 𝒫 = 2 types of neuronal populations, we have 𝒫^2^ = 4 independent scale factors. However, the incoming and outgoing synapses between the EPG compass neurons and the Δ7 neurons always appear together in the expressions for the effective weights involving the compass neurons. As a result, the dimensionality of connectome-constrained effective networks is actually three. This implies that symmetric networks, which must satisfy two equality conditions, are expected to form a one-dimensional space of connectome-constrained ring-attractor realizations. In contrast, mirror-symmetric networks must satisfy three equality conditions, and therefore we expect only discrete (zero-dimensional) realizations.

### H.2 Effective networks

We now explain how the above arguments can be generalized to networks of arbitrary size. To keep the discussion technically simple, we specialize to the case in which both 𝒩 and 𝒜 are even.

As discussed for the fly network, the key requirement for constructing a continuous ring attractor is that the effective subnetwork of active compass neurons must satisfy two eigenvalue conditions and, in the mirror-symmetric case, an additional smoothness condition. For the symmetric network we have min(𝒜 − 1, /2) independent weight parameters^11^, while the mirror-symmetric network has 𝒜 (𝒜 − 1)/2 independent parameters. Following the arguments presented in the previous subsection, the number of free parameters that characterize the manifold of continuous ring attractors is therefore

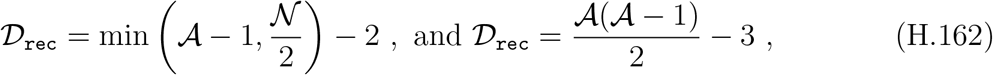

for symmetric and mirror-symmetric networks, respectively. Because the equalities and inequalities that the effective weights must satisfy are highly nonlinear, the resulting solution space may consist of disconnected regions in weight space that realize continuous ring attractors. Our dimensionality arguments apply to each such region.

We can also compute the dimensionality of ring attractors viewed as a submanifold of all networks that respect a given set of symmetries. A network obeying both circular symmetry and reciprocal symmetry has 𝒩/2 independent weights, whereas a mirror-symmetric network has a much larger number of parameters, 𝒩 (𝒩 − 1)/2. The arguments described for the fly network generalize directly and yield the dimensionalities of the symmetric and mirror-symmetric ring-attractor networks as

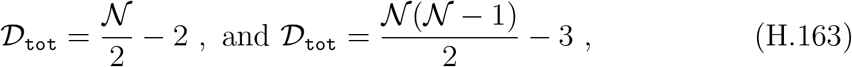

respectively.

One can also compute the number of effective weight parameters in these two types of networks. Since there are 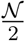 independent effective weight parameters in the symmetric network and 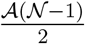 in the mirror-symmetric networks, the number of effective parameters characterizing the ring attractors is

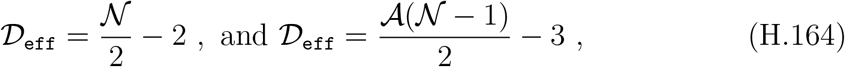

respectively.

### H.3 Connectome-constrained manifold

Our goal in this subsection is to demonstrate, using a specific example, how one might generalize the derivation of the dimensionality of connectome-constrained ring-attractor networks when more than two neuronal populations are present. The derivation can be broken into two steps. In the first step one computes the dimensionality, 𝒟_real_, of realizable effective networks obtained by scaling synapse counts in a given connectome to produce synaptic weights. As we will see below, this step depends strongly on the network architecture, and we therefore focus on a particular illustrative case. The second step involves computing the dimensionality, 𝒟_cc_, of connectome-constrained ring-attractor networks from 𝒟_real_. This step can be carried out more generally.

We begin by estimating 𝒟_real_. Consider a network composed of 𝒫 distinct neuronal populations, including the compass neurons. There are potentially 𝒫^2^ distinct pre–post synaptic pairs, and one might naively expect that such a connectome could realize a 𝒫^2^-dimensional space of effective networks. However, this need not be the case for several reasons, including: (i) not all populations may interact with one another, (ii) some scale factors may only appear in fixed combinations when the network is reduced to the effective compass-neuron network by summing over indirect paths, and (iii) recurrent couplings within a population may be absent. Below we consider an example architecture to illustrate how these complications can be handled when computing 𝒟_real_.

Consider the case in which the (𝒫 − 1) non-compass populations each connect internally and with the compass neurons, but do not interact with other non-compass populations. In this situation there are only 3(𝒫 − 1) + 1 = 3𝒫 − 2, distinct synaptic pairs. The factor of three accounts for incoming connections to the compass neurons, outgoing connections from the compass neurons, and recurrent couplings within each non-compass population. The additional one corresponds to recurrent couplings between the compass neurons. Furthermore, the incoming and outgoing synapses between the compass neurons and any other population always appear together. Therefore, the number of independent scale-factor combinations determining the dimensionality of the realizable effective networks is

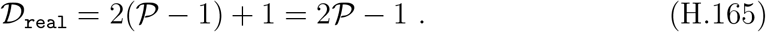

For the EPG–Δ7 connectome, 𝒫 = 2, leading to a three-dimensional space of realizable networks (Appendix C).

In addition, if any non-compass neuronal population lacks recurrent couplings within it, the dimensionality should be reduced further. For instance, because the self-coupling of the ER6 neuron can be ignored, the EPG–ER6 connectome-based space of realizable networks is two-dimensional. In summary, the number of independent scale-factor combinations, 𝒟_real_ ≤ 𝒫^2^, that relate the connectome to the effective network depends on both the number of neuronal populations and the specific network architecture.

Once 𝒟_real_ is determined, the dimensionality of the connectome-constrained ring-attractor manifold can be estimated more directly. For a fixed connectome, all effective weights depend solely on the scale factors. Because these effective weights must satisfy either two or three constraints—depending on whether the network is symmetric or mirror-symmetric—the resulting dimensionality is

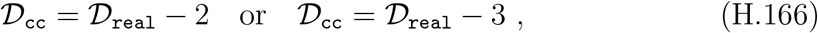

respectively, provided that at least one solution exists. As with the solution space of the effective networks, there may be multiple disconnected regions of connectome-constrained ring networks, and our estimates apply to each such region.

For the EPG–Δ7 connectome we have 𝒟_real_ = 3 relevant synaptic combinations. Consequently, the fly connectome admits a one-dimensional family of symmetric ring-attractor networks and a unique (zero-dimensional) set of mirror-symmetric ring attractors. For the EPG–ER6 network, the absence of an ER6–ER6 self-coupling implies 𝒟_real_ = 2, yielding only a unique (zero-dimensional) connectome-constrained ring-attractor realization.

## APPENDIX I Velocity Integration

In this appendix we provide an explicit example of a velocity drive that continuously deforms the shape of the activity profile, allowing the activity bump to track the angular location smoothly and faithfully. We focus exclusively on activity changes for a fixed set of active neurons within the effective network, and therefore consider the linear dynamics

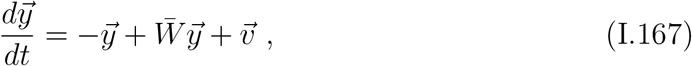

where 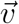 is a constant velocity drive, and we have again set the neuronal time constant to be one. As in Appendix F, the solution can be written as the sum of a *particular* solution, 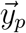, and a solution, 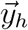, to the homogeneous equation,

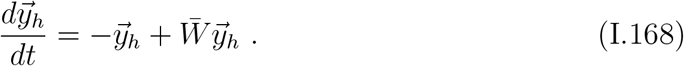

The homogeneous equation is linear and its solution can be decomposed into eigen-modes:

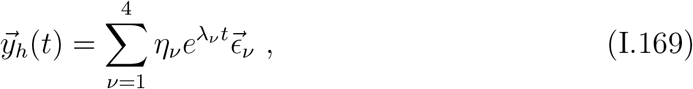

where the 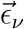 are eigenvectors satisfying

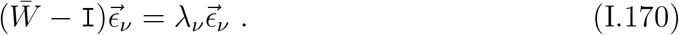

For the class of models considered here, two eigenvalues of 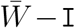 are zero, corresponding to the steady-state profiles, while the remaining two eigenvalues are negative by stability considerations. Consequently, the general solution to the homogeneous equation can be written as

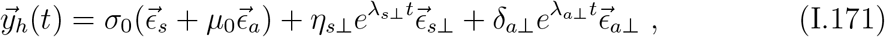

where *λ*_*s*⊥_, λ _*a*⊥_ < 0, and the four integration constants are *σ*_0_, *µ*_0_, *η*_*s*⊥_, and *η*_*a*⊥_. Since the modes associated with *λ*_*s*⊥_ and *λ*_*a*⊥_ decay exponentially, they become negligible at long times and will be ignored in what follows.

We now turn to the construction of a particular solution. For illustrative purposes, we consider a velocity drive directed along the anti-symmetric direction,

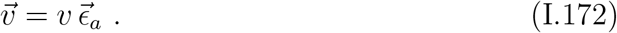

We claim that a particular solution can be chosen to be of the form

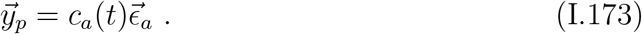

Using the fact that 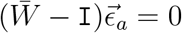, substitution into (I.167) yields

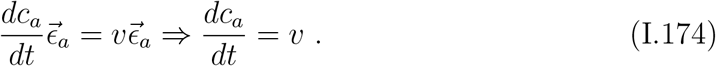

A particular solution is therefore

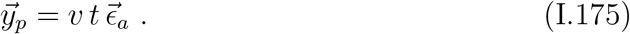

Combining the homogeneous and particular solutions, the full solution can be written as

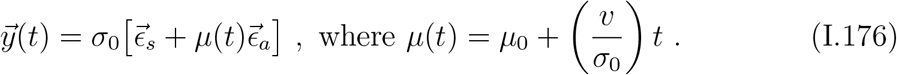

In other words, the rate of change of the encoded angular location, *ω* ≡ *d* ⟨*θ*⟩ /*dt*, is proportional to the velocity drive *v*:

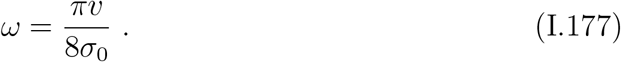

Interestingly, fruit flies navigating in darkness are observed to track head direction only proportionally. This suggests that additional sensory cues may be required to calibrate the proportionality factor and achieve precise angular tracking.

## Footnotes

1 By looking at the µ = 0 case, for instance.

2 Assuming (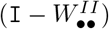) is nonsingular.

3 Mostly for conceptual simplicity we have also rescaled the weights onto the inactive neurons although self-loops involving them do not exist. Such rescaling does not affect any calculations as only the sign of the input drives to the inactive neurons matter but this uniform rescaling of all the EPG neurons, irrespective of whether they are active or inactive, allows us to conceptualize an effective EPG network.

4 We point out that although in (B.86) γ^*IE*^ > 0 appears individually, it can be factored out and therefore its value does not affect the inequality constraints.

5 We required all synapse counts to remain positive; typically this constraint was satisfied after ∼ 𝒪 (10) attempts with simple random sampling.

6 It may be possible for inhibitory recurrent connections to ameliorate the instability, but we will not consider this possibility here.

7 W_*a*_ is a symmetric matrix and therefore has orthogonal eigenvectors.

8 This setup trivially generalizes to accommodate more than one such inhibitory neuron, as long as each neuron is coupled uniformly to all excitatory units.

9 Similar arguments for cases in which the number of neurons (total or active) is not even lead to results with only minor differences.

10 As usual we ignore the self-couplings.

11 We remind the reader that because of circular and reciprocal symmetries there are only 𝒩/2 independent weight parameters, not counting the self-coupling.

## References

[1] Varshney, L. R., Chen, B. L., Paniagua, E., Hall, D. H. & Chklovskii, D. B. Structural properties of the caenorhab-ditis elegans neuronal network. PLoS computational biology 7, e1001066 (2011).

[2] Hildebrand, D. G. C. et al. Whole-brain serial-section electron microscopy in larval zebrafish. Nature 545, 345–349 (2017).

[3] Svara, F. et al. Automated synapse-level reconstruction of neural circuits in the larval zebrafish brain. Nature Methods 19, 1357–1366 (2022).

[4] Eichler, K. et al. The complete connectome of a learning and memory centre in an insect brain. Nature 548, 175–182 (2017).

[5] Bock, D. D. et al. Network anatomy and in vivo physiology of visual cortical neurons. Nature 471, 177–182 (2011).

[6] Scheffer, L. K. et al. A connectome and analysis of the adult drosophila central brain. Elife 9, e57443 (2020).

[7] Dorkenwald, S. et al. Neuronal wiring diagram of an adult brain. Nature 634, 124–138 (2024).

[8] Schrödel, T., Prevedel, R., Aumayr, K., Zimmer, M. & Vaziri, A. Brain-wide 3d imaging of neuronal activity in caenorhabditis elegans with sculpted light. Nature methods 10, 1013–1020 (2013).

[9] Ahrens, M. B., Orger, M. B., Robson, D. N., Li, J. M. & Keller, P. J. Whole-brain functional imaging at cellular resolution using light-sheet microscopy. Nature methods 10, 413–420 (2013).

[10] Lemon, W. C. et al. Whole-central nervous system functional imaging in larval drosophila. Nature communications 6, 7924 (2015).

[11] Jun, J. J. et al. Fully integrated silicon probes for highdensity recording of neural activity. Nature 551, 232–236 (2017).

[12] Li, H. et al. Fly cell atlas: A single-nucleus transcriptomic atlas of the adult fruit fly. Science 375, eabk2432 (2022).

[13] Yao, Z. et al. A high-resolution transcriptomic and spatial atlas of cell types in the whole mouse brain. Nature 624, 317–332 (2023).

[14] Kim, J. S. et al. Space–time wiring specificity supports direction selectivity in the retina. Nature 509, 331–336 (2014).

[15] Wanner, A. A. & Friedrich, R. W. Whitening of odor representations by the wiring diagram of the olfactory bulb. Nature neuroscience 23, 433–442 (2020).

[16] Lyu, C., Abbott, L. & Maimon, G. Building an allocentric travelling direction signal via vector computation. Nature 601, 92–97 (2022).

[17] Vishwanathan, A. et al. Predicting modular functions and neural coding of behavior from a synaptic wiring diagram. Nature Neuroscience 27, 2443–2454 (2024).

[18] Lappalainen, J. K. et al. Connectome-constrained networks predict neural activity across the fly visual system. Nature 1–9 (2024).

[19] Taube, J. S. The head direction signal: origins and sensory-motor integration. Annual Review of Neuro-science 30, 181–207 (2007).

[20] Hulse, B. K. & Jayaraman, V. Mechanisms underlying the neural computation of head direction. Annual Review of Neuroscience 43, 31–54 (2020).

[21] Rank, J. Head-direction cells in the deep layers of dorsal presubiculum of freely moving rats. In Soc. Neuroscience Abstr., vol. 10, 599 (1984).

[22] Taube, J., Muller, R. & Ranck, J. Head-direction cells recorded from the postsubiculum in freely moving rats. description and quantitative analysis. Journal of Neuroscience 10, 420–435 (1990)…

[23] Chaudhuri, R., Gerçek, B., Pandey, B., Peyrache, A. & Fiete, I. The intrinsic attractor manifold and population dynamics of a canonical cognitive circuit across waking and sleep. Nature Neuroscience 22, 1512–1520 (2019). .

[24] Khona, M. & Fiete, I. R. Attractor and integrator networks in the brain. Nature Reviews Neuroscience 23, 744–766 (2022).

[25] Chaudhuri, R. & Fiete, I. Computational principles of memory. Nature Neuroscience 19, 394–403 (2016).

[26] Brody, C. D., Romo, R. & Kepecs, A. Basic mechanisms for graded persistent activity: discrete attractors, continuous attractors, and dynamic representations. Current Opinion in Neurobiology 13, 204–211 (2003). .

[27] Knierim, J. J. & Zhang, K. Attractor dynamics of spatially correlated neural activity in the limbic system. Annual Review of Neuroscience 35, 267–285 (2012). . PMID: 22462545, .

[28] Skaggs, W., Knierim, J., Kudrimoti, H. & McNaughton, B. A model of the neural basis of the rat’s sense of direction. Advances in Neural Information Processing Systems 7, 173–180 (1995).

[29] Amari, S.-i. Dynamics of pattern formation in lateralinhibition type neural fields. Biological Cybernetics 27, 77–87 (1977).

[30] Ben-Yishai, R., Bar-Or, R. L. & Sompolinsky, H. Theory of orientation tuning in visual cortex. Proceedings of the National Academy of Sciences 92, 3844–3848 (1995)…

[31] Zhang, K. Representation of spatial orientation by the intrinsic dynamics of the head-direction cell ensemble: a theory. Journal of Neuroscience 16, 2112–2126 (1996). .

[32] Redish, D., Elga, A. & Touretzky, D. A coupled attractor model of the rodent head direction system. Network: Computation in Neural Systems 7, 671–685 (1996).

[33] Camperi, M. & Wang, X.-J. A model of visuospatial working memory in prefrontal cortex: recurrent network and cellular bistability. Journal of Computational Neuroscience 5, 383–405 (1998).

[34] Compte, A., Brunel, N., Goldman-Rakic, P. S. & Wang, X.-J. Synaptic mechanisms and network dynamics underlying spatial working memory in a cortical network model. Cerebral Cortex 10, 910–923 (2000).

[35] Sharp, P. E., Blair, H. T. & Cho, J. The anatomical and computational basis of the rat head-direction cell signal. Trends in Neurosciences 24, 289–294 (2001).

[36] Kakaria, K. S. & de Bivort, B. L. Ring attractor dynamics emerge from a spiking model of the entire protocerebral bridge. Frontiers in Behavioral Neuroscience 11 (2017). .

[37] Darshan, R. & Rivkind, A. Learning to represent continuous variables in heterogeneous neural networks. Cell Reports 39, 110612 (2022). .

[38] Kim, S. S., Rouault, H., Druckmann, S. & Jayaraman, V. Ring attractor dynamics in the drosophila central brain. Science 356, 849–853 (2017).

[39] Noorman, M., Hulse, B. K., Jayaraman, V., Romani, S. & Hermundstad, A. M. Maintaining and updating accurate internal representations of continuous variables with a handful of neurons. Nature Neuroscience 1–11 (2024).

[40] Scheffer, L. K. et al. A connectome and analysis of the adult drosophila central brain. elife 9, e57443 (2020).

[41] Berg, S. et al. Sexual dimorphism in the complete connectome of the drosophila male central nervous system. bioRxiv 2025–10 (2025).

[42] Seelig, J. D. & Jayaraman, V. Neural dynamics for landmark orientation and angular path integration. Nature 521, 186–191 (2015).

[43] Green, J. et al. A neural circuit architecture for angular integration in drosophila. Nature 546, 101–106 (2017).

[44] Turner-Evans, D. B. et al. Angular velocity integration in a fly heading circuit. eLife 6, e23496 (2017).

[45] Turner-Evans, D. B. et al. The neuroanatomical ultrastructure and function of a biological ring attractor. Neuron 108, 145–163 (2020).

[46] Hulse, B. K. et al. A connectome of the drosophila central complex reveals network motifs suitable for flexible navigation and context-dependent action selection. Elife 10 (2021).

[47] Zheng, Z. et al. A complete electron microscopy volume of the brain of adult drosophila melanogaster. Cell 174, 730–743 (2018).

[48] Marin, E. C. et al. Systematic annotation of a complete adult male drosophila nerve cord connectome reveals principles of functional organisation. eLife 13 (2024).

[49] Bates, A. S. et al. Distributed control circuits across a brain-and-cord connectome. bioRxiv (2025).

[50] Renart, A., Song, P. & Wang, X.-J. Robust spatial working memory through homeostatic synaptic scaling in heterogeneous cortical networks. Neuron 38, 473–485 (2003).

[51] Braitenberg, V. & Schüz, A. Anatomy of the cortex: statistics and geometry, vol. 18 (Springer Science & Business Media, 2013).

[52] Darshan, R. & Rivkind, A. Learning to represent continuous variables in heterogeneous neural networks. Cell Reports 39 (2022).

[53] Ságodi, Á., Martín-Sánchez, G., Sokol, P. & Park, M. Back to the continuous attractor. Advances in Neural Information Processing Systems 37, 66856–66906 (2024).

[54] Duan, S., Dong, L. L. & Fiete, I. From synapses to dynamics: Obtaining function from structure in a connectome constrained model of the head direction circuit. bioRxiv 2025–05 (2025).

[55] Clark, D. G., Abbott, L. & Sompolinsky, H. Symmetries and continuous attractors in disordered neural circuits. bioRxiv 2025–01 (2025).

[56] Sun, Y. et al. Neural signatures of dynamic stimulus selection in drosophila. Nature neuroscience 20, 1104–1113 (2017).

[57] Kim, S. S., Hermundstad, A. M., Romani, S., Abbott, L. F. & Jayaraman, V. Generation of stable heading representations in diverse visual scenes. Nature 576, 126–131 (2019).

[58] Fisher, Y. E., Lu, J., D’Alessandro, I. & Wilson, R. I. Sensorimotor experience remaps visual input to a heading-direction network. Nature 576, 121–125 (2019).

[59] Haberkern, H. et al. Maintaining a stable head direction representation in naturalistic visual environments. BioRxiv 2022–05 (2022).

[60] Lu, J. et al. Transforming representations of movement from body-to world-centric space. Nature 601, 98–104 (2022).

[61] Dan, C., Hulse, B. K., Kappagantula, R., Jayaraman, V. & Hermundstad, A. M. A neural circuit architecture for rapid learning in goal-directed navigation. Neuron 112, 2581–2599 (2024).

[62] Pisokas, I., Heinze, S. & Webb, B. The head direction circuit of two insect species. Elife 9, e53985 (2020).

[63] Stone, T. et al. An anatomically constrained model for path integration in the bee brain. Current Biology 27, 3069–3085 (2017).

[64] Touretzky, D. S., Redish, A. D. & Wan, H. S. Neural representation of space using sinusoidal arrays. Neural Computation 5, 869–884 (1993).

[65] Wittmann, T. & Schwegler, H. Path integration—a network model. Biological Cybernetics 73, 569–575 (1995).

[66] Varga, A. G. & Ritzmann, R. E. Cellular basis of head direction and contextual cues in the insect brain. Current Biology 26, 1816–1828 (2016).

[67] Petrucco, L. et al. Neural dynamics and architecture of the heading direction circuit in zebrafish. Nature Neuroscience 26, 765–773 (2023). .

[68] Wu, Y. K., Petrucco, L. & Portugues, R. Anatomical and functional organization of the interpeduncular nucleus in larval zebrafish. bioRxiv 2024–10 (2024).

[69] Tanaka, R. & Portugues, R. Mechanisms for plastic landmark anchoring in zebrafish compass neurons. bioRxiv 2024–12 (2024).

[70] Gardner, R. J. et al. Toroidal topology of population activity in grid cells. Nature 602, 123–128 (2022).

[71] Perea, G., Sur, M. & Araque, A. Neuron-glia networks: integral gear of brain function. Frontiers in cellular neuroscience 8, 378 (2014).

[72] Mountoufaris, G. et al. A line attractor encoding a persistent internal state requires neuropeptide signaling. Cell 187, 5998–6015 (2024).

[73] Turrigiano, G. G. Homeostatic plasticity in neuronal networks: the more things change, the more they stay the same. Trends in neurosciences 22, 221–227 (1999).

[74] Marder, E. From biophysics to models of network function. Annual review of neuroscience 21, 25–45 (1998).

[75] Dana, H. et al. High-performance calcium sensors for imaging activity in neuronal populations and microcompartments. Nature methods 16, 649–657 (2019).

[76] Dionne, H., Hibbard, K. L., Cavallaro, A., Kao, J.-C. & Rubin, G. M. Genetic reagents for making split-gal4 lines in drosophila. Genetics 209, 31–35 (2018).

[77] Seelig, J. D. et al. Two-photon calcium imaging from head-fixed drosophila during optomotor walking behavior. Nature methods 7, 535–540 (2010).

[78] Pologruto, T. A., Sabatini, B. L. & Svoboda, K. Scanimage: flexible software for operating laser scanning microscopes. Biomedical engineering online 2, 1–9 (2003).

[79] Berens, P. Circstat: a matlab toolbox for circular statistics. Journal of statistical software 31, 1–21 (2009).

[80] Hastie, T. Tibshirani, T. Friedman, J. H. & Friedman, J. H. The elements of statistical learning: data mining, inference, and prediction, vol. 2 (Springer, 2009).

## References

[1] Lappalainen, J. K. et al. Connectome-constrained networks predict neural activity across the fly visual system. Nature 1–9 (2024).

[2] Noorman, M., Hulse, B. K., Jayaraman, V., Romani, S. & Hermundstad, A. M. Maintaining and updating accurate internal representations of continuous variables with a handful of neurons. Nature Neuroscience 1–11 (2024).

[3] Burak, Y. & Fiete, I. R. Fundamental limits on persistent activity in networks of noisy neurons. Proceedings of the National Academy of Sciences 109, 17645–17650 (2012).

[4] Turner-Evans, D. B. et al. The neuroanatomical ultrastructure and function of a biological ring attractor. Neuron 108, 145–163 (2020).

